# Targeted protein degradation reveals BET bromodomains as the cellular target of Hedgehog Pathway Inhibitor-1

**DOI:** 10.1101/2022.08.16.504103

**Authors:** Meropi Bagka, Hyeonyi Choi, Margaux Heritier, Leonardo Scapozza, Yibo Wu, Sascha Hoogendoorn

## Abstract

Target deconvolution of small molecule hits from phenotypic screens presents a major challenge. Illustrative of these are the many screens that have been conducted to find inhibitors for the Hedgehog (Hh) signaling pathway – a major developmental pathway with many implications in health and disease - with many hits but very few identified cellular targets. We here present a strategy for target identification based on Proteolysis-Targeting Chimeras (PROTACs), combined with label-free quantitative proteomics. We developed a PROTAC based on the downstream Hedgehog Pathway Inhibitor-1 (HPI-1), a phenotypic screen hit with unknown cellular target. Using our Hedgehog Pathway PROTAC (HPP) we identified and validated BET bromodomains to be the cellular targets of HPI-1. Furthermore, we found that HPP-9 has a unique mechanism of action as a long-acting Hh pathway inhibitor through prolonged BET bromodomain degradation. Collectively, we provide a powerful PROTAC-based approach for target deconvolution, that has answered the longstanding question of the cellular target of HPI-1 and yielded the first PROTAC that acts on the Hh pathway.

## Main

The Hedgehog pathway is a complex cellular signaling cascade that regulates embryonic developmental processes, such as patterning, as well as stem cell maintenance and tissue homeostasis.^1,2^ Dysregulation of physiological levels of Hedgehog signal transduction results in developmental disorders and the onset and progression of various cancers, most notably basal cell carcinoma and medulloblastoma.^3,4^ Pathway activation under normal conditions is initiated by the binding of one of the Hedgehog proteins (IHH, DHH, SHH) to the receptor Patched (PTCH1).^5–7^ The binding of HH to PTCH1 releases the latter’s inhibitory effect on Smoothened (SMO).^8,9^ Further activation steps include the trafficking of GLI2/3 transcription factors bound to Suppressor of Fused (SUFU) through, and accumulation at the tip of, the primary cilium.^10–13^ Processing of the GLI transcription factors into their transcriptionally active form then results in the transcription of Hedgehog target genes, amongst which the positive regulator *Gli1* and, in a negative feedback loop, *Ptch1*.^14,15^ At present, the only clinically approved drugs to combat Hh pathway-driven cancers are those directed at SMO (vismodegib, sonidegib). Cancers driven by downstream pathway activation are inherently insensitive to those drugs and acquired resistance of initially responsive tumors is common.^16–20^ Strategies to inhibit the Hedgehog pathway beyond Smoothened are scarce, with only a handful of reported molecules with well-defined cellular targets and mechanism-of-action, including the ciliobrevins^21^, arsenic trioxide^22^, physalin H^23^, and Glabrescione B.^24^ The majority of reported molecules have been identified through phenotypic screens for the Hedgehog pathway, and it has proven highly challenging to unravel the cellular target and molecular mechanisms of these hit compounds, severely limiting their use as chemical probes or therapeutic leads.^25–32^

Exemplary of this is Hedgehog Pathway Inhibitor (HPI-1), a dihydropyridine molecule discovered by Hyman et al. as a robust downstream Hh signaling inhibitor, with anti-cancer properties.^26,33,34^ Often referred to as a GLI inhibitor,^35–37^ its cellular target has remained elusive for many years. Here, we present a target identification methodology based on targeted protein degradation coupled to label-free quantitative proteomics. Using a Hedgehog Pathway Proteolysis-Targeting Chimera (PROTAC) (HPP), a bifunctional molecule consisting of HPI-1 coupled to a CRBN ligand, we elucidated BET bromodomains as the cellular targets of HPI-1. Moreover, we show that degradation of BET bromodomains through HPP-9 results in extended modulation of the Hedgehog pathway, enabling novel pharmacological strategies for this important developmental pathway.

## Results

### Design and synthesis of Hedgehog Pathway PROTACS

Proteolysis targeting chimeras (PROTACs) are bifunctional molecules that induce proteasome-mediated degradation of a protein of interest (POI) through the formation of a ternary complex between the POI and an E3 ligase.^38,39^ PROTACs have found widespread use as target validation tools, and as promising therapeutic leads because of their unique mechanism of action.

We hypothesized that we could extend the use of PROTACs as a target deconvolution method for hits from phenotypic screens through quantitative comparative proteomics. For this, we synthesized a small library of PROTACs, based on Hedgehog Pathway Inhibitor 1 (HPI-1, Fig. 1a). Building on and expanding existing structure-activity relationship (SAR) studies^40^, we found the phenolic hydroxyl to be the optimal position to introduce functionality, as its modification did not change the overall inhibitory potency of the molecule and it can be modified using late-stage functionalization (Extended Fig. 1a). In order to turn HPI-1 into a bifunctional degrader molecule, we then explored various linkers (polyethylene glycol of different length, simple aliphatic chains), attachment chemistries (triazole, ether, amide), and E3 ligase targeting ligands (VHL peptide ligand^41^, Pomalidomide^42^, hydroxythalidomide^43^) and assessed the inhibitory potencies of the resulting Hedgehog Pathway PROTACs (HPP-1 to HPP-11) in a variety of Hedgehog pathway activity assays (Fig. 1). SHH-LIGHT2 cells (NIH-3T3 cells stably expressing a GLI-driven luciferase reporter^44^) were stimulated with either Sonic Hedgehog-containing medium (ShhN) or the small molecule Smoothened agonist (SAG)^45^ in the presence of 10 µM of HPP (Fig. 1c). We found that HPPs incorporating a CRBN ligand were more potent than the VHL peptide analogs in inhibiting pathway activation. Furthermore, HPPs with short aliphatic linkers performed better than those with PEG linkers.

**Fig. 1.**
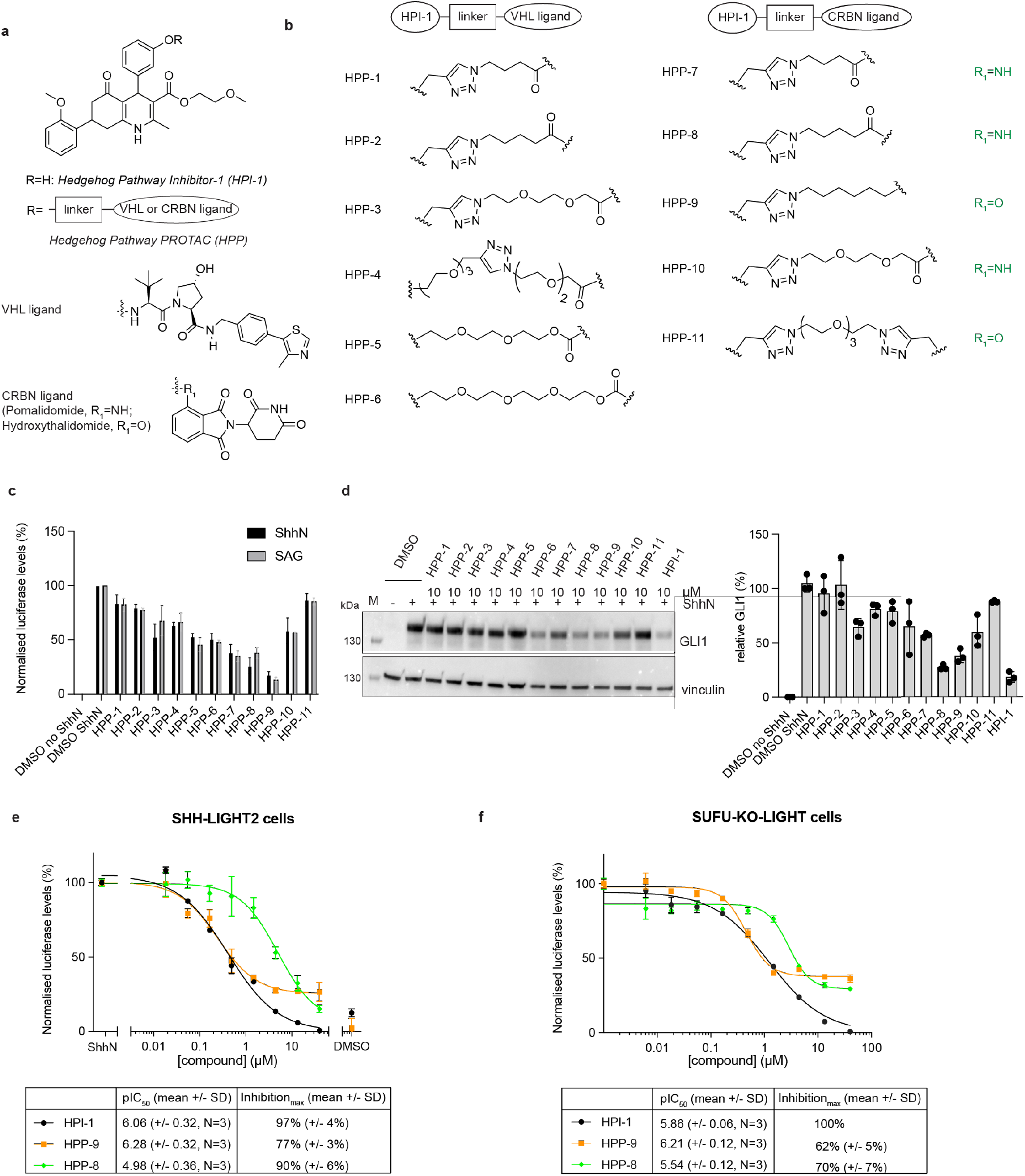
Design, synthesis and biological evaluation of Hedgehog Pathway PROTACs. a) Structure of HPI-1, VHL ligand and CRBN ligand and schematic design of the PROTACs. b) HPP-1 to HPP-11 varied in the linker used to connect the E3 ligase ligand (VHL or CRBN ligand) to HPI-1. The HPPs were evaluated c) in a luciferase assay in SHH-LIGHT2 cells stimulated with ShhN or SAG and d) by western blot for GLI1 levels for their inhibitory potential at 10 µM.

To exclude direct interference of the compounds with the luciferase enzymatic activity, the results were confirmed by western blot analysis for GLI1 protein expression (Fig. 1d). There was a good correlation between both readouts (extended Fig. 1b) and subsequently, full dose-response curves were generated for the HPPs that showed the lowest residual activity at 10 µM (HPP-8 and HPP-9). In contrast to the full Hh pathway inhibition that was obtained for the parent molecule HPI-1, inhibition by the PROTACs plateaued at 10-20% in SHH-LIGHT2 cells (Fig. 1e) and at 30-40% in constitutively activated SUFU-KO-LIGHT cells^35^ (Fig. 1f), which could point to a differential mode of action of the HPPs compared to the parent molecule. HPP-9 showed comparable potency to HPI-1 in both cell lines and was selected as the most suitable candidate for further studies.

Mean +/- SEM of at least 3 independent experiments is plotted. e,f) Full dose response curves were generated for most potent analogs HPP-8 and HPP-9 in e) SHH-LIGHT2 cells stimulated with ShhN or f) constitutively active SUFU-KO-LIGHT cells. Representative curves of three independent experiments are shown.

**Extended Fig. 1.**
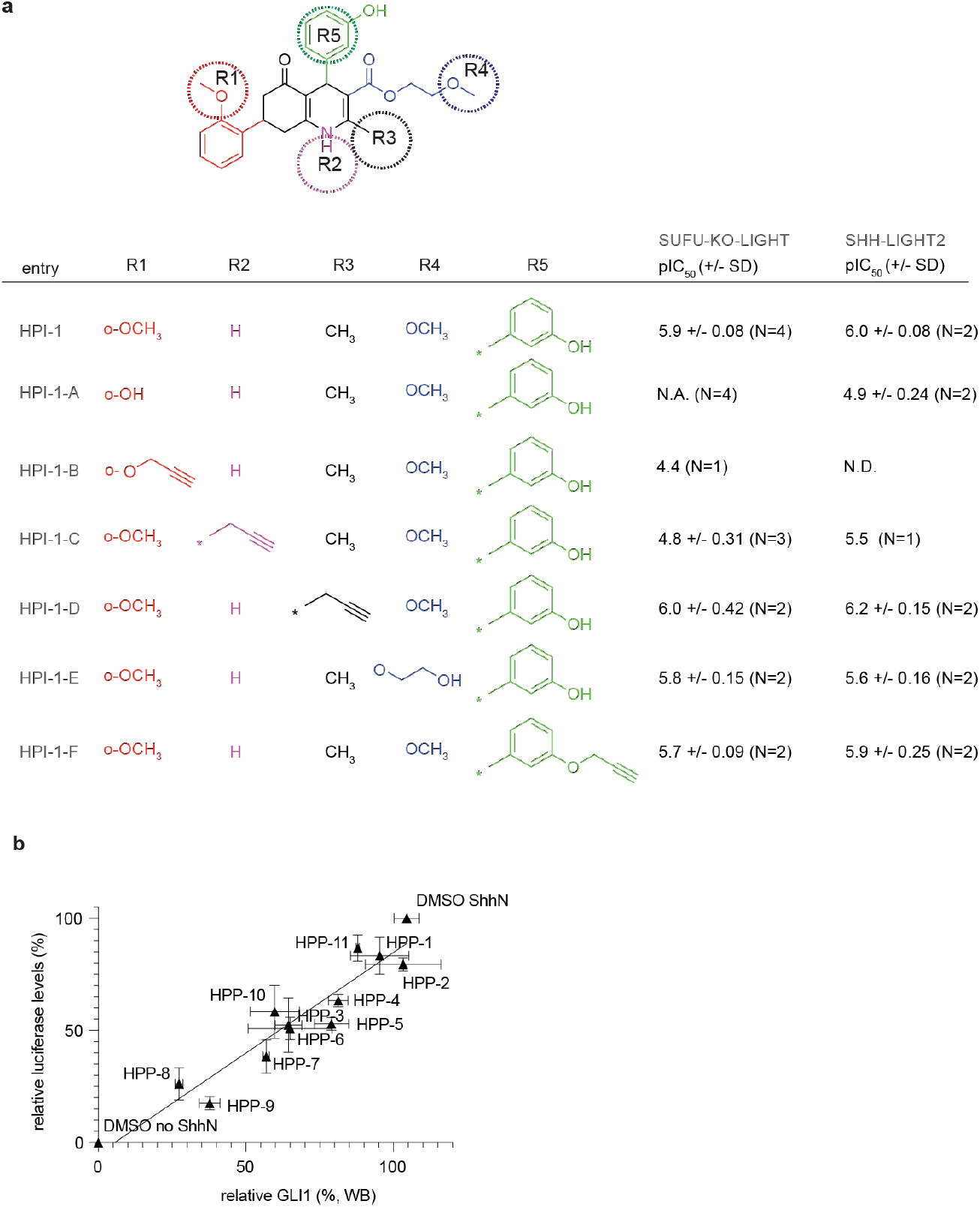
Design, synthesis and biological evaluation of Hedgehog Pathway PROTACs. a) SAR analysis for HPI-1 analogs. b) Readout by luciferase assay and GLI1 western blot for the HPP-9 show good correlation (related to data shown in Fig. 1c, d)

### Methylation of thalidomide blocks HPP-9 activity

As the target of HPI-1 is unknown, the mode of action of HPP-9 as a PROTAC could not be directly proven and its inhibitory effect on the Hedgehog pathway could be the result of direct inhibition and/or degradation of the HPI-1 target. We sought to differentiate between these options by implementing a small structural change to the thalidomide core (methylation) such that the molecule could no longer bind to CRBN (Fig. 2a)^46^. We developed a high throughput microscopy assay using NIH-3T3 cells containing an Hh pathway-driven GFP reporter (SHH-GFP cells)^47^, to assess the inhibitory potency of this degradation-deficient HPP-9 analog (inact-HPP-9) (Fig. 2b,c). We found that inact-HPP-9 was indeed a very poor inhibitor compared to HPI-1 or HPP-9. Furthermore, in this assay we observed a bell-shaped curve for HPP-9, which could be indicative of the presence of a Hook effect for this compound. When cells were incubated with various concentrations of HPP-9 or HPI-1 and analyzed by western blot for GLI protein expression or processing (Fig. 2d), we observed a similar trend. As previously reported for HPI-1^26^, no effect on GLI3 processing was found for either compound, indicating that these compounds act downstream of GLI3 activation at the level of the primary cilium. Whereas HPI-1 inhibited ShhN-induced GLI1 expression and reduced the GLI2 full-length/activator levels in a dose-dependent fashion (Fig. 2d,e), the effect of HPP-9 showed again an inverse correlation where 10 µM of compound was less efficient than 0.5 or 2 µM. We then assessed the induction of Hh pathway target genes (*Gli1, Ptch1*) by qPCR (Fig. 2f,g), and in agreement with what was found on the protein level by western blot, we found a significant reduction in transcript levels.^15^ As GLI2 protein levels were reduced below basal level for HPP-9 and the highest concentration of HPI-1, we performed a qPCR for *Gli2* (Fig. 2h) and found levels to be much lower in both the minus and plus ShhN conditions.^15^ Interestingly, the fold induction upon the addition of ShhN remained constant at slightly over 2-fold under all conditions. Taken together, these results strongly suggested that HPP-9 acts as a PROTAC and is thus a promising lead for our proteomic target identification studies.

**Fig. 2.**
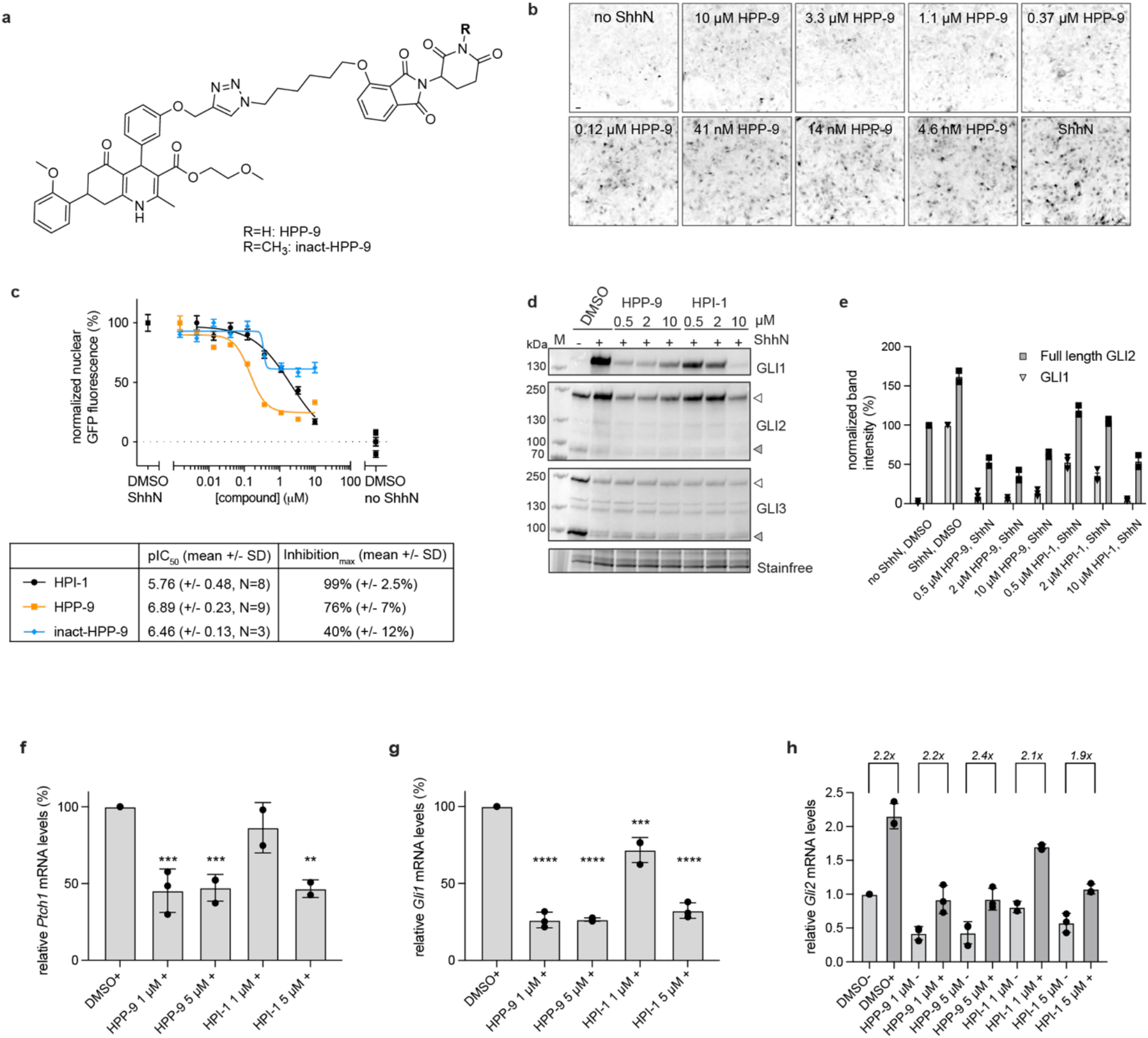
Biological evaluation of HPP-9 and its inactive analog. a) Chemical structures of HPP-9 and the methylated analog inact-HPP-9. b) Representative micrographs showing the dose-dependent inhibition of a Shh-driven GFP reporter by HPP-9, resulting in the dose-response curves shown in c). Scalebar 30 µm. c) Curves are representative of N independent experiments, with 9-18 images analyzed per experiment. d,e) NIH-3T3 cells were incubated with increasing concentrations of HPP-9 or HPI-1 in the presence of ShhN and lysates were probed for GLI1, GLI2 and GLI3. D) shows a representative immunoblot illustrating that both compounds inhibit GLI1 and GLI2 but have no effect on GLI3 processing. e) Quantification of GLI1 and GLI2 full length levels of 3 independent experiments. f-h) qPCR for f) *Ptch1*, g) *Gli1*, and h) *Gli2* shows that HPP-9 reduces the expression of Hh pathway target genes, while also decreasing basal *Gli2* transcript levels without affecting the fold-induction upon pathway stimulation (-: no ShhN, + ShhN). Data shown is the mean +/- SD for N=2-3 independent experiments. f, g) Student’s t-test, **** p<0.0001, *** p<0.001, ** p>0.01, compared to DMSO ShhN.

### Label free quantitative proteomics reveals BET Bromodomains as putative HPP-9 targets

We hypothesized that we could identify the target of HPI-1 through targeted protein degradation using HPP-9. For this, we applied label free quantitative proteomics on cells treated with DMSO, 1 µM HPI-1 or 1 µM HPP-9 for 27h. We applied data-independent acquisition mass spectrometry (DIA-MS) and quantified 5343 proteins across 12 samples. Expression of the 5343 proteins clearly separated the three sample groups under unsupervised clustering, with the HPI-1 group being intermediate between the DMSO and the HPP-9 group, suggesting distinguishable proteome changes in response to different treatments (Figure 3a, supplementary table 1). Among the 5343 quantified proteins, 246 proteins were significantly regulated by HPI-1 treatment (fold change > 1.5, Q value < 0.05), 468 proteins were significantly differentially expressed comparing HPP-9- and DMSO-treated groups, and 140 proteins showed differential expression upon both treatments (supplementary table 2). Interestingly, among the 468 significant proteins induced by HPP-9 treatment, 437 (93.4%) proteins were down-regulated in response to the treatment, indicating potent protein degradation induced by HPP-9. We then sought to discover proteins that were exclusively downregulated by the PROTAC, HPP-9, but not HPI-1.

For our analysis we focused on proteins that were significantly downregulated in our HPP-9 samples compared to HPI-1 for the following reasons. First, we expected that HPI-1 treatment results in changes in the proteome that would be mirrored by HPP-9, as they have shared biological activity. Second, as HPP-9 is designed to act as a PROTAC, it should lead to the degradation of its protein targets, whereas it is not expected that HPI-1 would do the same. We thus focused on the 64 proteins that were selectively downregulated in the HPP-9 treated samples (log2(fold change) >-1, -log 10 (Q value) > 2.5) (Figure 3b,c). Gene ontology (GO)-term analysis with the Database for Annotation, visualization and Integrated Discovery (DAVID) on the 64 unique HPP-9 hits, revealed ‘lysine acetylated histone binding’ (P= 1.6 × 10^−3^), ubiquitin protein ligase binding (P= 8.2 × 10^−3^), and ‘chromatin remodeling’ (P= 1 × 10^−2^). The significant depletion of two members of Bromodomain and Extra-Terminal Domain (BET) protein family (BRD3 and BRD4) exclusively in the HPP-9 treated samples caught our attention. The role of the BET bromodomains (BRD2/3/4 and testis-specific BRDT) and well-known inhibitors thereof (JQ1^48,49^, iBET-151^50^) in epigenetic modulation of the Hedgehog pathway is well-established. All available data on HPI-1 as a downstream inhibitor point towards it regulating GLI transcriptional activity, making BET bromodomains likely relevant targets. A major advantage of our approach is that we could directly test this hypothesis by using our PROTAC HPP-9 and evaluating BRD protein degradation.

**Fig. 3.**
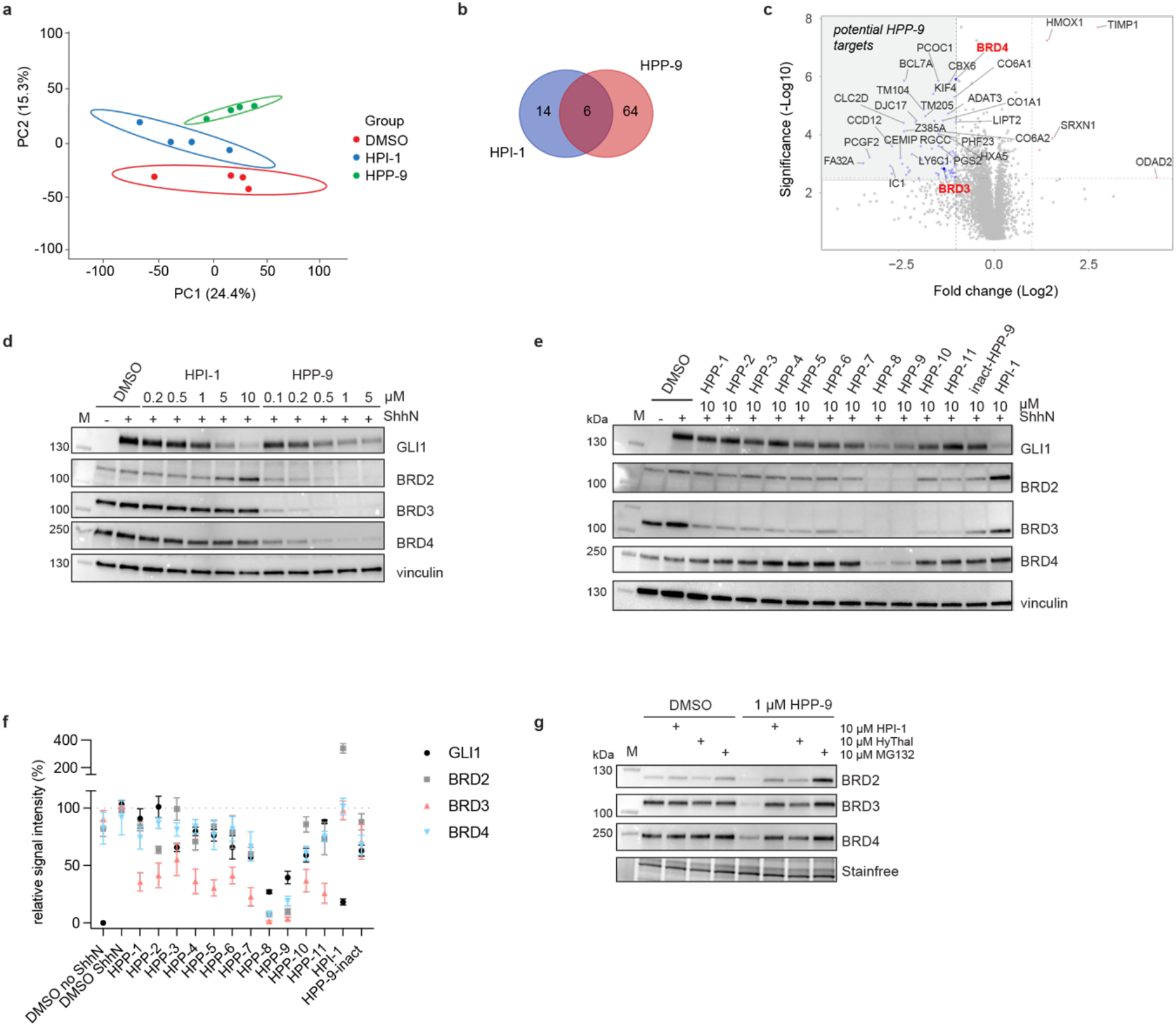
HPP-9 acts as a PROTAC for BET bromodomains. a) Principal component analysis of 5343 quantified proteins clearly separates all sample groups. b) A Venn diagram showing the overlap in most signifcantly downregulated proteomic hits between HPI-1 and HPP-9-treated cells. c) Volcano plot of log2 fold change to significance shows the potential direct targets of HPP-9. The top 25 hits are labeled, as well as BRD3. d) HPP-9 dose-dependently degrades the BET bromodomain proteins as shown by western blot analysis. Representative blot of 3 independent experiments. e, f) All HPPs were profiled for their ability to degrade BRD2/3/4 by western blot which revealed f) a good correlation between Hedgehog pathway inhibition and BET bromodomain degradation. Mean of 3 independent experiments +/- SEM is plotted. g) Representative immunoblot of a competition experiment between HPP-9 and HPI-1, hydroxythalidomide and MG-132. Three independent experiments.

### HPP-9 is a potent BET bromodomain degrader that acts through a PROTAC mechanism

To assess the capability of HPP-9 to degrade BET bromodomains, NIH-3T3 cells were incubated with increasing concentrations of HPP-9 for 27h in the presence of ShhN and probed for GLI1 (as a Hedgehog pathway readout), BRD2, BRD3, and BRD4. As shown in Fig. 3d, all three BRDs were almost completely degraded by our PROTAC at concentrations between 0.5 and 5 µM. In contrast, the probe was less effective at 10 µM, mirroring the Hook effect we observed before in our Hedgehog pathway assays (Extended Fig. 3a). We then profiled our original library of HPPs for their ability to degrade BRD2/3/4 (Fig. 3e) and found a good correlation between the degradation efficiency and extend of Hedgehog pathway inhibition for all PROTACs (Fig. 3f). As expected, the inact-HPP-9 was completely incapable of degrading the BET bromodomains.

To prove that HPP-9 acts through a typical PROTAC mechanism, we performed competition assays with an access of the proteasome inhibitor MG132 (to prevent proteasomal degradation), the CRBN ligand hydroxythalidomide (to block all available CRBN E3 ligase), and the parent HPI-1 (to compete for binding of HPP-9 to BRDs) (Fig. 3g). In all cases, degradation was completely prevented, confirming that HPP-9 degrades BRDs through the formation of a ternary complex between BET bromodomains and CRBN, and subsequent proteasomal degradation.

**Extended Fig. 3.**
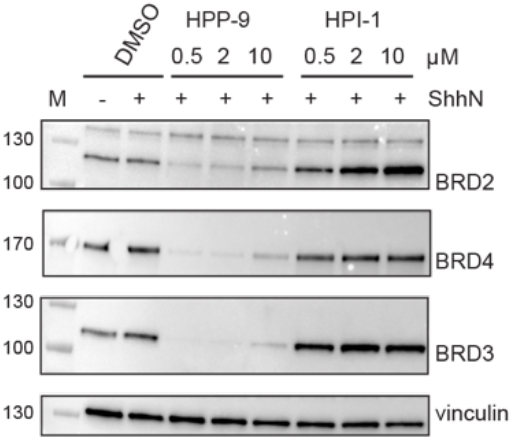
SHH-GFP cells were treated with various concentrations of HPP-9 or HPI-1 for 27h and probed for BRD2, BRD3 and BRD4 by WB. Representative immunoblot of two independent experiments. Related to Fig. 3d.

### HPP-9 and dBet6 have a differential mechanism of action

Various PROTACs targeting the BET bromodomains have been reported, including dBet6^51^, a JQ1-based pan-BET bromodomain PROTAC. We profiled our HPP-9 against dBet6 and surprisingly, we found that these compounds behaved differently (Fig. 4a-d). First, the partial inhibition of Hh signaling by HPP-9 was not found for dBet6, as it inhibited the Hh pathway slightly more potently (pIC^50^ 7.76 for dBet6 vs 6.71 for HPP-9) and fully. However, in contrast to previous reports, no significant degradation of BRD2 was observed. Of note, these experiments were done under Hedgehog signaling conditions (1 µM of compound in the presence of ShhN for 27h), to simultaneously assess the ability of the compounds to suppress the Hh pathway dependent GFP reporter (Fig. 4b-d) or endogenous GLI1 expression (Fig. 4a) as well as bromodomain degradation (Fig. 4a-d). In contrast, when we performed a time-course assay to assess how fast HPP-9 degrades the BRDs (Fig. 4e), we found that dBet6 is certainly an effective pan-bromodomain degrader at shorter timepoints, with complete degradation as soon as 2.5 hours after addition to the cells. HPP-9-mediated degradation of BRD2/3/4 reaches plateau around 7h, and in contrast to dBet6, remains so at longer timepoints. As JQ1-based BET bromodomain inhibitors typically reduce cell viability in a dose-dependent fashion, we treated a variety of mouse (NIH-3T3, IMCD3) and human (HEK293, HeLa) cell lines with increasing concentrations of HPP-9 or dBet6 for 48 h and we found HPP-9 to be much less toxic than dBet6 (Extended Fig. 4a).

**Fig. 4.**
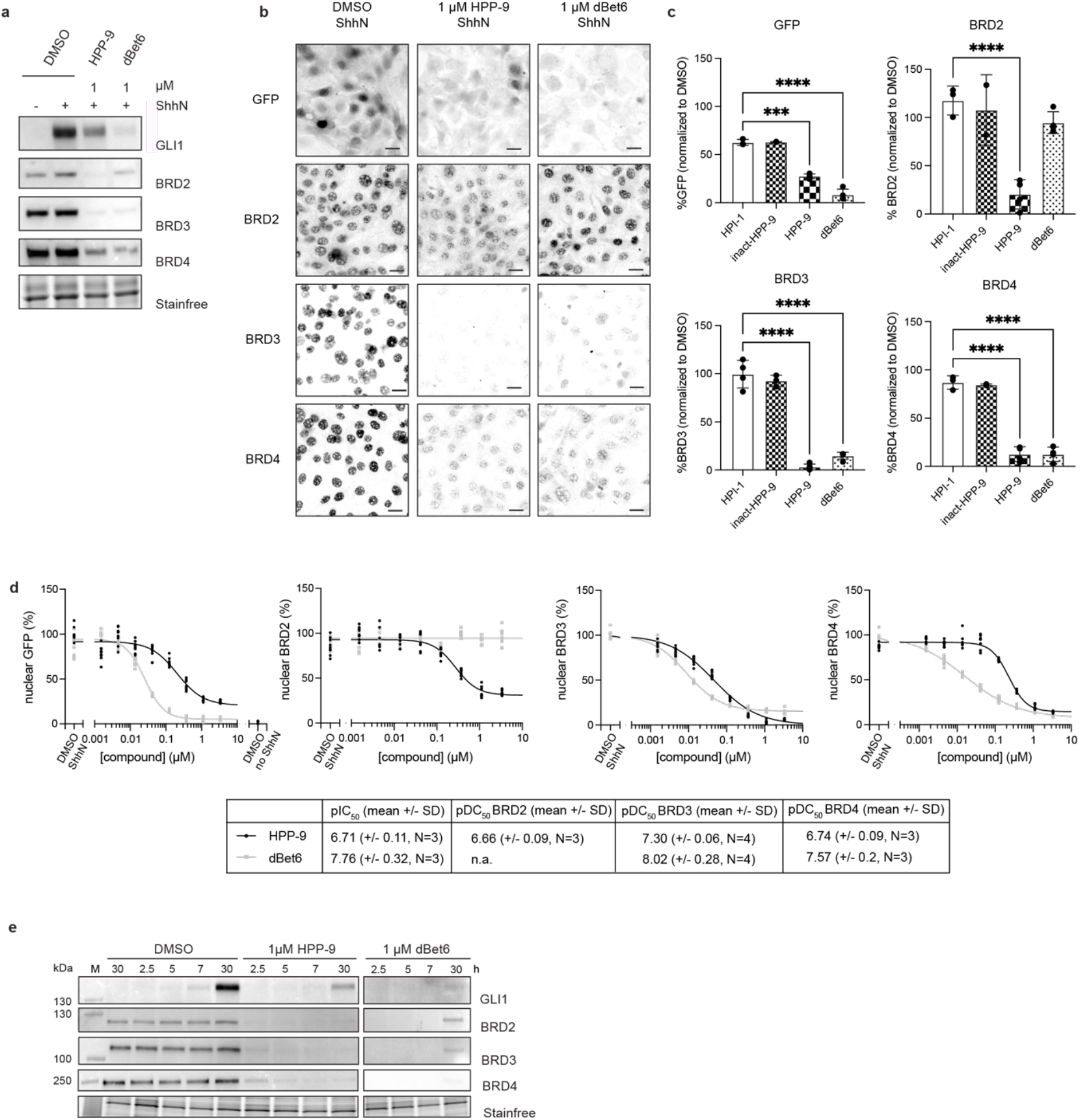
HPP-9 and dBet6 have a different mode of action. SHH-GFP cells were treated with 1 µM of HPP-9 or dBet6 for 27h and a) probed for GLI1, BRD2, BRD3 and BRD4 by WB and b,c) analyzed by fluorescence microscopy. Representative micrographs showing the nuclear localization of GFP, BRD2, BRD3 and BRD4 are shown in b) and the signal is quantified in c). Scalebar 25 µm. Data shown is from n=7-9 images analyzed per condition from N=2-4 independent experiments. D) Full dose-response curves were measured by high-content microscopy to determine the pDC_50_ values for HPP-9 and dBet6. Each dot is an individual image (n=7-9 images/experiment, N=3,4 independent experiments) E) SHH-GFP cells were treated with 1 µM of HPP-9 or dBet6 for the indicated times and lysates were probed for GLI1, BRD2, BRD3 and BRD4 by WB. Representative blot of two independent experiments is shown.

**Extended Fig. 4.**
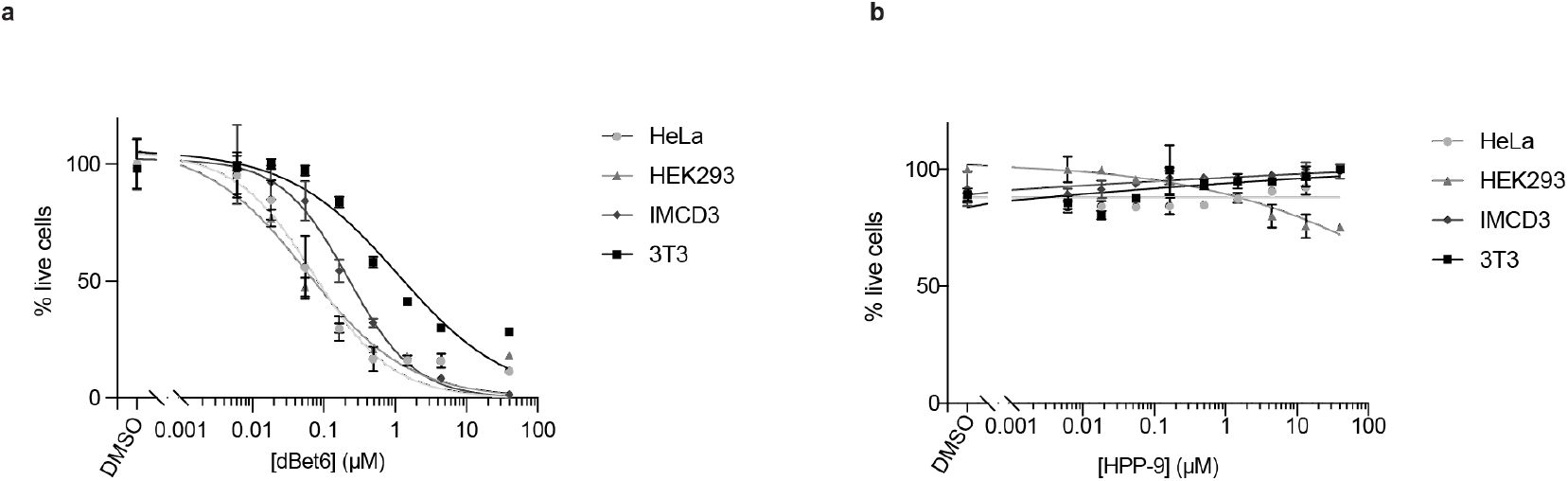
The indicated cell lines were incubated with varying concentrations of a) dBet6 or b) HPP-9 and cell viability was measured 48h later. Representative curves of 2 independent experiments are shown.

### HPI-1 is a high affinity BET bromodomain binder

Through our competition assays (Fig. 3f), we found that HPI-1 can prevent HPP-9-induced BRD2/3/4 degradation, which strongly indicates that they are acting on the same targets. As shown in Figure 3d-f, we observed a strong increase in BRD2 protein levels by western blot when we incubated cells with increasing concentrations of HPI-1 for 28h. This was validated by dose-response fluorescence microscopy to be around 2-fold more nuclear BRD2 at 20 µM concentration of HPI-1 (Fig. 5a,b). We then wanted to confirm if this increase in protein levels is a result of enhanced *Brd2* transcription as reported for JQ1 ^52^. Indeed, by qPCR we found *Brd2* to be highly expressed when treating the cells with 1 µM JQ1, and a much smaller but significant increase was found with 5 µM of HPI-1 (Fig. 5c). However, no change in BRD2 protein could be detected when cells were treated with 1 µM JQ1 (data not shown).

**Fig. 5.**
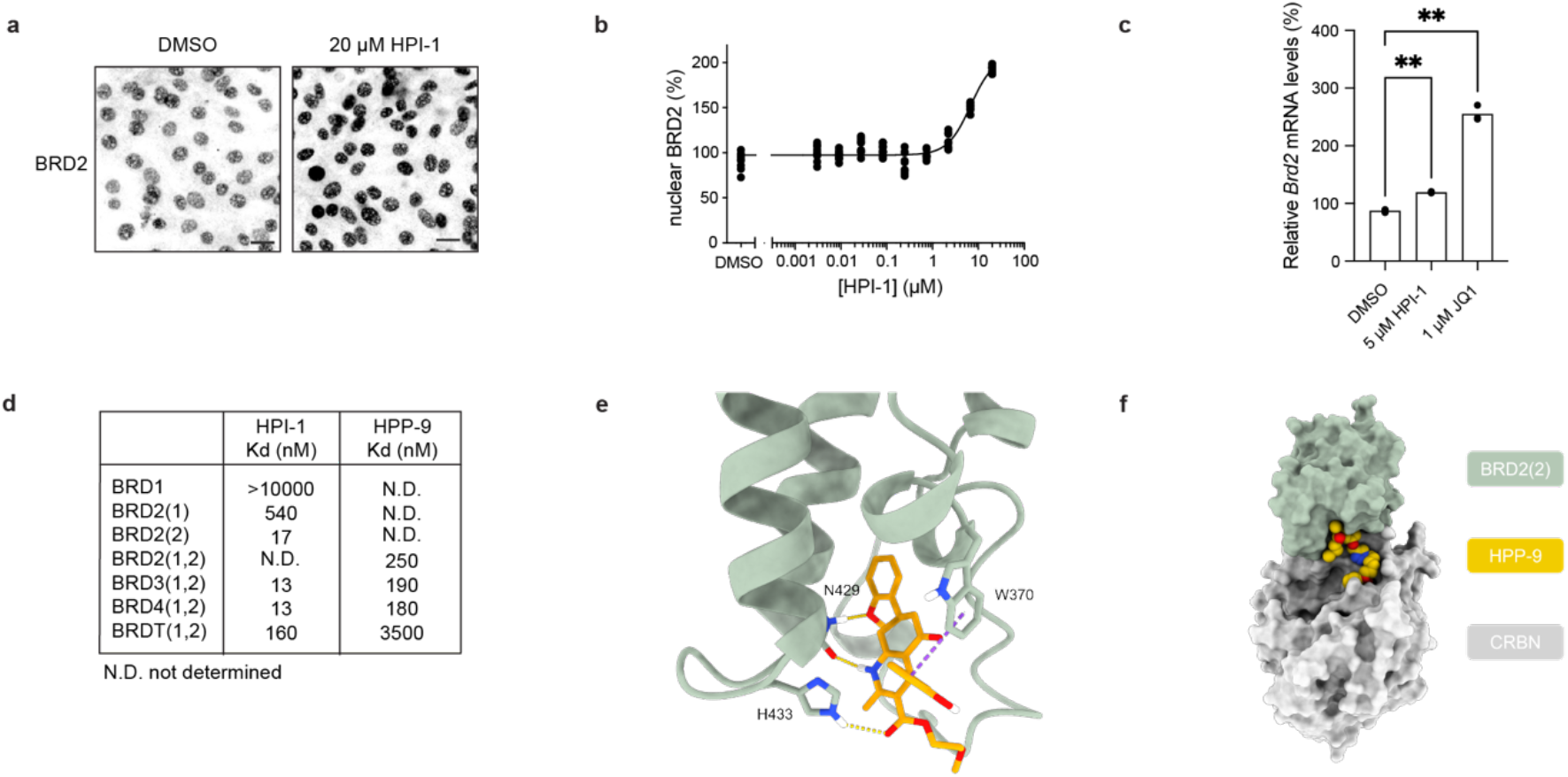
HPI-1 is a high-affinity BET bromodomain binder that increases cellular BRD2 levels. a) Representative micrographs of SHH-GFP cells treated with DMSO or 20 µM of HPI-1 for 27h. Scalebar 25 µm. b) High content microscopy dose-response curve for cells treated with increasing concentrations of HPI-1 and probed for nuclear BRD2. Representative curve from N=3 independent experiments, with n=9 images analyzed per condition. c) qPCR analysis of *Brd2* mRNA levels from cells treated with 5 µM HPI-1 or 1 µM JQ1 for 27h in the presence of ShhN. Data from N=2-3 independent experiments performed in duplicate. d) The affinity of HPI-1 and HPP-9 to the indicated bromodomains (BromoKd, Eurofins discovery). (1,2) indicates which of the two bromodomain subunits were included. e) Proposed binding mode for HPI-1. The interacting residue and HPI-1 are modelled in stick, while BRD2(2) is shown in cartoon. The dashed yellow lines represent H-bond while the π-π stacking is shown in purple dashed line. f) Possible physical model of the ternary complex. BRD2(2) (green) and CRBN (gray) are shown in a surface representation while HPP9 (orange) is shown in sphere.

To further prove the direct interaction between HPI-1/HPP-9 and BET bromodomain proteins, the compounds were sent for affinity measurements (BromoKd assay, Eurofins Discovery) against BRD2/3/4 and BRDT. For HPI-1, we included the non-BET bromodomain BRD1 as this protein was also found to be downregulated by HPI-1 in our proteomics dataset. Since the effect on BRD2 protein levels when treating cells with HPI-1 was so large, we determined the affinity for the two different bromodomains of BRD2, BD1 and BD2, separately. As shown in Figure 5d, HPI-1 is a high affinity bromodomain binder with low nM Kd for BRD2/3/4 and about 10-fold lower affinity for BRDT. HPI-1 showed much higher affinity to BD2 of BRD2 (17 nM), than to BD1 (540 nM), but otherwise no apparent selectivity between BRD2/3/4. No binding for BRD1 could be detected, indicating that the reduction in protein levels found by proteomics likely arose from an indirect effect. The affinity of HPP-9 was an order of magnitude lower than that of HPI-1 (∼200 nM versus ∼15 nM), which is in agreement with the low inhibitory potency of inact-HPP-9, thus confirming that HPP-9 inhibits mostly through degradation and not direct inhibition.

To get insights in a potential binding mode of HPI-1, we docked the compound in the bromodomain for which high affinity has been measured unequivocally, namely the second bromodomain of BRD2 (BRD2(2)) (Fig 5d). We used different crystal structures in both apo and holo form to explore the variety of conformations of the binding site, and validated the poses with the ligand-based SAR of the HPI-1 derivatives (extended Fig. 1) and the HPP series (Fig. 1). The most plausible binding mode considering both the information from the ligand-based SAR and the molecular docking results, suggests that HPI-1 forms strong interactions with the amino acids forming the binding site (Fig. 5e, extended Fig. 5). In particular, the conserved residue N429 forms the same network of H-bonds with HPI-1 as other inhibitors.^53–55^ HPI-1 interacts via a T-shaped π-π stacking with tryptophan 370 (W370) and one of its carbonyl groups binds to histidine 433 (H433) being the H-bond donor. This last interaction might explain the difference in affinity between the second and the first bromodomain of BRD2, where histidine is replaced by an aspartate, abrogating this predicted H-bond. The number and type of the interactions found within the obtained model (Fig. 5e, extended Fig. 5) are consistent with the high affinity found experimentally and recapitulate the ligand-based SAR extending it to 3D.

We next sought to assess *in silico* the possibility of forming a physical ternary complex between BRD2(2), HPP9 and CRBN starting with the proposed binding mode of HPI-1. We repurposed an existing protocol developed for evaluating different linkers in PROTAC design ^56^ to generated physical models. The protocol consists of three steps: the generation of BRD2(2)-CRBN complexes by protein-protein docking, the overlap of HPP9 -linker conformers on the resulting complexes, and finally a minimization of the ternary complex. The complex that displays the best interaction score (Fig. 5f) shows that the linker of HPP-9 is long enough to allow both moieties to bind their respective protein and form a low-energy structure (Fig. 5f).

Finally, HPI-1 has been deposited in the public repository of the NCI-60 as a putative SMO inhibitor. We employed the NIH COMPARE algorithm (https://nci60.cancer.gov/publiccompare/), to find compounds that share the highest overlap in cellular activity profiles. We found that HPI-1 overlapped most significantly (Pearson’s t-test) with other annotated BET bromodomain inhibitors, including INCB-057643 (R^2^>0.67), JQ1 (R^2^>0.64), and I-BET151 (R^2^>0.64), providing further evidence of a shared cellular target.

**Extended Fig. 5.**
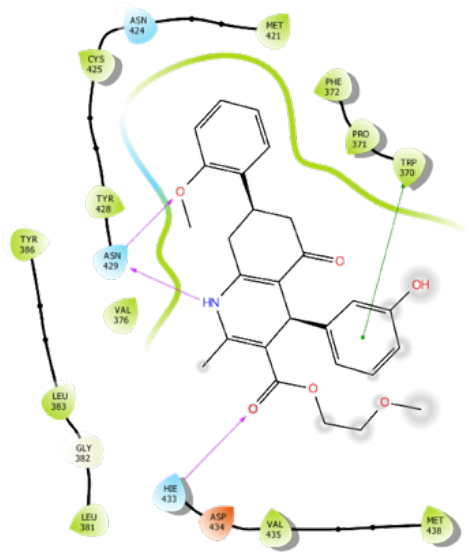
2D diagram of the proposed binding mode for HPI-1, showing the interactions with the amino acids forming the binding site. H-bonds are shown in purple lines while the green line represent π-π stacking. The grey circles show the solvent exposure and the residues in green the hydrophobic environment. Related to Fig. 5e.

### HPP-9 is a long-acting Hedgehog pathway inhibitor

Intrigued by the differential degradation efficiency found between dBet6 and HPP-9 at later timepoints, we wondered if, rather than just a target-identification tool, HPP-9 could be an interesting Hedgehog pathway inhibitor in its own right. Clearly, HPP-9 is a partial inhibitor, but since its action is through degradation rather than inhibition, we hypothesized that it could act longer than the parent compound HPI-1. To test this hypothesis, we incubated SHH-GFP cells for 25h with 1 µM of HPP-9 or HPI-1 and subsequently removed the compounds and induced the Hedgehog signaling pathway for 27h (Fig. 6a,b). Strikingly, we found that the cells that were treated with HPP-9 had strongly reduced signaling output, both at the level of a GFP reporter (Fig. 6a), as well as endogenous GLI1/GLI2 levels (Fig. 6b), whereas cells pre-treated with HPI-1 responded normally. One of the upstream activation events of the Hh pathway is the accumulation of GLI2 and GLI3 at the tip of the primary cilium, which is necessary for their subsequent activation into transcriptional activators.^10^ As HPP-9-treated cells showed strongly reduced *Gli2* transcript levels in the absence of ShhN, as well as GLI2 full-length levels below untreated cells (Fig. 2e,h), we assessed the ability of cells to accumulate GLI proteins at the ciliary tip when (pre-treated) with HPP-9 (Fig. 6c,d). For this, NIH-3T3 cells were pre-incubated for 24h with 1 µM of HPP-9 or DMSO, and subsequently treated with ShhN for 6h in the presence or absence of 1 µM HPP-9. As shown in Fig. 6c,d, we found that the globally reduced GLI2 levels translate to those found at the ciliary tip when pre-incubating cells with 1 µM HPP-9. After pre-treatment, cells could still accumulate GLI2/3 at the tip (1.5-2-fold), but overall levels were reduced by a factor 2 (GLI2) or 1.3 (GLI3). No differences were found at the 6h timepoint, indicating that this is not an acute effect on ciliary trafficking but rather a consequence of transcriptional regulation of global GLI protein levels.

**Fig. 6.**
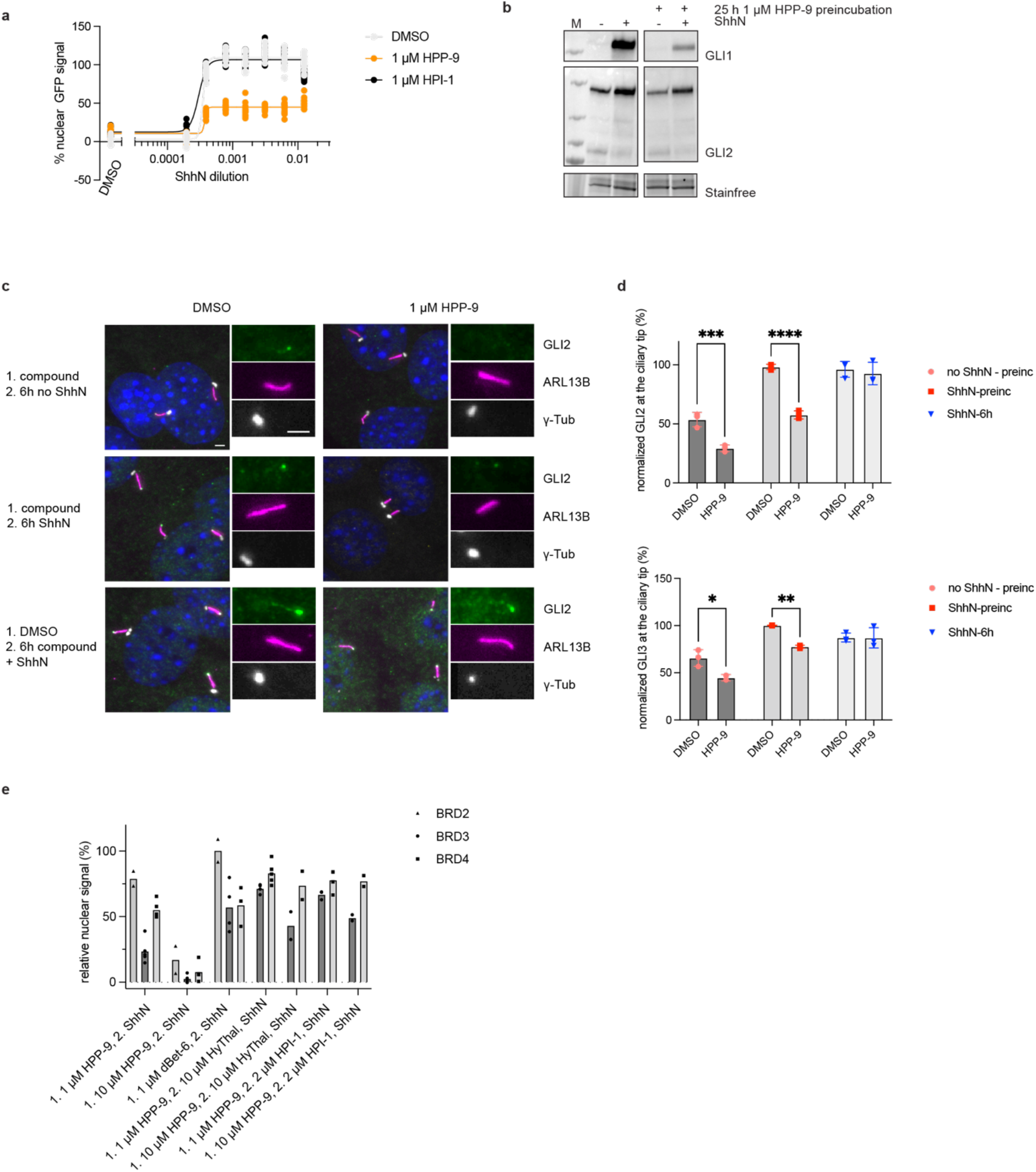
HPP-9 is a long-acting Hedgehog pathway inhibitor. a) SHH-GFP cells were treated for 26h with DMSO, 1 µM HPI-1 or 1 µM HPP-9, before the medium was changed to various concentrations of ShhN-containing medium for 28h. Nuclear GFP levels were quantified using fluorescence microscopy. Representative curves of 3 independent experiments are shown, with 9-16 images analyzed per condition. B) GLI1 and GLI2 levels of cells pre-incubated with DMSO or 1 µM HPP-9 were determined. Representative blots of 3 independent experiments are shown. C,d) The effect of HPP-9 (pre-)incubation on GLI2 and GLI3 ciliary trafficking was assessed through fluorescence microscopy. Representative images for GLI2 trafficking are shown in c), and all data is quantified in d). Three independent experiments, n=300-500 cilia analyzed per condition. Scalebar 2 µm. e) SHH-GFP cells were incubated with the indicated compounds for 26h, before the medium was changed to ShhN-containing medium for 28h. Nuclear BRD protein levels were determined using high content fluorescence microscopy. Data shown is from 2-5 independent experiments with 9-16 images analyzed per condition.

We then determined levels of BET bromodomain proteins 30h after removal of HPP-9 (1 or 10 µM) or dBet6 (1 µM). As shown in Fig. 6e pre-incubation with 10 µM HPP-9 resulted in almost complete and a much higher level of degradation compared to 1 µM HPP-9, whereas HPP-9 globally showed more degradation at this late timepoint than dBet6. As we observed such a strong Hook effect for HPP-9 compared to dBet6, we wondered if this compound could potentially be very difficult to wash out. In such a scenario, removal of the compound from the cell culture medium reduces the effective concentration to where the Hook effect is no longer in effect, rather than removing it altogether. We tested this by addition of either hydroxythalidomide (10 µM), or HPI-1 (2 µM) to the ShhN-containing medium after the pre-incubation with HPP-9 to compete with the formation of a ternary complex by HPP-9. Indeed, both compounds are able to partially restore the BET bromodomains showing that HPP-9 remains physically present in the cells after changing the medium which results in its prolonged action.

## Discussion

Target deconvolution of small molecule hits from phenotypic screening campaigns remains a major challenge and there is unfortunately no one-size fits all approach.^25^ Here, we report a PROTAC-based strategy for target identification which we applied to discover the cellular targets of Hedgehog pathway inhibitor HPI-1. Since their discovery, PROTACs have found widespread use due to their unique mechanism of action, as probes for target validation and off-target discovery, and as drugs.^38,39^ A recent preprint study furthermore describes the use of a PROTAC to identify the target of a natural product.^57^ To convert a molecule into a PROTAC, it is necessary to first determine the structure-activity relationship for the parent molecule, which is often in itself part of a medicinal chemistry campaign and also a required step in the development of for example photo-affinity probes. Through our SAR analysis we arrived at HPP-9, a PROTAC version of HPI-1 that retained the inhibitory potency of the parent molecule, but was a partial rather than a full inhibitor. Using a quantitative label-free proteomics strategy, we were able to identify the BET bromodomains as potential target candidates. A major advantage of our approach is that we could directly verify the cellular target engagement using HPP-9. Indeed, HPP-9 is a potent BRD2/3/4 degrader, whose action can be competed with HPI-1, indicating that it robustly reports on the target of the parent molecule.

BET bromodomains are epigenetic modulators involved in a plethora of cellular processes.^58,59^ Their involvement in the Hedgehog pathway has been partially deciphered through the use of the small molecule pan-BET bromodomain inhibitor JQ1, which decreases binding of BRD4 to the GLI1 and GLI2 promoters, thereby regulating GLI activity.^48^ In a genome-wide CRISPR knockout screen for modulators of ciliary Hedgehog signaling *Brd2* was found as a strong hit, whereas knockout of *Brd4* resulted in a growth phenotype.^60^ We found a couple of interesting differences when comparing HPP-9 and dBet6 action. First, HPP-9 is much less cytotoxic, second, Hh pathway inhibition is partial while full for dBet6, and third, the timing and duration of degradation is very different. This raises the question if there is a specific timepoint after pathway induction where the BET bromodomains are most critical and that HPP-9 action is simply too late to effect full inhibition. Alternatively, the differences could be the result of potential differential off-targets of these compounds. We have detected many more significantly regulated proteins in our proteomics dataset, indicating the possibility that some of these proteins are direct (off)-targets, while others may reflect the indirect consequences of BET bromodomain depletion from the cells.

While the in vitro binding affinity of HPP-9 and HPI-1 for BRD2, 3 and 4 is similar, it does appear that HPP-9 (and HPI-1), have a particular effect on cellular BRD2 which is not found for JQ1-based probes. Notably, HPP-9 differs completely from dBet6 in its degradation capacity at longer timepoints, illustrating how molecules that act on the same target can still have a different mechanism of action. We have no definitive prove at this point, but the most likely explanation for this is that HPP-9 remains stably inside the cells and is inefficiently washed out. While we found that JQ1 increased *Brd2* transcript levels much more than HPI-1, the latter did increase BRD2 protein whereas JQ1 did not. The mechanism and the functional consequences of this are currently unknown. More research will be necessary to fully understand the contributions of the various BET bromodomains on Hh signal transduction, cell viability, and the effect of compounds on these.

No PROTACs have been reported to date that act on the Hh pathway, most likely because most molecules target the transmembrane protein Smoothened. We demonstrated that BET bromodomain degradation through HPP-9 results in prolonged Hh pathway inhibition, in line with what has been shown for some other PROTACs^61^. This suggests that HPPs could form the basis for novel therapeutic modalities targeting Hedgehog pathway driven cancers. While we were focused on the effect of HPP-9 on Hedgehog signal transduction, there may be many other interesting applications of this molecule (and the parent HPI-1), especially when targeting BRD2. For example, a recent report has shown that inhibition of BRD2 can block SARS-CoV-2 infection.^62–64^

In summary, we here show that PROTACs provide a valid methodology for the target deconvolution of small molecules and are a useful addition to the chemical biology toolbox to decipher the target of phenotypic hits. We conclude that HPI-1 inhibits the Hh pathway through the inhibition of the BET bromodomains, opening new biological applications for this molecule beyond Hh pathway inhibition. Finally, targeted degradation of BET bromodomain proteins by HPP-9 is long-lasting, providing the unique opportunity of prolonged Hh pathway inhibition. We anticipate that this strategy is widely applicable, with the strong advantage that once the PROTAC is synthesized, it functions directly as a target validation tool, and, as in our case, could provide a promising pharmacological entity in itself.

## Supporting information

supplementary information containing chemical synthesis

## Acknowledgements

We would like to thank the ACCESS Geneva facility for their help with high-content microscopy, and in particular dr. Dimitri Moreau for image analysis. We kindly acknowledge the Chemical Biology Mass Spectrometry Platform (CHEMBIOMS) at the Faculty of Science, University of Geneva for the proteomics experiments. We thank James Chen for SHH-LIGHT2, SHH-GFP, HEK239T-EcR-ShhN, and SUFU-KO-LIGHT cells, David Mick for IMCD3-FlpIn cells and Aurelien Roux for HeLa cells. This research was funded by the Swiss National Science Foundation (project grant 189246 to S.H. and 186405 to L.S.) and the National Research Programme ‘NCCR Chemical Biology’, which was funded by the Swiss National Science Foundation (to S.H.).

## Contributions

M.B., H.C., Y.W. and S.H. designed the experiments and analyzed the data. M.B. and H.C. designed and synthesized the compounds. M.B. and S.H. performed the biological experiments. M.B and S.H. prepared the samples for the proteomics experiment. Y.W. performed the label-free quantitative proteomics experiment and Y.W. and S.H. analyzed the data. M.H. and L.S. performed the docking studies. M.B. and S.H. wrote the manuscript, with edits from Y.W., M.H. and L.S. L.S. and S.H. acquired funding for the project.

## Methods

### Chemical synthesis

Synthetic routes and structural characterization data for HPI-1 analogs and HPPs are described in the Supplementary Information.

### Cell lines

SHH-LIGHT2^44^, SHH-GFP^47^, HEK239T-EcR-ShhN cells and SUFU-KO-LIGHT^35^ cells were provided by James Chen (Stanford University). SHH-LIGHT2 cells were maintained in DMEM containing 10% CS, 1% sodium pyruvate, 100 U/mL penicillin, 100 μg/mL streptomycin, 150 μg/mL zeocin and 400 μg/mL G418. SHH-GFP cells were maintained in DMEM containing 10% CS, 1% sodium pyruvate, 100 U/mL penicillin, 100 μg/mL streptomycin and 150 μg/mL zeocin. SUFU-KO-LIGHT were maintained in DMEM containing 10% FBS, 1% sodium pyruvate, 100 U/mL penicillin, 100 μg/mL streptomycin and 150 μg/mL zeocin. NIH-3T3 cells were purchased from ATCC and maintained in DMEM containing 10% calf serum (CS), 1% sodium pyruvate, 100 U/mL penicillin and 100 μg/mL streptomycin. HEK293T cells (ATCC) and HEK239T-EcR-ShhN cells were maintained in DMEM containing 10% FBS, 100 U/mL penicillin and 100 μg/mL streptomycin. IMCD3-FlpIn cells were a gift from David Mick (University of Saarland) and were maintained in DMEM/F12, 10% FBS, 1% glutamine, 100 U/mL penicillin and 100 μg/mL streptomycin. HeLa cells were a gift from Aurelien Roux (University of Geneva) and maintained in DMEM containing 10% FBS, 100 U/mL penicillin and 100 μg/mL streptomycin. All cells were cultured at 37 °C with 5% CO2.

### ShhN production and tittering

HEK239T-EcR-ShhN cells were grown to 80% confluence after which the medium was changed to 2% FBS DMEM, followed by collection of conditioned medium after 48 h and filtration through a 0.22 µm filter device (Corning) The titer of ShhN was determined using SHH-LIGHT2 luciferase reporter cells (see *Luciferase reporter assays*), and a concentration approximately two-fold over the minimum dilution needed for full luciferase induction was used for further experiments.

### Luciferase reporter assays

SUFU-KO-LIGHT cells were seeded in a 96 well plate (30.000 cells/well). The next day, the medium was removed and starvation medium (DMEM w/o phenol red, 0.5% FBS) with probes in different concentrations or an equivalent amount of DMSO vehicle was added and the cells incubated for 16-18 h. Afterwards, the medium was removed and the cells were lysed for 30 min at r.t (12.2 mM Tris pH 7.4, 4% glycerol, 0.5% Triton X-100, 0.5 mg/mL BSA, 1 mM EGTA, 1 mM DTT) and the luminescence was quantified using homemade firefly luciferase reagent (0.025 M di-glycine, 0.015 M KxPO^4^ pH=8, 4 mM EGTA PH=8, 0.5 mM DTT, 0.015 M MgSO^4^, 2 mM ATP, 25 mM Coenzyme A, and 0.9 μΜ luciferin) on a multi-mode microplate reader GloMax (Promega). The data was normalized to DMSO control and curves were fitted and analyzed to determine the IC^50^ using GraphPad Prism.

SHH-LIGHT2 cells were seeded in a 96 well plate (35.000 cells/well). The next day, the medium was removed and ShhN-containing starvation medium (DMEM w/o phenol red, 0.5% CS) with compounds in the indicated concentrations or an equivalent amount of DMSO vehicle was added and the cells incubated for 28 h. Afterwards, the medium was removed and the cells were lysed for 30 mins at r.t (12.2 mM Tris pH 7.4, 4% glycerol, 0.5% Triton X-100, 0.5 mg/mL BSA, 1 mM EGTA, 1 mM DTT) and the luminescence was quantified using homemade firefly luciferase reagent (0.025 M di-glycine, 0.015 M KxPO^4^ pH=8, 4 mM EGTA PH=8, 0.5 mM DTT, 0.015 M MgSO^4^, 2 mM ATP, 25 mM Coenzyme A, and 0.9 μΜ luciferin) on a multi-mode microplate reader GloMax (Promega). Data was normalized to the DMSO with or without ShhN controls and curves were fitted and analyzed to determine the IC^50^ using GraphPad Prism.

### Cell viability assays

NIH-3T3, HEK293T, HeLa and IMCD3 cells were seeded in a 96 well plate (3k cells/well). After 5 hours, the cells were treated with the indicated compound concentrations or DMSO vehicle and incubated for 48h. Cell viability was determined using Celltiter-Blue (Promega) and fluorescence measured using multi-mode microplate reader GlowMax (Promega) using excitation and emission wavelengths of 570 nm and 600 nm, respectively. The results were analyzed using a non-linear regression (Prism 9, GraphPad Software, La Jolla, CA).

### Sample preparation for proteomic analysis

SHH-GFP cells were seeded in 3.5 cm dishes (500.000 cells/dish) and grown to confluency. The next day cells were treated with HPI-1 or HPP-9 (1 µM final) or an equivalent amount of DMSO vehicle (0.1%) in serum starvation medium (phenol red-free DMEM with 0.5% CS) for 27h (5 dishes/group). Cells were washed with PBS, trypsinized, and collected by centrifugation (500 rpm, 7 min). After centrifugation, the cell pellets were washed with PBS (1x), centrifuged (500 rpm, 7 min), and flash frozen with liquid nitrogen and stored at -80 °C until further processing. Samples were prepared for liquid chromatography/mass spectrometry (LC/MS) using the phase-transfer surfactant method, with minor modifications. First, proteins were extracted from islets and solubilized using buffer containing 12 mM sodium deoxycholate, 12 mM sodium N-dodecanoylsarcosinate, and 100 mM Tris pH 9.0, with EDTA-free Protease Inhibitor Cocktail (Roche, Switzerland). Samples were sonicated for 4 min using a Bandelin Sonorex ultrasonic bath (FAUST) with 20-sec on/20-sec off cycles. Cell debris was removed after centrifugation at 18,000 × g for 20 min at 4 °C. Protein concentrations were adjusted to a uniform concentration for a set of samples (0.5–1.0 μg/μL), and between 5 and 20 μg protein was used for peptide preparation. Cysteine–cysteine disulfide bonds were reduced with 5 mM TCEP at 37 °C for 30 min. Free thiol groups were alkylated with 20 Mm iodoacetamide in the dark at room temperature for 30 min. Alkylation reactions were quenched with 75 mM cysteine at room temperature for 10 min. Samples were diluted with 3.1 volumes of 50 mM ammonium bicarbonate. Lysyl endopeptidase (Wako, Japan) and trypsin (Promega, USA) were added at a 100:1 ratio of sample protein:enzyme (w/w) and samples were digested for 16 h at 37 °C. Afterward, 1.77 volumes ethyl acetate were added, and samples were acidified with trifluoroacetic acid (TFA), which was added to 0.46% (v/v). Following centrifugation at 12,000 × g for 5 min at room temperature, samples separated into two phases. The upper organic phase containing sodium deoxycholate was removed, and the lower aqueous phase containing digested tryptic peptides was dried using a centrifugal vacuum concentrator. Digested peptides were dissolved in 300 μL of 0.1% (v/v) TFA in 3% acetonitrile (v/v). Samples were sonicated for 1 min, centrifuged at 15,000 g for 15 min, and desalted using MonoSpin C18 columns (GL Sciences Inc., Japan). Peptides were eluted from C18 columns using 0.1% TFA in 50% acetonitrile and dried in a vacuum concentrator. Tryptic peptides were dissolved in 0.1% (v/v) formic acid in 2% (v/v) acetonitrile for MS analysis.

### MS measurements

Samples were measured using and Easy Nano LC - Orbitrap Fusion System (Thermo Fisher Scientific, USA), equipped with a PST The Nimbus ion source (Phoenix s&t). The same amount of peptide was injected for each sample in a given set of samples, which was typically 300-600 ng in a volume of 2 to 5 μL. Peptides were separated on a 3-μm particle, 75-μm inner diameter, 12-cm filling length homemade C18 column. A flow rate of 300 nL/min was used with a 2-h gradient (2–25% solvent B in 122 min, 25–45% solvent B in 4 min, 45–75% solvent B in 4 min. The gradient was followed with two rounds of washing steps, in each step, the gradient switched to 98% solvent B and keep it for another 5 min, and switched to 2% solvent B in 1 min and kept it for another 2 min. In the second round of washing, an extra 12 min of 2% solvent B was kept for system equilibration. Solvent A was 0.1% (v/v) formic acid in LC/MS grade water and solvent B was 0.1% (v/v) formic acid in 100% (v/v) acetonitrile. The ion source settings from Tune were used for the mass spectrometer ion source properties. For data-dependent acquisition (DDA), full MS spectra were acquired from 375 to 1500 m/z at a resolution of 120,000. The default charge state for the MS2 was set to 2. Charge states 2-7 were included for MS2. HCD fragmentation was set to fixed collision energy of 35%, and dynamic exclusion was set to 30 s. For both full MS and MS2, the AGC target was set to standard with a maximum injection time (IT) set to auto.

For data-independent acquisition (DIA), data were acquired with 1 full MS and 38 overlapping isolation windows constructed covering the precursor mass range of 350–1200 m/z. For full MS, Orbitrap resolution was set to 120,000. The AGC target was set to custom and maximum IT was set to 60 ms. DIA segments were acquired at 30,000 resolution with an AGC target custom and a dynamic maximum IT. HCD fragmentation was set to normalized collision energy of 27%.

### Protein identification and quantification

Raw files from DDA measurements were searched against the Uniprot mouse database using SpectroMine software (Biognosys, Switzerland) with default settings. Digestion enzyme specificity was set to Trypsin/P. Modification included carbamidomethylation of cysteine as a fixed modification, and oxidation of methionine and acetyl (protein N-terminus) as variable modifications. Up to two missed cleavages were allowed. A decoy database was included to calculate the FDR. Search results were filtered with FDR 0.01 at both peptide and protein levels. Filtered output was used to generate a sample-specific spectral library using Spectronaut software (Biognosys, Switzerland). Raw files from DIA measurements were used for quantitative data extraction with the generated spectral library. FDR was estimated with the mProphet approach and set to 0.01 at both peptide precursor level and protein level. Data were filtered with FDR < 0.01 in at least half of the samples.

### Docking of HPI-1

The crystal structure of the second bromodomain of BRD2 was downloaded from the Protein Data Bank (PDB code 7OE8^54^). The protein was then prepared with Maestro release 2021-1 (Schrödinger, LLC, New York, NY, 2021). Briefly, hydrogens were added, the H-bond assignment was optimized at pH 7.0 with PROPKA^65^ and a final minimization with the OPLS4^66^ force field was realized. Finally, all water molecules were discarded, and a grid was generated with the Grid Generation utility, centered on the co-crystallized ligand.

The ligand HPI-1 was parameterized with the LigPrep utility with the OPLS4 force field and docked with Glide within Maestro using default settings, that is standard precision and flexible ligand sampling. The 2-dimension interaction diagram was generated by Maestro and all pictures were rendered with ChimeraX^67^.

### Generation of ternary complexes

The protocol was adapted from Bai et al.^56^. Briefly, 5000 binary complexes of the second bromodomain of BRD2 (PDB 7OE8^54^) and cereblon E3 ligase (PDB 5FQD^68^ were thalidomide from PDB 4CI12^69^ replaced lenalidomide) were generated with the Rosetta software package^70^ using a local docking protocol. In both proteins, the ligands (HPI-1 for BRD2 and thalidomide for cereblon) were kept and considered rigidly as part of their respective protein. The first 500 binary complexes that displayed the highest interface score provided by Rosetta were kept for the next step.

The linker of the PROTAC and few atoms from the ligands (stubs) were extracted from HPP9 and OMEGA 4.1.0.0 (OpenEye Scientific Software, Santa Fe, NM, USA) was used to generated low-energy conformers. We randomly picked 1000 conformers for the next step.

Finally, all the linker conformers were overlapped on the binary complexes and ternary complexes were formed when the RMSD between the ligands and the stubs of the linker was less than 0.4 Å.

All complexes underwent a minimization and the lowest-energy ternary complex was taken as a physical example of a BRD2-CRBN-PROTAC ternary complex. The picture was rendered with ChimeraX^67^

### Fluorescence microscopy – Hh pathway-driven GFP reporter

Cells were seeded in 96 well microscopy plates (Ibidi). For dose-response curves, 35.000 cells/well were seeded, to reach confluency the next day. Then, the cells were treated with the indicated compound concentrations (final concentration of DMSO 0.1%) in serum starvation medium (phenol red-free DMEM with 0.5% CS) containing an appropriate dilution of ShhN-containing medium to give full pathway activation. Control wells contained medium with or without ShhN-containing medium plus the same amount of DMSO vehicle. After 24h-30h of stimulation, the medium was removed, and cells were fixed according to the protocol below. For pre-incubation experiments, cells were seeded at 20.000 cells/well, and treated with the indicated compound concentrations (final concentration of DMSO 0.1%) in serum starvation medium (phenol red-free DMEM with 0.5% CS) for 26h. The compound was removed, and the cells were stimulated with varying dilutions of ShhN-containing medium for 28h, before removing the medium and fixing the cells. Fixation was done with 4% PFA for 7 minutes at room temperature, followed by PBS washes (3x) and staining with Hoechst (Life Technologies, 1h, 1:10000 in PBS). Cells were imaged using a 20x water immersion objective on a high content confocal microscope (Molecular Devices™ ImageXpress Micro XL).

### Immunofluorescence Imaging

For BRD2/3/4 immunostaining, fixed cells from the fluorescence microscopy experiment were permeabilized with 0.5% Triton X-100 for 5 min at room temperature, washed with 0.1% Triton X-100 in PBS (PBS-T, 3x), and blocked using 1% BSA in PBS-T (blocking solution) for 1h at room temperature. Cells were incubated with primary antibody in blocking solution (1:500 rabbit anti-BRD2 (Bethyl Laboratories, A302-583A), 1:500 mouse anti-BRD3 (Santa Cruz Biotechnology Inc., 2088C3a) 1:1000 rabbit anti-BRD4 (Bethyl Laboratories, A301-985A) overnight at 4 °C, washed 5x 5 minutes with PBS-T, and incubated with appropriate secondary antibodies in blocking solution (1:500 Jackson ImmunoResearch) for 1h at room temperature. Cells were washed 5x 5 minutes with PBS-T and once with PBS before being imaged using a 20x water immersion objective on a high content confocal microscope (Molecular Devices™ ImageXpress Micro XL). For GLI2/GLI3 trafficking studies, cells were grown to confluency, serum-starved for 20 h in the presence of 1 µM compound or DMSO vehicle, followed by 6 h incubation in the presence or absence of the appropriate amount of ShhN-conditioned medium and 1 µM compound or DMSO as indicated. Cells were sequentially fixed with 4% PFA for 10 minutes and ice-cold methanol for 5 minutes, before being washed with PBS (3x), and blocked in blocking solution for 1h at room temperature. Cells were incubated with primary antibody in blocking solution (1:500 mouse anti-gamma tubulin (Sigma Aldrich, GTU-88), 1:3000 mouse anti-ARL13B (Biolegend, clone N295B/66), 1:500 Goat anti-mouse GLI2 (R&D systems, AF3635), 1:500 Goat anti-mouse GLI3 (R&D systems, AF3690)) overnight at 4 °C, washed 5x 5 minutes with PBS-T, and incubated with appropriate secondary antibodies in blocking solution (1:500 Jackson ImmunoResearch) for 1h at room temperature. Cells were washed 5x 5 minutes with PBS-T and once with PBS before being imaged using a 60x water immersion objective on a high content confocal microscope (Molecular Devices™ ImageXpress Micro XL).

### Quantification of fluorescence microscopy images

Image analysis was performed using MetaXpress software, and a custom Matlab (Mathworks) script. Local background subtraction was performed on all images before analysis. For nuclear signal, the Hoechst stain was used to mask the nuclei, and the signal intensity within the mask determined for the other channels. Data was plotted and curves fitted using GraphPad Prism 9. Data was normalized to DMSO controls (with and without ShhN). To determine GLI levels at the tip of the primary cilium, the ARL13B channel was used to create a ciliary mask. The ciliary mask was then used to identify and measure ciliary signal in the other channels. The γ-tubulin signal (as a centriole marker) was used to orient all cilia from base to tip. Each cilium was divided in 10 bins, and the tip fluorescence for GLI2 and GLI3 was defined as the summed fluorescence in the final five bins of each cilium, regardless of length.

### Western blotting

For signaling assays, SHH-GFP cells were seeded in a 24-well plate and grown until confluency. Growth medium was then replaced with serum starvation medium with or without ShhN conditioned medium and compounds at the appropriate dilution. After 28 h, cells were lysed in SDS sample buffer (50 mM Tris HCl pH 6.8, 8% v/v glycerol, 2% w/v SDS, 100 mM DTT, 0.1 mg/mL bromophenol blue), boiled and sonicated.

For competition assays, SHH-GFP cells were seeded in a 24-well plate and grown until confluency. Growth medium was then replaced with serum starvation medium containing DMSO or 1 µM HPP-9 plus the indicated concentrations of competitors for 7h. Cells were washed with PBS, lysed in SDS sample buffer, boiled and sonicated.

For time-course assays, all wells were changed to serum-starvation at the time compound treatment was started for the longest timepoint, and the medium was changed to compound-containing medium at the indicated times, before lysing all wells at the same time.

Samples were loaded onto a 4-15% Criterion TGX Stainfree gel (Bio-Rad), and run for 45 min, 200V in Tris/Glycine/SDS buffer (Bio-Rad). Gels were irradiated (1 min) and the stain free imaged, before being transferred onto a PVDF membrane using a Transblot Turbo system (Bio-Rad). Membranes were blocked in 5% milk in 0.1% Tween-20 in Tris-buffered saline (TBST) for 1 h at room temperature, and subsequently incubated with the indicated primary antibody in blocking buffer for 16 h at 4 °C. Primary antibodies used:

Goat anti-mouse Gli3 (R&D systems, AF3690), 1:200; Mouse anti-Gli1 (Cell signaling, 2643S), 1:1000; Goat anti-mouse Gli2 (R&D systems, AF3635), 1:1000; Mouse anti-vinculin (Proteintech, 66305-1), 1:5000; Mouse anti-BRD3 (Santa Cruz, sc-81202), 1:200; Rabbit anti BRD4 (Bethyl laboratories, A301-985A-M), 1:1000; Rabbit anti BRD2 (Bethyl laboratories, A302-583A-T), 1:1000. Membranes were washed (3x 10 min TBST), incubated with HRP-conjugated secondary antibody, washed again, developed using Supersignal West Atto Maximum Sensitivity Substrate (Thermo Fisher) and imaged on a Fusion FX geldoc (Vilber). Membranes were stripped using Restore Western Blot stripping buffer (Thermo-Fisher) and re-probed as described above. Band intensities were determined using Fiji Image J (National Insitute of Health) on background substracted images and normalized to total protein loaded using an appropriate housekeeping protein or stainfree total protein.

### Real-time quantitative PCR

SHH-GFP cells were seeded in a 12-well plate and grown until confluency. Growth medium was then replaced with serum starvation medium with or without ShhN conditioned medium and compounds or DMSO vehicle at the appropriate dilution. After 24h, cells were washed with ice-cold PBS and RNA extracted using the Reliaprep RNA Cell miniprep system (Promega). The obtained RNA was reverse transcribed using the First strand DNA synthesis kit, random primers (Promega). The cDNA was diluted and used with TaqMan Fast Advanced Master Mix (Applied Biosciences) and the appropriate Taqman primers (Life Technologies) in a CFX qPCR thermocycler (Bio-Rad). Cycle numbers were normalized to the housekeeping gene B2M or Gapdh. Taqman primers used: Mm00494654_m1 mGli1; Mm01293117_m1 mGli2; Mm00436026_m1 mPtch1; Mm00437762_m1 mB2M; Mm99999915_g1 Gapdh; Mm01271171_g1 mBrd2

