## supplementary information containing chemical synthesis for "Targeted protein degradation reveals BET bromodomains as the cellular target of Hedgehog Pathway Inhibitor-1"

**Materials and Methods**

Briefly, reagents for synthesis were purchased from Fluka, Sigma-Aldrich, TCI and Across. Salts of the best grade available from Fluka or Sigma-Aldrich were used as received. Column chromatography was carried out on silica gel 60 (SilicaFlash P60, 40-63 µm). Analytical (TLC) and preparative thin layer chromatography (PTLC) were performed on silica gel 60 (Merck, 0.2 mm) and silica gel GF (SiliCycle, 1 or 0.25 mm), respectively. Reverse phase flash chromatography was performed on a Biotage® Isolera Spektra using pre-packed 12 g Biotage® SNAP Ultra C18 cartridge. LCMS were recorded using a Thermo Scientific Accela HPLC equipped with a Thermo C18 Hypersil GOLD column (50 ´ 2.1 mm, 1.9 µm particles size) coupled with a LCQ Fleet three-dimensional ion trap mass spectrometer (ESI, Thermo Scientific) with a linear elution gradient from 95% $H_2O$ / 5% $CH_3CN$ + 0.1% TFA to 10% $H_2O$ / 90% $CH_3CN$ + 0.1% TFA in 4.0 minutes at a flow rate of 0.75 mL/min (B5), 70% $H_2O$ / 30% $CH_3CN$ + 0.1% TFA to 10% $H_2O$ / 90% $CH_3CN$ + 0.1% TFA in 4.0 minutes at a flow rate of 0.75 mL/min (B30) or 40% $H_2O$ / 60% $CH_3CN$ + 0.1% TFA to 10% $H_2O$ / 90% $CH_3CN$ + 0.1% TFA in 4.0 minutes at a flow rate of 0.75 mL/min (B60). All $^1H$ and $^{13}C$ NMR spectra were recorded (as indicated) on a Bruker 300 MHz, 400 MHz or 500 MHz spectrometer at room temperature (25 °C) and are reported as chemical shifts (δ) in ppm relative to TMS (δ = 0). Spin multiplicities are reported as a singlet (s), doublet (d), triplet (t), quartet (q), and quintet (p) with coupling constants (J) given in Hz, or multiplet (m). Broad peaks are marked as br. 1 H and 13C resonances were assigned with the aid of additional information from 1D and 2D NMR spectra (H,H-NOESY, H,H-COSY, DEPT 135, HSQC and HMBC). ESI-HRMS for the characterisation of new compounds was measured on Xevo G2-S Tof (Waters). All mass data are reported as mass-per-charge ratio m/z (intensity in %, [assignment]).

**Abbreviations**. $BMIMBF_4$: 1-Butyl-3-methyl-imidazolium-tetrafluoroborate; DIAD: Diisopropyl azodicarboxylate; DIPEA: N,N-Diisopropylethylamine; DMF: Dimethylformamide; DMSO: Dimethyl sulfoxide; HATU: Hexafluorophosphate azabenzotriazole tetramethyl uronium; HBTU: 3 [Bis (dimethylamino) methyliumyl]-3 *H* -benzotriazole-1-oxide hexafluorophosphate; n-Buli: n-Butylithium; $PPh_3$: Triphenylphosphine; rt: Room temperatureTBAF: Tetra-n-butylammonium fluoride; TBSMSCl: Tert-Butyldimethylsilyl chloride TBTA: Tris((1- benzyl-1H-1,2,3-triazol-4-yl)methyl)amine; TEA: Triethylamine; TFA: Trifluoroacetic acid;THF: Tetrahydrofuran; TsCl: 4-Toluenesulfonyl chloride.

**Synthesis**

### Multicomponent Hantzsch reaction

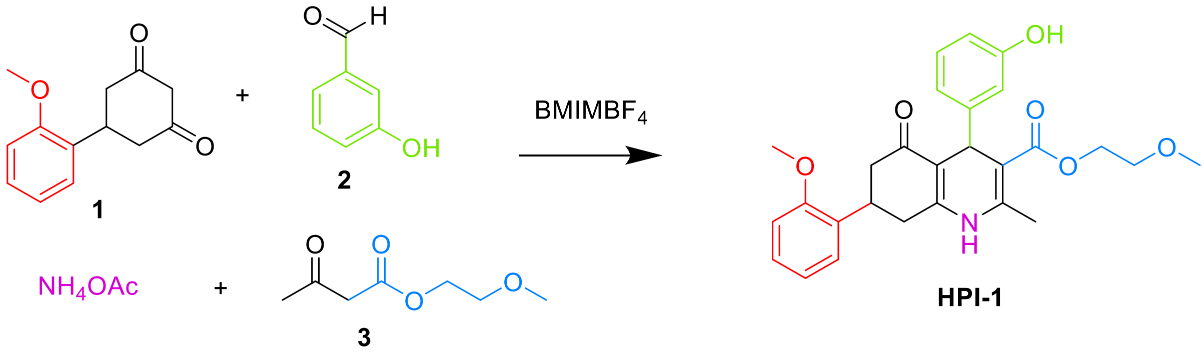


**Scheme S1.** Synthesis of HPI-1 through Hantzsch reaction

### Hantzsch reaction building blocks

Compound **1** was prepared following previously reported procedures[1]

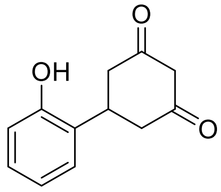


**4**

**Compound 4**. In a tricolon flask the dione **1** (500 mg, 2.3 mmol, 1 eq.) was added and then the system was put under $N_2$. Next, it was dissolved in anhydrous $CH_2Cl_2$ (32 ml) and cooled to 0 °C. A 1.0 M $BBr_3$ solution in $CH_2Cl_2$ (6.9 ml, 6.9 mmol, 3 eq.) was added dropwise and slowly and the resulting mixture stirred for two days at rt. The reaction mixture was quenched with cold water and the forming precipitate was filtered and dried in Büchner funnel to give **4** as a white powder (130 mg, 28%). $^1$H NMR (400 MHz, $CD_2Cl_2$): δ 6.97-7.09 (m, 2H), 6.68-6.81 (m, 2H), 3.59-3.65 (m, 1H), 3.27 (s, 2H), 2.62-2.72 (m, 2H), 2.44-2.54 (m, 2H) ppm.

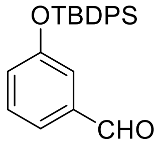


**5**

**Compound 5.** Tert-butyldiphenylsilyl chloride (0.56 mL, 2.4 mmol) was added dropwise to a stirred solution of 3-hydroxybenzaldehyde (300 mg, 2 mmol) and imidazole (300 mg, 4.4 mmol) in DMF

(1 mL) at room temperature. The reaction mixture was stirred for 10 h. Subsequently, water (2 mL) was added and the resulting mixture was extracted with diethyl ether (5 mL) three times. Combined organic layers were dried over anhydrous $MgSO_4$ and concentrated under high vacuum. The crude product was purified by column chromatography on silica gel (Pentane/EtOAc, 10:1, $R_f$ =0.45) to afford the aldehyde **5** as a colorless oil (690 mg, 96% yield). $^1$H NMR (400MHz, $CDCl_3$): δ 9.82 (s, 1H), 7.69-7.73 (d, 4H), 7.40-7.47 (m, 7H), 7.35-7.39 (m, 1H), 7.19-7.30 (m, 1H), 6.93-6.97 (dd, 1H), 1.12 ( s, 9H) ppm.

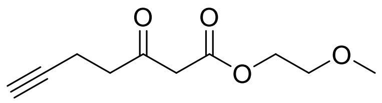

**6**

**Compound 6**. A heterogeneous mixture of NaH (60% suspension in mineral oil) (420 mg, 10.5 mmol, 2.1 eq) in dry THF (8 mL) at 0 °C was slowly added to 2-methoxyethyl acetoacetate (0.8 mL, 5 mmol, 1.0 eq). After 30 min the yellow heterogeneous mixture was added to the solution of n-BuLi (1.6 M in hexane) (3.75 mL, 6 mmol, 1.2 eq) at 0 °C and the resulting deep red-orange heterogeneous mixture was allowed to stir for an additional 30 min before it was cooled to -78 °C. Propargyl bromide (0.7 mL, 6.5 mmol, 1.3 eq) was then added slowly to the reaction mixture. The reaction continued to stir for 2 h before it was quenched by slow addition of water. The mixture was extracted with EtOAc and the combined organic phases were washed with sat. aq. NaCl and dried over anh. $Na_2SO_4$, concentrated under reduced pressure and purified by silica gel column chromatography (Petroleum ether/EtOAc 2.5:1, $R_f$ =0.38) to yield the desired product as a pale-yellow oil (400 mg, 40 %). $^1$H NMR (400 MHz, $CDCl_3$) δ 4.30 (dd, $J$ = 5.5, 3.7 Hz, 2H), 3.61 (dd, $J$ = 5.6, 3.9 Hz, 2H), 3.52 (s, 2H), 3.38 (s, 3H), 2.82 (t, $J$ = 7.5 Hz, 2H), 2.47 (td, $J$ = 7.4, 2.5 Hz, 2H), 1.96 (q, $J$ = 2.7, 2.2 Hz, 1H).

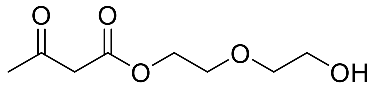

**7**

**Compound 7**. A solution of diethylene glycol (743 mg, 7 mmol) and 2,2,6-trimethyl-4H-l,3-dioxin-4-one (1 g,7 mmol) in 1.4 mL of xylene was placed in a 50-mL Erlenmeyer flask. The flask was immersed in an oil bath that had been preheated to 150 °C, and the solution was vigorously stirred. The evolution of acetone became apparent within several minute, heating was continued for a total of 30 min. The reaction was cooled, the xylene was removed in vacuo. Purification by silica gel column chromatography ($CH_2Cl_2$/MeOH 100:1, $R_f$ =0.44) to afford compound **7** as a yellow oil. $^1$H NMR (400 MHz, $CDCl_3$) δ 4.35 – 4.30 (m, 2H), 3.76 – 3.70 (m, 4H), 3.62 – 3.58 (m, 2H), 3.49 (d, $J$ = 3.4 Hz, 2H), 2.27 (s, 3H), 1.91 (s, 1H).

**HPI-1 (General procedure for Hantzsch reaction)**[1]. **HPI-1.** To a dry round-bottom flask was added **1** (0.100 g, 0.460 mmol), **2** (56 mg, 0.46 mmol), **3** (73 mg, 0.46 mmol), ammonium acetate (35 mg, 0.46 mmol), and the ionic liquid $BMIMBF_4$ (10 µL). The mixture was stirred for 15 min at 90 °C, after which it was cooled to rt and directly purified by silica gel column chromatography (Pentane/EtOAc 3:7, $R_f$ = 0.40) to give **HPI-1** as a pale-yellow solid (140 mg, 66%). $^1$H NMR (400 MHz, $CDCl_3$): δ 7.22 − 6.61 (m, 8H), 6.21 (br, 1H, NH), 5.14 (s, 1H), 4.25 − 4.13 (m, 2H), 3.79 (s, 3H), 3.64 − 3.57 (m, 1H), 3.56 − 3.53 (m, 2H), 3.32 (s, 3H), 2.77 − 2.54 (m, 4H), 2.37 (s, 3H). $^{13}$C NMR (101 MHz, $CDCl_3$): δ 196.2, 167.3, 157.1, 155.7, 150.4, 148.3, 143.9, 130.2, 129.1, 128.1, 127.1, 120.7, 120.1, 115.3, 113.3, 112.6, 110.7, 105.9, 70.5, 62.8, 58.7, 55.2, 42.2, 36.2, 33.2, 33.0, 19.5. LC-MS ($ES^+$): m/z 464.09 $[M+H]^+$.

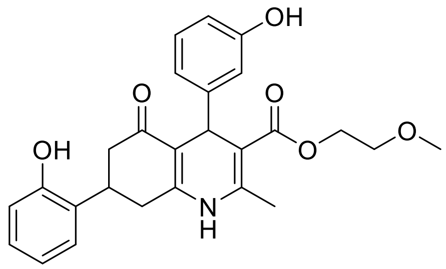


**HPI-1-A**

**HPI-1-A** was synthesized following the general procedure for Hantzsch reaction, engaging **2** (33 mg, 0.22 mmol), **3** (35 mg, 0.22 mmol), **4** (45 mg, 0.22 mmol), ammonium acetate (17 mg, 0.22 mmol), and the ionic liquid $BMIMBF_4$ (5 µL). The resulting crude was purified by column chromatography (Pentane/EtOAc 1:1, $R_f$ =0.3) to yield **HPI-1-A** as a yellow powder (33 mg, 33%). $^1$H NMR (400 MHz, MeOD and $CDCl_3$): δ 6.92-6.38 (m,8H), 5.14 (s, 1H), 4.18 (m, 2H), 3.80 (s, 3H), 3.55(m, 2H), 3.32 (s, 3H), 2.56-2.47 (m, 4H), 2.16 (s,2H) ppm. LC-MS ($ES^+$): m/z 450.08 $[M+H]^+$.

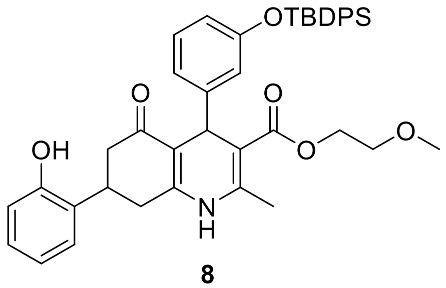


**8**

**Compound 8** was synthesised following the general procedure for Hantzsch reaction, engaging **5** (80 mg, 0.22 mmol), **3** (35 mg, 0.22 mmol), **4** (45 mg, 0.22 mmol), ammonium acetate (17 mg, 0.22 mmol), and the ionic liquid $BMIMBF_4$ (5 µL). The resulting crude was purified by column chromatography (Pentane/EtOAc 1:1, $R_f$ =0.45) to yield 8 as a yellow powder (30 mg, 20%). LC-MS ($ES^+$): m/z 688.17 $[M+H]^+$.

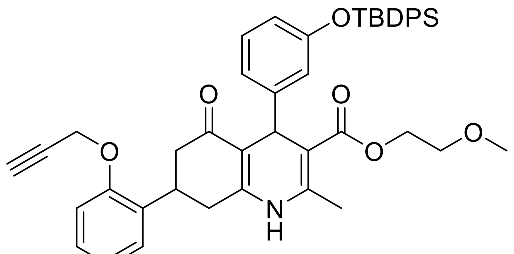

**9**

**Compound 9. 8** (30 mg, 0.044 mmol) was dissolved in 5 ml of acetone and 1.4 eq of anhydrous potassium carbonate (8.5 mg, 0.061 mmol) was added and refluxed for 45 min. Then 2 eq of propargyl bromide (0.009 mL, 0.088 mmol) were added to the mixture and refluxed for 16h. The reaction was monitored by LC-MS until the consumption of the starting material. The solvent was evaporated in vacuo and the residue redissolved in EtOAc and washed with water 3 times and once with brine to yield **9** as a brown oil (28 mg, 89%). LC-MS ($ES^+$): m/z 726.22 $[M+H]^+$.

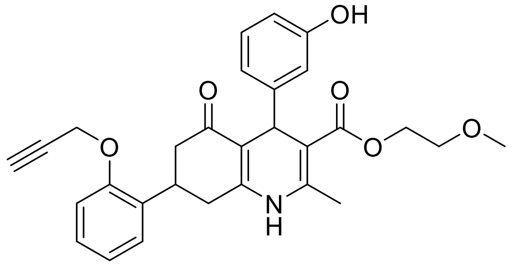

**HPI-1-B**

**HPI-1-B. 9** (28 mg, 0.039 mmol, 1.0 eq) was dissolved in THF (0.5 mL). To this solution was added 1.0 M TBAF (0.077 mL, 0.077 mmol in THF, 2.0 eq) and the mixture was stirred at 40 °C overnight under an argon atmosphere. The resulting solution was concentrated in vacuo and the resulting oil was taken up in EtOAc. The solution was washed with water (× 3), washed with saturated $NH_4Cl$ (x5) to remove the salts, dried with $MgSO_4$, and concentrated in vacuo. The resulting crude was purified by column c hromatography ($CH_2Cl_2$/MeOH 15:1, $R_f$ =0.34) to give **HPI-1-B** as a yellow oil (10 mg or 53%). $^1$H NMR (400 MHz, $CDCl_3$) δ 7.25 – 7.15 (m, 1H), 7.11 (dd, *J* = 7.7, 1.7 Hz, 1H), 7.07 – 6.92 (m, 3H), 6.91 – 6.79 ( m, 2H), 6.67 (s, 1H), 6.60 (ddd, *J* = 8.1, 2.6, 1.0 Hz, 1H), 6.53 – 6.48 (m, 1H), 5.11 (d, *J* = 15.6 Hz, 1H), 4 .67 (q, *J* = 2.1, 1.5 Hz, 2H), 4.26 – 4.08 (m, 2H), 3.54 (td, *J* = 4.8, 3.2 Hz, 3H), 3.30 (d, *J* = 3.6 Hz, 3H), 2.7 6 – 2.42 (m, 5H), 2.29 (d, *J* = 12.2 Hz, 3H), 1.79 (s, 1H). LC-MS ($ES^+$): m/z 488.09 $[M+H]^+$.

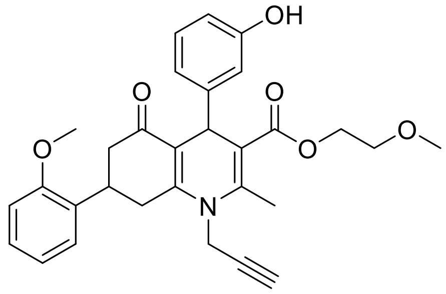


**HPI-1-C**

**HPI-1-C**. A solution of **1** (460 mg, 2mmol) ,**2** (226 mg, 2mmol), **3** (320 mg, 2 mmol) and propargylamin e hydrochloride (184 mg, 2 mmol) in pyridine (2 mL) was refluxed for 24 hours. Then the reaction mix ture was poured in cold water and extracted 3 times with EtOAc, dried with $MgSO_4$, filtered, and con centrated under reduced pressure. Finally, it was purified by column chromatography (Pentane/EtOA c 6:4, $R_f$= 0.48) to give **HPI-1-C** as a yellow powder (230 mg, 15%). $^1$H NMR (400 MHz, CDCl3): δ 7.26-6.59 (m, 8H), 5.17 (s, 1H), 4.35-4.18 (m, 4H), 3.80 (s, 3H), 3.60-3.65 (m, 1H), 3.55(m, 2H), 3.32 (s, 3H), 3.06-3.10 (m, 1H 2.89-2.65 (m, 3H), 2.64 (s, 3H), 2.42 (s,1H) ppm. LC-MS ($ES^+$): m/z 502.02 $[M+H]^+$.

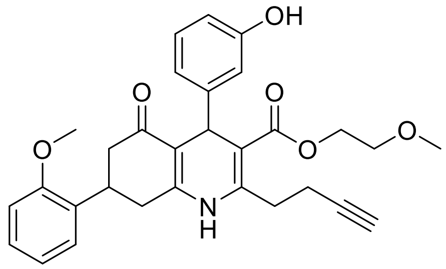


**HPI-1-D**

**HPI-1-D** was synthesised following the general procedure for Hantzsch reaction, engaging **1** (65 mg, 0.3 mmol), **2** (37 mg, 0.3 mmol), **6** (60 mg, 0.22 mmol), ammonium acetate (23 mg, 0.3 mmol), and the ionic liquid $BMIMBF_4$ (7 µL). The resulting crude was purified by column chromatography (Pentane/EtOAc 4:7, $R_f$ =0.41) to yield **HPI-1-D** as a yellow oil (62 mg, 41%). $^1$H NMR (400 MHz, $CDCl_3$) δ 7.21 (d, *J* = 0.9 Hz, 1H), 7.08 (d, *J* = 16.4 Hz, 1H), 6.99 (dd, *J* = 2.6, 1.5 Hz, 1H), 6.96 – 6.87 (m, 2H), 6.84 (ddd, *J* = 8.0, 6.9, 1.1 Hz, 2H), 6.61 (ddd, *J* = 8.1, 2.6, 1.0 Hz, 1H), 5.12 (d, *J* = 18.1 Hz, 1H), 4.25 – 4.06 (m, 2H), 3.77 (d, *J* = 5.7 Hz, 3H), 3.65 – 3.56 (m, 1H), 3.53 (t, *J* = 4.8 Hz, 2H), 3.30 (d, *J* = 3.9 Hz, 3H), 3.14 – 3.02 (m, 1H), 2.80 – 2.46 (m, 7H), 2.04 (q, *J* = 2.9 Hz, 1H). LC-MS ($ES^+$): m/z 502.299 $[M+H]^+$.

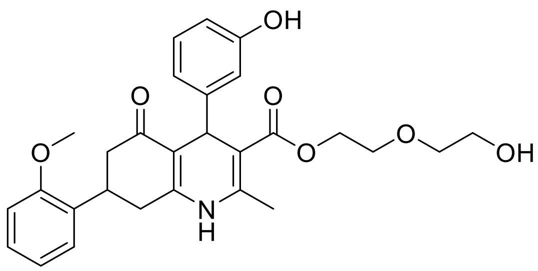

**HPI-1-E**

**HPI-1-E** was synthesised following the general procedure for Hantzsch reaction, engaging **1** (65 mg, 0 .3 mmol), **2** (37 mg, 0.3 mmol), **7** (57 mg, 0.22 mmol), ammonium acetate (23 mg, 0.3 mmol), and the ionic liquid $BMIMBF_4$ (7 µL). The resulting crude was purified by column chromatography ($CH_2Cl_2$/Me OH 100:6, $R_f$ =0.52) and then reverse phase chromatography (BGB Scorpius C18 4.5 g, $H_2O$ + 0.1% TFA /$CH_3CN$ + 0.1% TFA 70:30 to 20:80) to yield **HPI-1-D** as a pale yellow powder (32 mg, 22%). $^1H$ NMR (5 00 MHz, MeOD) δ 7.25 – 7.12 (m, 2H), 7.05 – 6.90 (m, 3H), 6.79 (d, *J* = 1.8 Hz, 2H), 6.59 – 6.50 (m, 1H) , 5.01 (d, *J* = 24.1 Hz, 1H), 4.24 – 4.10 (m, 2H), 3.80 (d, *J* = 8.0 Hz, 3H), 3.66 – 3.62 (m, 3H), 3.60 – 3.40 (m, 3H), 2.86 – 2.39 (m, 4H), 2.36 (d, *J* = 8.1 Hz, 3H). LC-MS ($ES^+$): m/z 494.03 $[M+H]^+$.

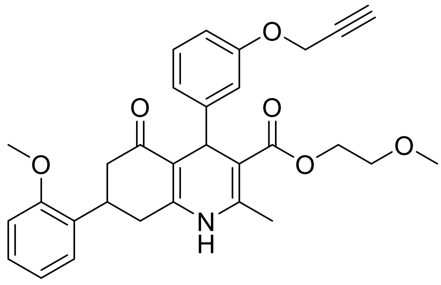

**HPI-1-F**

**HPI-1-F.** To a solution of **HPI-1** (424 mg, 0.91 mmol) in acetone (74 mL) was added anhydrous $K_2CO_3$ (177 mg, 1.28 mmol) followed by propargyl bromide (80% w/v in toluene, 0.16 mL, 1.83 mmol). The r eaction mixture was refluxed overnight, after which it was cooled and concentrated *in vacuo*. The res ulting residue was re-dissolved in EtOAc, washed with $H_2O$, dried over $MgSO_4$, and concentrated. The resulting crude was purified by silica gel column chromatography (pentane/EtOAc 3:2, $R_f$ = 0.32) and reverse phase chromatography (BGB Scorpius C18 4.5 g, $H_2O$ + 0.1% TFA/$CH_3CN$ + 0.1% TFA 70:30 to 20:80) to give **HPI-1-F** as a pale-yellow solid (270 mg, 59%).$^1H$ NMR (400 MHz, $CDCl_3$): δ 7.25 − 6.71 ( m, 8H), 6.07 (br, 1H, NH), 5.16 (s, 1H), 4.66 − 4.58 (m, 2H), 4.20 – 4.14 (m, 2H), 3.78 (s, 3H), 3.66 − 3.5 8 (m, 1H), 3.52 (t, *J* = 5.1 Hz, 2H), 3.31 (s, 3H), 2.77 − 2.54 (m, 4H), 2.53 (t, *J* = 2.4 Hz, 1H), 2.38 (s, 3H). $^{13}C$ NMR (101 MHz, $CDCl_3$): δ 195.6, 167.2, 157.6, 157.1, 149.5, 148.7, 143.9, 130.4, 128.8, 128.0, 127 .0, 121.7, 120.7, 114.9, 112.8, 112.0, 110.7, 105.7, 78.8, 75.3, 70.5, 62.9, 58.9, 55.7, 55.1, 42.4, 36.4, 3 3.2, 33.0, 19.5. LC-MS ($ES^+$): m/z 502.08 $[M+H]^+$.

**VHL ligand** was prepared following previous reported procedures [2,3].

**Pomalidomide** was prepared following previous reported procedures[4].

**Hydroxythalidomide** and **compound 10** were prepared following previous reported procedures[5].

Linker 1 was prepared following previous reported procedures [6].

Linker 4 was prepared following previous reported procedures[7].

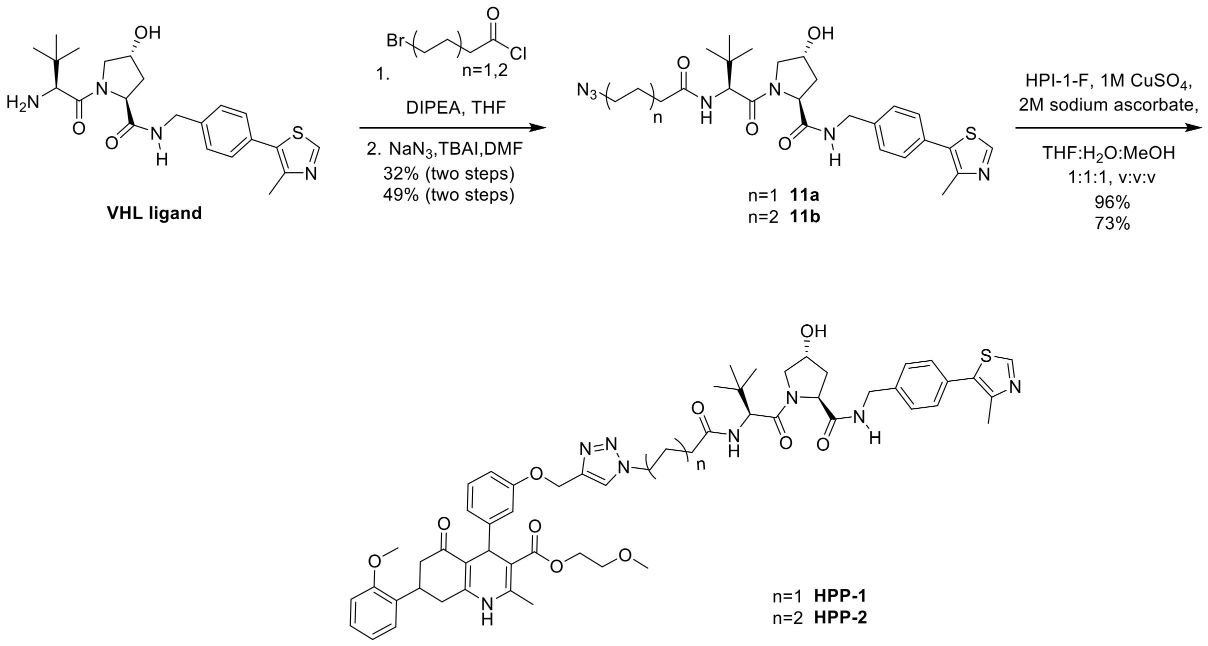


**Scheme S2.** Synthesis of **HPP-1** and **HPP-2**.

**Compound 11a.** To a solution of **VHL ligand** (10 mg, 0.23 mmol) in anhydrous THF (1.0 mL) was added DIPEA (80 µL, 0.46 mmol) and stirred for 10 min at 0 °C. 4-bromobutanoyl chloride (54 µL, 0.46 mmol) was added slowly and stirred for 2 hours at 0 °C. Upon completion, $CH_2Cl_2$/MeOH (9:1) solution was added to the reaction mixture. The organic layer was washed with $H_2O$, dried over $MgSO_4$, and concentrated to give the crude as a brown solid; $R_f$= 0.52 ($CH_2Cl_2$/MeOH 10:1). LC-MS ($ES^+$): m/z 578.81 $[M+H]^+$. The resulting crude was re-dissolved in anhydrous DMF (0.3 mL). $NaN_3$ (34 mg, 0.52 mmol) and TBAI (3 mg, 5 mol %) were added and the mixture was stirred at 80 ℃ for 16 h. After cooling to rt, the reaction mixture was extracted with $CH_2Cl_2$ (×3). The organic layer was dried over $MgSO_4$ and concentrated. The resulting crude was purified by PTLC ($SiO_2$, $CH_2Cl_2$/MeOH 15:1, eluted once) to give **11a** as an off-white solid (40 mg, 32% two steps).; $R_f$= 0.32 ($CH_2Cl_2$/MeOH 15:1). $^1$H NMR (400 MHz, $CDCl_3$): δ 8.71 (s, 1H), 7.39 – 7.33 (m, 4H), 6.11 (d, *J* = 9.0 Hz, 1H), 4.73 (t, *J* = 7.6 Hz, 1H), 4.61 – 4.54 (m, 2H), 4.47 (s, 1H), 4.33 (d, *J* = 15.7 Hz, 1H), 4.08 (d, *J* = 10.9 Hz, 1H), 3.60 (d, *J* = 11.3 Hz, 1H), 3.33 (t, *J* = 6.5 Hz, 2H), 2.62 – 2.56 (m, 1H), 2.53 (s, 3H), 2.31 (t, *J* = 6.8 Hz, 2H), 2.15 – 2.09 (m, 1H), 1.89 (p, *J* = 6.9 Hz, 2H), 0.93 (s, 9H). LC-MS (ES$^+$): m/z 541.85 $[M+H]^+$.

**Compound 11b. VHL** ligand (0.16 g, 0.37 mmol) was reacted with 6-bromohexanoyl chloride (113 µL, 0.74 mmol) according to the procedure described above. The crude product was purified by PTLC ($SiO_2$, $CH_2Cl_2$/MeOH 15:1, eluted once) to give **11b** as a white solid (104 mg, 49% two steps).; $R_f$= 0.25 ($CH_2Cl_2$/MeOH 15:1). $^1$H NMR (400 MHz, $CDCl_3$): δ 8.71 (s, 1H), 7.39 – 7.33 (m, 4H), 6.03 (s, 1H), 4.73 (t, *J* = 8.0 Hz, 1H), 4.61 – 4.53 (m, 2H), 4.48 (s, 1H), 4.33 (d, *J* = 14.9 Hz, 1H), 4.12 (d, *J* = 11.3, 1H), 3.60 (d, *J* = 10.9 Hz, 1H, OH), 3.26 (t, *J* = 6.9 Hz, 2H), 2.62 – 2.56 (m, 1H), 2.53 (s, 3H), 2.21 (t, *J* = 7.3 Hz, 2H), 2.15 – 2.10 (m, 1H), 1.64 – 1.57 (m, 4H), 1.39 – 1.37 (m, 2H), 0.93 (s, 3H).LC-MS (ES$^+$): m/z 569.86 $[M+H]^+$.

**HPP-1.** To a solution of **11a** (26 mg, 0.05 mmol) and **HPI-1-F** (24 mg, 0.05 mmol) in degassed THF/$H_2O$/MeOH (2.4 mL, 1:1:2) were added degassed aqueous solutions of 2 M sodium ascorbate (48 µL, 0.10 mmol) and 1 M $CuSO_4$ (10 µL, 0.01 mmol, 20 mol%). The mixture was stirred at rt overnight, after which it was concentrated under reduced pressure. $CH_2Cl_2$/MeOH (9:1) and $H_2O$ were added to the reaction mixture and after partitioning of the layers, the aqueous layer was extracted with $CH_2Cl_2$/MeOH (9:1). The organic layer was dried over $MgSO_4$ and concentrated. The resulting crude was purified by PTLC ($SiO_2$, $CH_2Cl_2$/MeOH 12:1, eluted once) to give **HPP-1** as a white solid (48 mg, 96%).; $R_f$= 0.28 ($CH_2Cl_2$/MeOH 12:1). $^1$H NMR (500 MHz, $CD_3OD$) δ 8.85 (s, 1H), 8.06 (s, 1H), 7.49 – 7.35 (m, 4H), 7.27 – 6.65 (m, 8H), 5.19 – 4.99 (m, 3H), 4.61 – 4.46 (m, 4H), 4.43 (m, 2H), 4.34 (dd, *J* = 15.4, 4.6 Hz, 1H), 4.14 (m, 2H), 3.91 (dd, *J* = 11.2, 1.8 Hz, 1H), 3.84 – 3.75 (m, 1H), 3.80 (s, 3H), 3.60 – 3.47 (m, 3H), 3.28 (s, 3H), 2.85 – 2.54 (m, 3H), 2.48 – 2.39 (m, 1H), 2.46 (s, 3H), 2.35 (s, 3H), 2.29 (m, 2H), 2.25 – 2.16 (m, 3H), 2.12 – 2.03 (m, 1H), 1.03 (s, 9H). $^{13}$C NMR (126 MHz, $CD_3OD$) δ 197.2, 173.1, 172.9, 170.9, 167.8, 158.1, 157.1, 152.7, 151.4, 149.1, 145.3, 143.8, 138.8, 130.4, 132.0, 130.1, 128.9 , 128.7, 127.6, 126.7, 124.1, 120.8, 120.6, 120.4, 120.2, 114.4, 112.0, 111.3, 110.4, 110.2, 104.7, 70.2, 69.7, 62.6, 60.9, 59.4, 57.9, 57.7, 56.5, 54.4, 49.2, 42.3, 37.5, 36.6, 36.3, 35.0, 33.1, 32.0, 31.5, 25.6, 17.3, 14.4. LC-MS (ES$^+$): m/z 522.44 $[M+2H]^{2+}$ and m/z 1043.14 $[M+H]^+$. ESI-HRMS (m/z): calcd. for $[C_{56}H_{66}N_8O_{10}S + H]^+$ 1043.4623; obsd. 1043.4617.

**HPP-2.** To a solution of **11b** (30 mg, 0.05 mmol) and **HPI-1-F** (26 mg, 0.05 mmol) in degassed THF/$H_2O$/MeOH (2.6 ml, 1:1:2) were added degassed aqueous solutions of 2 M sodium ascorbate (53 μL, 0.11 mmol) and 1 M $CuSO_4$ (11 μL, 0.01 mmol, 20 mol%) and the resulting mixture was stirred overnight at rt, after which it was concentrated under reduced pressure. $CH_2Cl_2$/MeOH (9:1) and $H_2O$ were added to the reaction mixture and after partitioning of the layers, the aqueous layer was extracted with $CH_2Cl_2$/MeOH (9:1). The organic layer was dried with $MgSO_4$ and concentrated. The resulting crude was purified by PTLC ($SiO_2$, $CH_2Cl_2$/MeOH 12:1, eluted once) to give **HPP-2** as an off-white solid (41 mg, 73%).; $R_f$= 0.37 ($CH_2Cl_2$/MeOH 10:1). $^1H$ NMR (500 MHz, $CD_3OD$): δ 8.76 (s, 1H), 7.93 (s, 1H), 7.38 – 7.28 (m, 4H), 7.14 – 6.56 (m, 8H), 5.07 – 4.89 (m, 3H), 4.51 (s, 1H), 4.49 – 4.35 (m, 3H), 4.29 (t, *J* = 7.0 Hz, 2H), 4.24 (dd, *J* = 15.6, 2.4 Hz, 1H), 4.09 – 3.99 (m, 2H), 3.79 (d, *J* = 11.4 Hz, 1H), 3.74 – 3.64 (m, 1H), 3.70 (s, 3H), 3.49 – 3.37 (m, 3H), 3.18 (s, 3H), 2.75 – 2.46 (m, 3H), 2.38 – 2.29 (m, 4H), 2.25 (s, 3H), 2.21 – 2.08 (m, 3H), 2.01 – 1.93 (m, 1H), 1.81 (p, *J* = 7.3 Hz, 2H), 1.53 (p, *J* = 7.4 Hz, 2H), 1.20 (m, 2H), 0.92 (s, 9H). $^{13}C$ NMR (126 MHz, $CD_3OD$) δ 197.2, 174.2, 173.1, 170.9, 167.8, 158.0, 157.1, 152.7, 152.1, 151., 149., 148.6, 147.6, 145.3, 143.7, 138.8, 132.0, 130.4, 130.1, 128.9 (2C), 128.7, 127.7, 127.6 (2C), 126.7, 123.8, 120.8, 120.6, 120.4, 120.2, 114.4, 112.1, 111.3, 110.4, 110.2, 104.8, 70.2, 69.7, 62.6, 60.9, 59.4, 57.6, 56.6, 54.4, 49.8, 42.3, 37.5, 36.3, 34.8, 33.1, 32.0, 29.5, 25.6 (3C), 17.3, 14.4. LC-MS ($ES^+$): m/z 536.45 $[M+2H]^{2+}$ and m/z 1071.11 $[M+H]^+$. ESI-HRMS (m/z): calcd. for $[C_{58}H_{70}N_8O_{10}S + H]^+$ 1071.4923; obsd. 1071.4918.

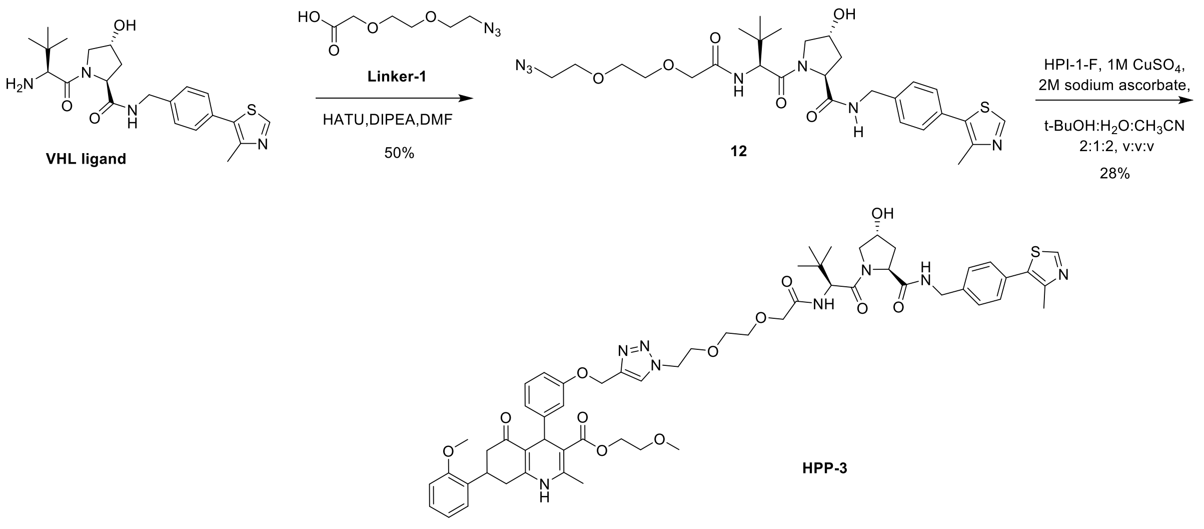


**Scheme S3.** Synthesis of **HPP-3.**

**Compound 12.** To a solution of **VHL ligand** (68 mg, 0.16 mmol) in anhydrous DMF (3.0 mL) were added DIPEA (40 μL, 0.22 mmol). To another round-bottom flask was added **Linker-1** (33 mg, 0.17 mmol), HBTU (10 mg, 0.26 mmol) followed by DIPEA (40 μL, 0.17 mmol) in anhydrous DMF (2.0 mL). Both mixtures were stirred for 10 min separately, then combined, and stirred for another 30 min at rt. Upon completion, the reaction mixture was diluted with $CH_2Cl_2$, washed with sat. $NH_4Cl$ aq. (×2). The first aqueous layer was re-extracted with $CH_2Cl_2$. The organic layer was dried over $MgSO_4$, concentrated.

The resulting crude was purified by PTLC ($SiO_2$, $CH_2Cl_2$/MeOH 15:1, eluted once) to give **12** as a yellow solid (51 mg, 50%).; $R_f$= 0.35 ($CH_2Cl_2$/MeOH 10:1). $^1$H NMR (400 MHz, $CDCl_3$): δ 8.76 (s, 1H), 7.83 – 7.22 (m, 4H), 4.65 (t, *J* = 7.6 Hz, 1H), 4.49 – 4.43 (m, 3H), 4.26 (d, *J* = 15.2 Hz, 1H), 4.01 (d, *J* = 11.7 Hz, 1H), 3.62 – 3.53 (m, 7H), 3.32 – 3.28 (m, 2H), 2.44 – 2.39 (m, 1H), 2.43 (s, 3H), 2.09 – 2.04 (m, 1H), 1.40 (t, *J* = 7.5 Hz, 2H), 0.88 (s, 9H). LC-MS ($ES^+$): m/z 601.84 $[M+H]^+$.

**HPP-3.** To a solution of **12** (21 mg, 0.035 mmol) and **HPI-1-F** (18 mg, 0.035 mmol) in degassed *t*-BuOH/$CH_3CN/H_2O$ (1.8 mL, 2:2:1) were added degassed aqueous solutions of 2 M sodium ascorbate (87 µL, 0.2 mmol) and 1 M $CuSO_4$ (7 µL, 0.01 mmol, 20 mol%) and the mixture was stirred overnight at rt. $CH_2Cl_2$/MeOH (9:1) and $H_2O$ were added to the reaction mixture and after partitioning of the layers, the aqueous layer was extracted with $CH_2Cl_2$/MeOH (9:1, ×2). The organic layer was dried over $MgSO_4$ and concentrated. The resulting crude was purified by reverse phase chromatography (BGB Scorpius C18 4.5 g, $H_2O$ + 0.1% TFA/$CH_3CN$ + 0.1% TFA 80:20 to 20:80) and silica gel column chromatography ($CH_2Cl_2$/MeOH 15:1) to give **HPP-3** as a yellow solid (11 mg, 28%).; $R_f$= 0.40 ($CH_2Cl_2$/MeOH 10:1). $^1$H NMR (500 MHz, $CD_3OD$): δ 8.8 (s, 1H), 8.11 (s, 1H), 7.44 – 7.31 (m, 4H), 7.25 – 6.65 (m, 8H), 5.16 – 5.00 (m, 3H), 4.70 (m, 1H), 4.65 – 4.43 (m, 5H), 4.34 – 4.27 (m, 1H), 4.13 (m, 2H), 4.03 – 3.74 (m, 6H), 4.78 (s, 3H), 3.66 – 3.46 (m, 7H), 3.28 (s, 3H), 2.86 – 2.52 (m, 4H), 2.44 (s, 3H), 2.35 (s, 3H), 2.25 – 2.15 (m, 1H), 2.07 (m, 1H), 1.02 (s, 9H). $^{13}$C NMR (126 MHz, $CD_3OD$): δ 197.1, 172.9, 170.7, 170.2, 167.8, 158.0, 157.1, 152.7, 152.1, 151.5, 149.1, 148.6, 147.6, 145.2, 138.7, 130.4, 130.1, 128.9 (2C), 128.1, 127.6 (2C), 126.6, 125.0, 120.6, 120.4, 114.5, 112.0, 111.9, 110.4, 104.7, 78.1, 70.9, 70.2, 69.9, 69.6, 69.2, 62.6, 60.8, 59.4, 57.7, 56.7, 54.4, 50.0, 42.3, 41.8, 39.9, 37.5, 36.6, 36.3, 35.8, 33.1, 32.0, 25.6(3C), 17.4, 14.5. LC-MS ($ES^+$): m/z 552.33 $[M+2H]^{2+}$ and m/z 110.13 $[M+H]^+$. ESI-HRMS (m/z): calcd. for $[C_{58}H_{70}N_8O_{12}S + H]^+$ 1103.4834; obsd. 1103.4886.

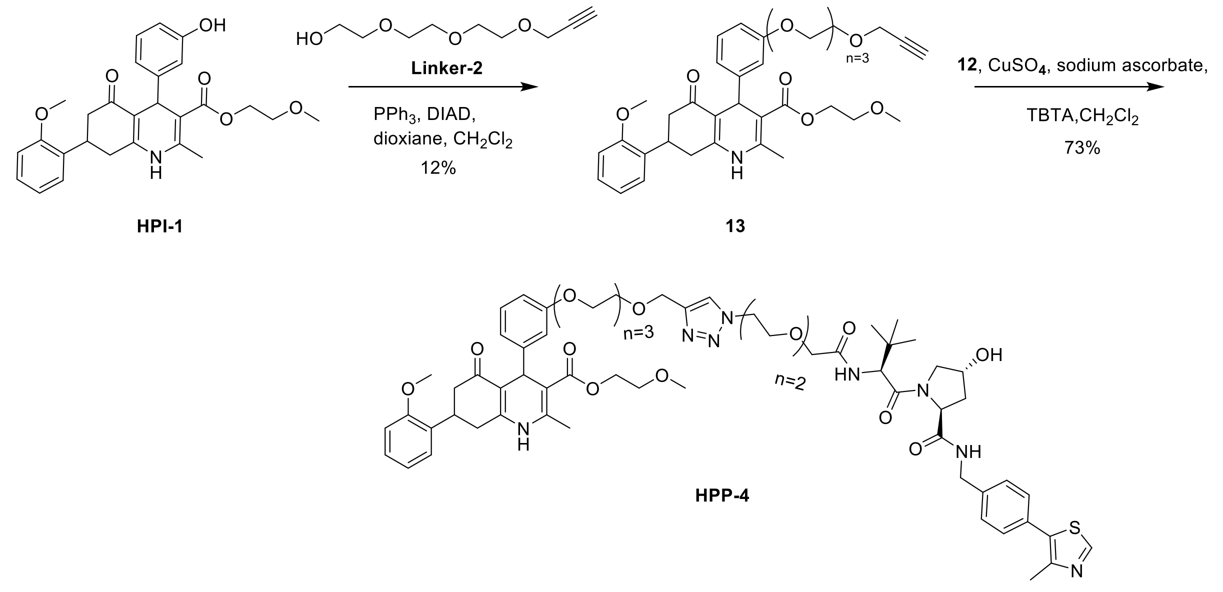


**Scheme S4.** Synthesis of **HPP-4**

**Compound 13.** To a solution of **HPI-1** (183 mg, 0.39 mmol) in anhydrous $CH_2Cl_2$/dioxane (4:1, 1.0 mL) were added $PPh_3$ (155 mg, 0.59) and **Linker-2** (74 mg, 0.39 mmol), and the solution was ice cooled. DIAD (0.12 mL, 0.59 mmol) was added at 0 °C slowly over 15 min. The mixture was stirred at 0 °C for 30 min and then at rt overnight. The mixture was concentrated *in vacuo*. The resulting crude was purified by silica gel column chromatography ($CH_2Cl_2$/MeOH 30:1) and reverse phase chromatography (BGB Scorpius C18 4.5 g, $H_2O$ + 0.1% TFA/$CH_3CN$ + 0.1% TFA gradient from 75:35 to 35:75) to give **13** as a pale-yellow solid (30 mg, 12%).; $R_f$= 0.22 ($CH_2Cl_2$/MeOH 30:1). $^1$H NMR (400 MHz, $CDCl_3$): δ 7.25 – 6.68 (m, 8H), 6.15 (br, 1H, NH), 5.14 (s, 1H), 4.19 (s, 2H), 4.17 (t, *J* = 4.8 Hz, 2H), 4.12 (t, *J* = 4.3 Hz, 2H), 3.83 (t, *J* = 4.7 Hz, 2H), 3.80 (s, 3H), 3.74 – 3.67 (m, 8H), 3.62 – 3.57 (m, 2H), 3.52 (t, *J* = 4.8 Hz, 2H), 3.31 (s, 3H), 2.76 – 2.56 (m, 4H), 2.41 (t, *J* = 2.3 Hz, 1H), 2.39 (s, 3H). $^{13}$C NMR (101 MHz, $CDCl_3$): δ 196.6, 167.2, 158.6, 157.2, 150.5, 148.4, 143.7, 130.2, 128.8, 128.1, 127.1, 121.1, 120.7, 114.8, 112.8, 112.0, 110.7, 106.0, 79.5, 74.7, 70.7, 70.6, 70.4, 70.4, 69.8, 69.1, 67.1, 62.9, 58.8, 58.4, 55.1, 41.9, 36.4, 33.1, 33.1, 19.5. LC-MS (ES$^+$): m/z 634.22 $[M+H]^+$.

**HPP-4.** To a solution of **13** (20 mg, 0.032 mmol), **12** (19 mg, 0.032 mmol) and TBTA (5 mg, 9 µmol) in $CH_2Cl_2$ (2.0 mL) was added a solution of sodium ascorbate (6 mg, 0.032 mmol) and $CuSO_4$ (5 mg, 0.032 mmol) in $H_2O$ (0.1 mL). The mixture was vigorously stirred overnight at rt. The mixture was diluted with $CH_2Cl_2$, dried over $MgSO_4$ and concentrated. The resulting crude was purified by PTLC ($SiO_2$, $CH_2Cl_2$/MeOH 10:1, eluted once) to give **HPP-4** as a white solid (41 mg, 73%).

; $R_f$= 0.35 ($CH_2Cl_2$/MeOH 10:1). $^1$H NMR (500 MHz, $CD_3OD$) δ 8.06 (s, 1H), 7.44 (s, 1H), 7.42 – 7.33 (m, 4H), 7.25 – 6.62 (m, 8H), 5.06 (s, 1H), 4.71 (s, 1H), 4.61 – 4.47 (m, 7H), 4.32 (d, *J* = 15.4 Hz, 1H), 4.13 (m, 2H), 4.10 – 4.05 (m, 2H), 4.02 – 3.94 (m, 2H), 3.91 – 3.72 (m, 10H), 3.69 – 3.57 (m, 12H), 3.53 (m,

2H), 2.85 – 2.54 (m, 3H), 2.49 – 2.38 (m, 4H), 2.35 (s, 3H), 2.24 (m, 1H), 2.08 (m, 1H), 1.03 (s, 9H). $^{13}$C NMR (126 MHz, $CD_3OD$) δ 197.2, 172.9, 170.6, 170.1, 167.8, 158.6, 157.1, 152.7, 149.0, 145.2, 138.8, 130.3, 128.9 (2C), 128.6, 127.6 (2C), 126.6, 120.4, 120.2, 114.3, 111.5, 110.4, 104.8, 70.9, 70.3, 70.2, 70.2, 70.1, 69.9, 69.7, 69.6, 69.5, 69.3, 69.2, 66.9, 63.7, 62.6, 59.4, 57.7, 56.6, 54.4, 49.9, 42.3, 37.6, 36.2, 35.9, 33.1, 32.0, 25.5 (3C), 17.3, 14.5. LC-MS (ES$^+$): m/z 618.54 $[M+2H]^{2+}$ and m/z 1235.16 $[M+H]^+$. ESI-HRMS (m/z): calcd. for $[C_{64}H_{82}N_8O_{10}S + H]^+$ 1235.5620; obsd. 1235.5629.

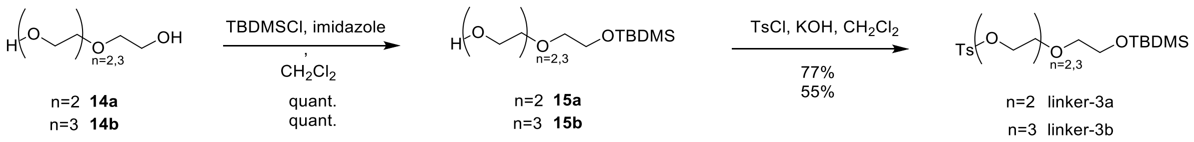


**Scheme S5.** Synthesis of **linker-3a** and **3b**

**Compound 15a**. To a solution TBDMSCl (1.21 g, 8.0 mmol) and imidazole (907 mg, 13.3 mmol) in anhydrous DMF (15 mL) was added triethylene glycol **14a** (2.0 g, 13.3 mmol). The reaction mixture was stirred at rt for 4 h. The reaction mixture was concentrated and co-evaporated with toluene. The resulting crude was purified by silica gel column chromatography (petroleum ether/EtOAc 1:1) to give **14a** as a colorless oil (0.60 g, 29%).; $R_f$= 0.40 (pentane/EtOAc 1:1). $^1$H NMR (400 MHz, $CDCl_3$): δ 3.76 (t, *J* = 5.2 Hz, 2H), 3.73 – 3.70 (m, 2H), 3.66 (s, 4H), 3.61 – 3.59 (m, 2H), 3.56 (t, *J* = 5.2 Hz, 2H), 2.46 (br, 1H, OH), 0.89 (s, 9H), 0.06 (s, 6H). LC-MS (ES$^+$): m/z 264.94 $[M+H]^+$.

**Compound 15b.** Tetraethylene glycol **14b** (4.51 g, 23.2 mmol) and imidazole (379 mg, 5.6 mmol) were dissolved in 3 mL anhydrous $CH_2Cl_2$. The reaction mixture was cooled down to 0 °C and a solution of TBDMSCl (0.80 ml, 4.64 mmol) in anhydrous $CH_2Cl_2$ (7.0 mL) was added dropwise over 2 h. The reaction mixture was stirred at rt overnight. $H_2O$ (10 mL) was added to the mixture and the organic layer was washed thrice with $H_2O$ to remove the excess tetraethylene glycol. The organic layer was dried with $MgSO_4$ and concentrated. Compound **15b** was obtained as a colorless oil (1.46 g, quant.), which was used directly for the next step without further purification.; $R_f$= 0.27 (pentane/EtOAc 1:1). $^1$H NMR (400 MHz, $CDCl_3$): δ 3.76 (t, *J* = 5.4 Hz, 2H), 3.73 – 3.70 (m, 2H), 3.68 – 3.64 (m, 8H), 3.63 – 3.60 (m, 2H), 3.56 (t, *J* = 5.3 Hz, 2H), 0.89 (s, 9H), 0.06 (s, 6H). LC-MS (ES$^+$): m/z 309.01 $[M+H]^+$.

**Linker-3a.** To a solution of **15a** (600 mg, 2.30 mmol) in anhydrous $CH_2Cl_2$ (10.0 mL) was added anhydrous KOH (510 mg, 9.10 mmol). The reaction mixture was stirred for 15 min at 0 °C before TsCl (0.52 g, 2.70 mmol) was added carefully, and stirred for 3 h. A solution of cold $H_2O$ and $CH_2Cl_2$ was added into the reaction mixture. The aqueous layer was extracted with $CH_2Cl_2$ (×2). The organic layer was washed with cold $H_2O$, dried over $MgSO_4$ and concentrated. The resulting crude was purified by silica gel column chromatography (pentane/EtOAc 7:1) to give **Linker-3a** as a colorless oil (0.74 g, 77 %).; $R_f$= 0.29 (pentane/EtOAc 7:1). $^1$H NMR (400 MHz, $CDCl_3$): δ 7.80 (d, $J$ = 8.4 Hz, 2H), 7.34 (d, $J$ = 7.9 Hz, 2H), 4.16 (t, $J$ = 4.6 Hz, 2H), 3.74 (t, $J$ = 5.6 Hz, 2H), 3.69 (t, $J$ = 4.9 Hz, 2H), 3.59 – 3.56 (m, 4H), 3.52 (t, $J$ = 5.0 Hz, 2H), 2.44 (s, 3H), 0.88 (s, 9H), 0.05 (s, 6H). LC-MS ($ES^+$): m/z 418.91 $[M+H]^+$.

**Linker-3b. 15b** (1.46 g, 4.74 mmol) was reacted according to the procedure of **15a** described above to yield **Linker-3b** as a colorless oil (1.21 g, 55 %).; $R_f$= 0.50 (pentane/EtOAc 2:1). $^1$H NMR (400 MHz, $CDCl_3$): δ 7.80 (d, $J$ = 8.3 Hz, 2H), 7.33 (d, $J$ = 8.3 Hz, 2H), 4.16 (t, $J$ = 5.0 Hz, 2H), 3.75 (t, $J$ = 5.5 Hz, 2H), 3.68 (t, $J$ = 5.0 Hz, 2H), 3.65 – 3.59 (m, 4H), 3.58 (s, 4H), 3.54 (t, $J$ = 5.6 Hz, 2H), 2.44 (s, 3H), 0.89 (s, 9H), 0.06 (s, 6H). LC-MS ($ES^+$): m/z 462.86 $[M+H]^+$.

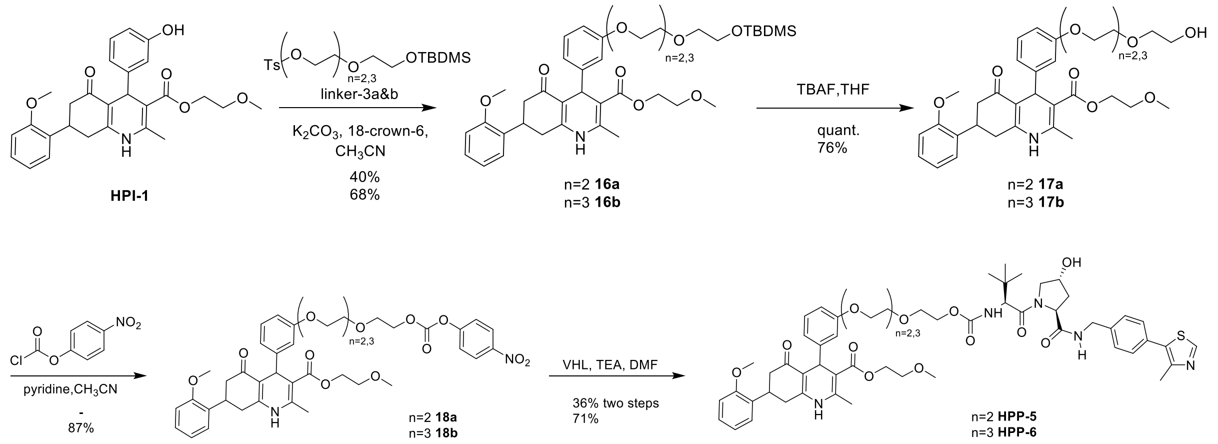


**Scheme S6.** Synthesis of **HPP-5** and **HPP-6**

**Compound 16a.** To a solution of **HPI-1** (240 mg, 0.52 mmol) in anhydrous $CH_3CN$ (20 mL) were added **Linker-3a** (180 mg, 0.43 mmol), $K_2CO_3$ (360 mg, 2.60 mmol) and 18-Crown-6 (11 mg, 40 µmol). The reaction mixture was refluxed overnight, after which it was cooled to rt and filtered. The filtrate was concentrated *in vacuo*. To remove the remaining salt, the acquired oil was re-dissolved in EtOAc, washed with $H_2O$ (×2) and brine, dried over $MgSO_4$ and concentrated. The resulting crude was purified by silica gel column chromatography (pentane/EtOAc 1:1 to 1:2) to give **16a** as a yellow oil (115 mg, 40 %).; $R_f$= 0.36 (pentane/EtOAc 1:1). $^1H$ NMR (400 MHz, $CDCl_3$): δ 7.23 (d, *J* = 8.0 Hz, 1H), 7.14 (t, *J* = 8.0 Hz, 2H), 7.01 (d, *J* = 7.3 Hz, 1H), 6.94 (t, *J* = 7.0 Hz, 2H), 6.88 (d, *J* = 8.6 Hz, 1H), 6.69 (d, *J* = 8.0 Hz, 1H), 5.81 (br, 1H, NH), 5.15 (s, 1H), 4.16 (t, *J* = 5.5 Hz, 2H), 4.12 (t, *J* = 4.5 Hz, 2H), 3.84 (t, *J* = 4.5 Hz, 2H), 3.81 (s, 3H), 3.77 (t, *J* = 5.5 Hz, 2H), 3.71 – 3.68 (m, 4H), 3.65 – 3.61 (m, 1H), 3.57 (t, *J* = 5.5 Hz, 2H), 3.52 (t, *J* = 4.5 Hz, 2H), 3.31 (s, 3H), 2.76 – 2.53 (m, 4H), 2.40 (s, 3H), 0.89 (s, 9H), 0.06 (s, 6H). LC-MS ($ES^+$): m/z 710.30 $[M+H]^+$.

**Compound 16b.** To a solution of **HPI-1** (190 mg, 0.42 mmol) in anhydrous $CH_3CN$ (8.0 mL) were added **Linker-3b** (240 mg, 0.52 mmol), $K_2CO_3$ (430 mg, 3.12 mmol), and 18-Crown-6 (14 mg, 50 µmol). The reaction mixture was refluxed overnight, after which it was cooled to room temperature and poured into EtOAc, washed with $H_2O$ (×2) and brine, dried over $MgSO_4$ and concentrated. The resulting crude was purified by silica gel column chromatography ($CH_2Cl_2$/MeOH 10:0 to 10:1) and another silica gel column chromatography (Pentane/EtOAc 1:1 to 1:2) to give **16b** as a yellow oil (0.265 g, 68%).; $R_f$= 0.22 (pentane/EtOAc 1:1). $^1H$ NMR (400 MHz, $CDCl_3$): δ 7.23 (d, *J* = 8.0 Hz, 1H), 7.13 (t, *J* = 8.0 Hz, 2H), 7.01 (d, *J* = 7.4 Hz, 1H), 6.93 (t, *J* = 7.5 Hz, 2H), 6.86 (d, *J* = 8.0 Hz, 1H), 6.68 (d, *J* = 7.1 Hz, 1H), 6.15 (br, 1H, NH), 5.14 (s, 1H), 4.16 (t, *J* = 5.5 Hz, 2H), 4.11 (t, *J* = 5.5 Hz, 2H), 3.82 (t, *J* = 5.1 Hz, 2H), 3.79 (s, 3H), 3.76 (t, *J* = 5.5 Hz, 2H), 3.72 – 3.67 (m, 5H), 3.65 (s, 4H), 3.55 (t, *J* = 5.3 Hz, 2H), 3.51 (t, *J* = 5.0 Hz, 2H), 3.30 (s, 3H), 2.77 – 2.53 (m, 4H), 2.38 (s, 3H), 0.88 (s, 9H), 0.06 (s, 6H). LC-MS ($ES^+$): m/z 754.36 $[M+H]^+$.

**Compound 17a.** To a solution of **16a** (120 mg, 0.16 mmol) in anhydrous THF (3.0 mL) was added 1.0 M TBAF in THF (90 µl, 0.32 mmol) and the mixture was stirred at 40 °C overnight. The reaction mixture was concentrated, and the resulting oil was re-dissolved in EtOAc. The organic layer was washed with water (×8), sat. $NH_4Cl$ aq. (×3) and brine, dried over $MgSO_4$ and concentrated. Compound **17a** was obtained as a pale-yellow oil (102 mg, quant.), which was used directly for the next step without further purification.; $R_f$= 0.35 ($CH_2Cl_2$/MeOH 15:1). $^1H$ NMR (400 MHz, $CDCl_3$): δ 7.25 – 6.68 (m, 8H), 6.02 (br, 1H, NH), 5.15 (s, 1H), 4.18 – 4.11 (m, 4H), 3.85 (t, *J* = 5.0 Hz, 2H), 3.80 (s, 3H), 3.74 – 3.68 (m, 6H), 3.65 – 3.58 (m, 1H), 3.61 (t, *J* = 4.2 Hz, 2H), 3.52 (t, *J* = 5.0 Hz, 2H), 3.30 (s, 3H), 2.79 – 2.53 (m, 4H), 2.40 (s, 3H). LC-MS ($ES^+$): m/z 596.15 $[M+H]^+$.

**Compound 17b. 16b** (0.22 g, 0.29 mmol) was reacted according to the procedure of **16a** described above. The resulting crude was purified by silica gel column chromatography (EtOAc 100% to $CH_2Cl_2$/MeOH 10:1) to give **17b** as a yellow oil (140 mg, 76%).; $R_f$= 0.27 ($CH_2Cl_2$/MeOH 10:1). $^1H$ NMR (400 MHz, $CDCl_3$): δ 7.23 – 6.66 (m, 8H), 5.13 (s, 1H), 4.17 (t, *J* = 5.0 Hz, 2H), 4.13 – 4.10 (m, 2H), 5.56

– 3.79 (m, 2H), 3.80 (s, 3H), 3.72 – 3.66 (m, 10H), 3.60 – 3.58 (m, 3H), 3.52 (t, *J* = 5.0 Hz, 2H), 3.31 (s, 3H), 2.81 – 2.58 (m, 4H), 2.41 (s, 3H). LC-MS ($ES^+$): m/z 640.22 $[M+H]^+$.

**HPP-5.** A stirred mixture of 4-nitrophenyl chloroformate (51 mg, 0.25 mmol) and pyridine (30 µL, 0.34 mmol) in $CH_3CN$ (2.0 mL) was cooled to 0 °C for 15 min. A solution of **17a** (100 mg, 0.17 mmol) in anhydrous $CH_3CN$ (2.0 mL) was added slowly. The reaction mixture was stirred at rt overnight, after which it was concentrated *in vacuo*. The resulting crude was re-dissolved in $CH_2Cl_2$ and washed with brine. The organic layer was dried over $MgSO_4$ and concentrated to give crude **18a.** LC-MS ($ES^+$): m/z 761.18 $[M+H]^+$. The crude containing **18a** was re-dissolved in anhydrous DMF (20 mL). **VHL** (40 mg, 92.1 µmol) and $Et_3N$ (50 µL, excess) were added, and stirred at rt overnight. The reaction mixture was concentrated and purified by PTLC ($SiO_2$, $CH_2Cl_2$/MeOH 17:1, eluted twice) to give **HPP-5** as a yellow solid (65 mg, 36% two steps).; $R_f$= 0.39 ($CH_2Cl_2$/MeOH 15:1). $^1H$ NMR (500 MHz, $CD_3OD$): δ 8.87 (s, 1H), 7.47 – 7.37 (m, 4H), 7.24 – 6.67 (m, 8H), 5.06 (s, 1H), 4.57 (t, *J* = 8.3 Hz, 1H), 4.52 (m, 2H), 4.34 (d, *J* = 13.4 Hz, 2H), 4.20 – 4.06 (m, 6H), 3.88 (d, *J* = 11.1 Hz, 1H), 3.81 (d, *J* = 7.5 Hz, 6H), 3.71 – 3.62 (m, 6H), 3.59 – 3.48 (m, 3H), 3.30 (s, 3H), 2.86 – 2.50 (m, 3H), 2.47 (s, 4H), 2.36 (s, 3H), 1.01 (s, 9H). $^{13}C$ NMR (126 MHz, $CD_3OD$): δ 197.2, 173.0, 171.2, 167.8, 158.6, 157.1, 152.6, 151.4, 148.9, 147.5, 145.2, 138.8, 132.0, 130.4, 130.0, 128.9 (2C), 128.5, 127.7, 127.6 (2C), 126.6, 120.4, 120.2, 114.3, 111.5, 111.4, 110.4, 104.8, 70.3, 70.2, 70.1, 69.6, 69.5, 69.1, 66.9, 64.0, 62.6, 59.5, 59.4, 57.7, 56.5, 54.4, 47.8, 42.3, 42.2, 37.5, 36.2, 35.2, 33.1, 32.0, 25.5 (3C), 17.3, 14.4. LC-MS ($ES^+$): m/z 527.03 $[M+2H]^{2+}$ and m/z 1052.11 $[M+H]^+$. ESI-HRMS (m/z): calcd. for $[C_{56}H_{69}N_5O_{13}S + H]^+$ 1052.4613; obsd. 1052.4646.

**HPP-6.** 4-Nitrophenyl chloroformate (88 mg, 0.44 mmol) was reacted with **17b** (0.140 g, 0.22 mmol) according to the procedure of **17a** described above. The resulting crude was purified by silica gel column chromatography (pentane/EtOAc 1:4 to 100 % EtOAc) to give **18b** (0.153 g, 87%); $R_f$ = 0.36 (pentane/EtOAc 1:4). $^1H$ NMR (400 MHz, $CDCl_3$): δ 8.26 (d, *J* = 9.1 Hz, 2H). 7.37 (d, *J* = 9.1 Hz, 2H), 7.24 – 6.64 (m, 8H), 5.90 (br, 1H, NH), 5.14 (s, 1H), 4.42 (t, *J* = 4.6 Hz, 2H), 4.16 (t, *J* = 4.7 Hz, 2H), 4.13 – 4.11 (m, 2H), 3.84 (t, *J* = 5.0 Hz, 2H), 3.81 – 3.79 (m, 2H), 3.73 – 3.68 (m, 8H), 3.64 – 3.57 (m, 4H), 3.51 (t, *J* = 5.0 Hz, 2H), 3.31 (s, 3H), 2.79 – 2.52 (m, 4H), 2.39 (s, 3H). LC-MS ($ES^+$): m/z 805.24 $[M+H]^+$. **18b** (145 mg, 0.18 mmol) was re-dissolved in anhydrous DMF (5.0 mL). **VHL ligand** (109 mg, 0.25 mmol) and $Et_3N$ (130 µL, excess) were added, and stirred overnight at room temperature. The reaction mixture was concentrated and purified by PTLC ($SiO_2$, $CH_2Cl_2$/MeOH 13:1 and 15:1, eluted once, respectively) to give **HPP-6** as a yellow solid (140 mg, 71%).; $R_f$= 0.28 ($CH_2Cl_2$/MeOH 13:1). $^1H$ NMR (500 MHz, $CD_3OD$): δ 8.74 (s, 1H), 7.35 – 7.26 (m, 4H), 7.12 – 6.55 (m, 8H), 4.96 (s, 1H), 4.48 (t, *J* = 8.3 Hz, 1H), 4.44 – 4.35 (m, 2H), 4.27 – 4.20 (m, 2H), 4.10 – 3.92 (m, 6H), 3.77 (d, *J* = 11.0 Hz, 1H), 3.74 – 3.66 (m,

6H), 3.58 – 3.48 (m, 10H), 3.46 – 3.39 (m, 3H), 3.19 (s, 3H), 2.71 – 2.45 (m, 3H), 2.35 (s, 4H), 2.25 (s, 3H), 2.15 – 2.06 (m, 1H), 1.97 (m, 1H), 0.91 (s, 9H). $^{13}$C NMR (126 MHz, $CD_3OD$): δ 197.2, 173.0, 171.2, 167.8, 158.6, 157.1, 152.6, 151.4, 149.0, 147.6, 145.2, 138.8, 132.0, 130.3, 130.1, 128.9 (2C), 128.6, 127.7, 127.5 (2C), 126.8, 126.6, 120.4, 120.2, 114.3, 111.5, 110.4, 104.8, 78.1, 70.3, 70.2, 70.2, 70.1, 70.1, 69.7, 69.5, 69.0, 66.9, 64.0, 62.6, 59.5, 59.4, 57.7, 56.6, 54.4, 42.3, 37.5, 36.5, 36.2, 35.3, 33.1, 32.0, 25.6 (3C), 17.4, 14.5. LC-MS (ES$^+$): m/z 548.75 $[M+2H]^{2+}$ and m/z 1096.42 $[M+H]^+$. ESI-HRMS (m/z): calcd. for $[C_{58}H_{73}N_5O_{14}S + H]^+$ 1096.4875; obsd. 1096.4941.

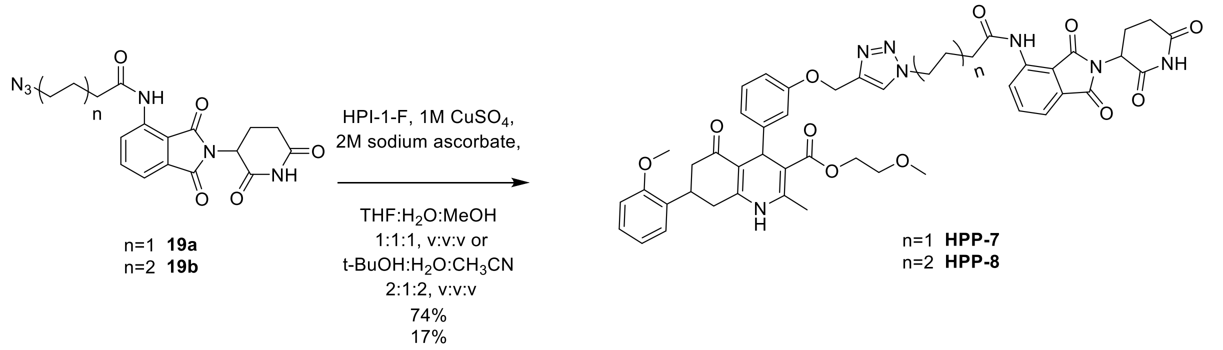


**Scheme S7.** Synthesis of **HPP-7** and **HPP-8**

**Compound 19a 19b were prepared following previous reported procedures[8].**

**HPP-7.** To a solution of **19a** (20 mg, 0.052 mmol) and **HPI-1-F** (26 mg, 0.052 mmol) in degassed THF/$H_2O$/MeOH (3.6 ml, 1:1:1) were added degassed aqueous solutions of 2 M sodium ascorbate (52 µL, 0.10 mmol) and 1 M $CuSO_4$ (10 µL, 0.01 mmol, 20 mol%). The mixture was stirred at rt overnight, after which it was concentrated under reduced pressure. The reaction mixture washed with $H_2O$ and extracted with $CH_2Cl_2$. The aqueous layer was re-extracted with $CH_2Cl_2$ (×6). The combined organic layers were dried over $MgSO_4$ and concentrated. The resulting crude was purified by PTLC ($SiO_2$, $CH_2Cl_2$/MeOH 13:1, eluted once) to give **HPP-7** as an off-white solid (34 mg, 74%).; $R_f$= 0.28 ($CH_2Cl_2$/MeOH 13:1). $^1$H NMR (500 MHz, $CDCl_3$): δ 9.39 (br, 1H, NH), 8.76 (d, *J* = 8.5 Hz, 1H), 8.19 (d, *J* = 16.4 Hz, 1H, NH), 7.74 (s, 1H), 7.72 – 7.66 (m, 1H), 7.55 (d, *J* = 7.3 Hz, 1H), 7.25 – 6.67 (m, 8H), 5.99 (br, 1H, NH), 5.26 – 5.07 (m, 3H), 4.94 (m, 1H), 4.49 (t, *J* = 6.6 Hz, 2H), 4.26 – 4.10 (m, 2H), 3.79 (s, 3H), 3.68 – 3.55 (m, 1H), 3.52 (t, *J* = 4.9 Hz, 2H), 3.29 (s, 3H), 2.95 – 2.53 (m, 7H), 2.50 (t, *J* = 7.0 Hz, 2H), 2.38 – 2.32 (m, 4H), 2.19 – 2.09 (m, 1H). $^{13}$C NMR (126 MHz, $CDCl_3$): δ 193.3, 168.2, 166.6, 165.4, 165.4, 164.9, 164.3, 155.6, 154.8, 147.0, 146.4, 142.2, 141.5, 141.5, 135.1, 134.1, 128.8, 128.1, 126.6, 125.7, 124.7, 122.9, 120.9, 119.0, 118.9, 118.4, 116.4, 112.3, 110.5, 110.2, 108.3, 68.1, 60.6, 59.4, 56.5, 52.8, 46.9, 46.8, 40.0, 34.2, 34.2, 31.4, 30.8, 29.0, 23.0, 20.3, 17.2. LC-MS (ES$^+$): m/z 886.15 $[M+H]^+$. ESI-HRMS (m/z): calcd. for $[C_{47}H_{47}N_7O_{11} + H]^+$ 886.3334; obsd. 886.3428.

**HPP-8.** To a solution of compound **19b** (13 mg, 0.032 mmol) and **HPI-1-F** (16 mg, 0.032 mmol) in degassed *t*-BuOH/$CH_3CN/H_2O$ (1.6 ml, 2:2:1) were added degassed aqueous solutions of 2 M sodium ascorbate (160 µL, 0.32 mmol) and 1 M $CuSO_4$ (16 µL, 0.02 mmol, 50 mol%) and the mixture was stirred at rt overnight. $CH_2Cl_2$ and $H_2O$ were added to the reaction mixture and after partitioning of the layers, the aqueous layer was extracted with $CH_2Cl_2$. The organic layer was dried over $MgSO_4$ and concentrated. The resulting crude was purified by silica gel column chromatography ($CH_2Cl_2$/MeOH 50:1 to 20:1) to give **HPP-8** as a white solid (5 mg, 17%).; $R_f$= 0.23 ($CH_2Cl_2$/MeOH 20:1). $^1$H NMR (500 MHz, $CDCl_3$): δ 9.40 (br, 1H, NH), 8.79 (d, *J* = 8.5 Hz, 1H), 8.36 (d, *J* = 11.6 Hz, 1H, NH), 7.77 – 7.61 (m, 2H), 7.53 (d, *J* = 7.2 Hz, 1H), 7.24 – 6.69 (m, 8H), 6.12 (br, 1H, NH), 5.25 – 5.06 (m, 3H), 4.99 – 4.87 (m, 1H), 4.36 (t, *J* = 6.9 Hz, 2H), 4.16 (t, *J* = 4.6, 2H), 3.79 (s, 3H), 3.65 – 3.54 (m, 1H), 3.54 – 3.49 (m, 2H), 3.29 (s, 3H), 2.91 – 2.51 (m, 7H), 2.45 (dd, *J* = 8.3, 6.6 Hz, 2H), 2.35 (s, 3H), 2.21 – 2.11 (m, 1H), 2.00 – 1.90 (m, 2H), 1.78 (m, 2H), 1.43 - 1.38 (m, 2H). $^{13}$C NMR (126 MHz, $CDCl_3$): δ 195.6, 171.8, 170.7, 169.1, 167.3, 166.7, 158.0, 157.1, 149.5, 148.8, 144.0, 137.7, 136.5, 131.1, 130.4, 128.9, 128.0, 127.1, 127.0, 125.2, 121.4, 121.3, 120.7, 120.6, 118.5, 115.4, 114.7, 112.4(2C), 110.7, 105.6, 70.5, 62.9, 61.8, 58.8, 55.2, 50.1, 49.3, 42.4, 37.4, 36.6, 33.2, 33.1, 31.3, 29.9, 25.9, 24.4, 22.7, 19.5. LC-MS (ES$^+$): m/z 457.82 $[M+2H]^{2+}$ and m/z 914.16 $[M+H]^+$. ESI-HRMS (m/z): calcd. for $[C_{49}H_{51}N_7O_{11} + H]^+$ 914.3647;obsd. 914.3668.

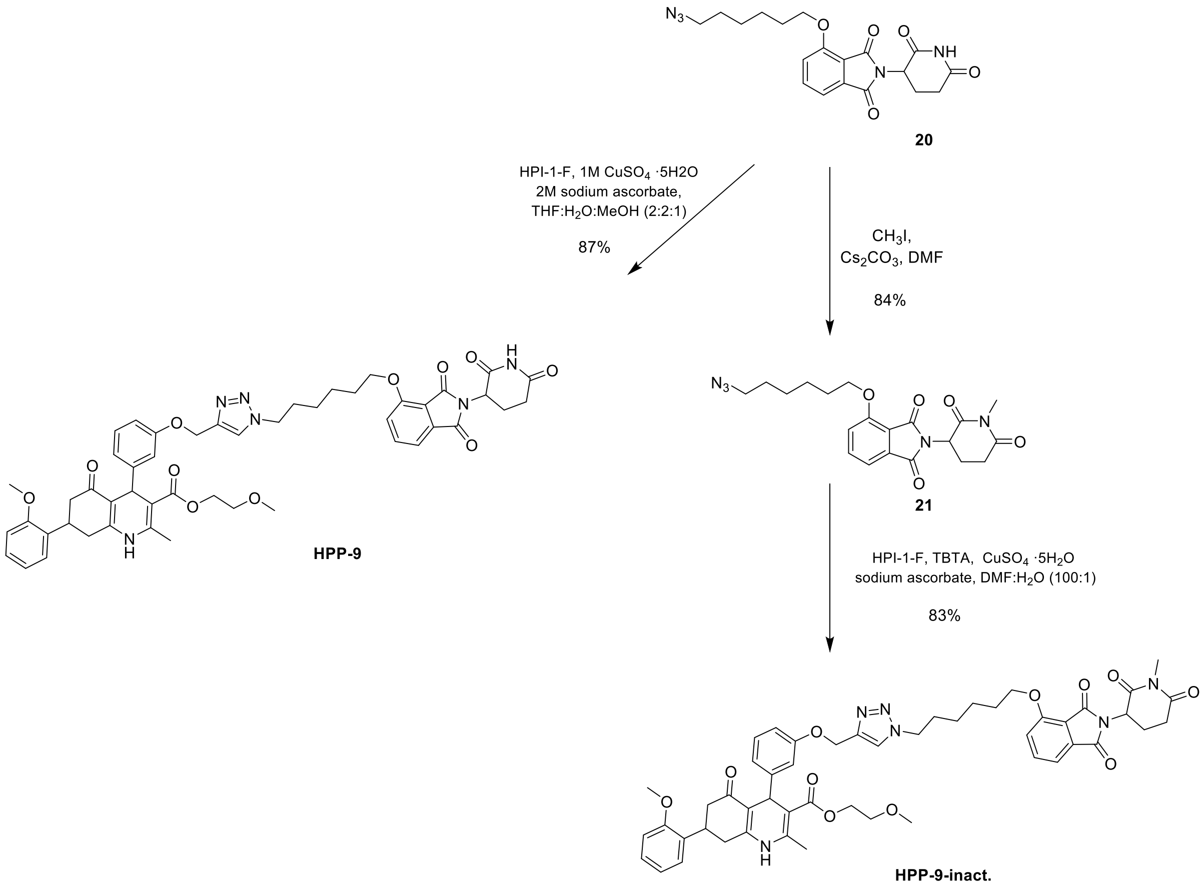


**Scheme S8.** Synthesis of **HPP-9** and **HPP-9-inact.**

Compound 20 was prepared following previous reported procedures [5].

**HPP-9.** To a solution of **20** (23 mg, 0.058 mmol) and **HPI-1-F** (23 mg, 0.046 mmol) in degassed THF/$H_2O$/MeOH (3.0 ml, 1:1:1) were added degassed aqueous solutions of 2 M sodium ascorbate (46 µL, 0.092 mmol) and 1 M $CuSO_4$ (9 µL, 0.009 mmol, 20 mol%). The mixture was stirred overnight at rt, after which it was concentrated under reduced pressure. $CH_2Cl_2$ and $H_2O$ were added to the reaction mixture and after partitioning of the layers, the aqueous layer was extracted with $CH_2Cl_2$ (×6). The organic layer was dried over $MgSO_4$ and concentrated. The resulting crude was purified by PTLC ($SiO_2$, $CH_2Cl_2$/MeOH 13:1, eluted once) to give **HPP-9** as a white solid (36 mg, 87%).; $R_f$= 0.49 ($CH_2Cl_2$/MeOH 10:1). $^1$H NMR (500 MHz, $CDCl_3$): δ 8.28 (br, 1H, NH), 7.72 (d, *J* = 7.9 Hz, 1H), 7.66 (dd, *J* = 8.2, 7.2 Hz, 1H), 7.44 (dd, *J* = 7.4, 1.9 Hz, 1H), 7.26 – 6.69 (m, 9H), 6.12 (br, 1H, NH), 5.24 – 5.07 (m, 3H), 4.93 (dd, *J* = 12.4, 5.4 Hz, 1H), 4.42 – 4.31 (m, 2H), 4.23 – 4.09 (m, 4H), 3.79 (s, 3H), 3.66 – 3.52 (m, 1H), 3.52 (t, *J* = 5.0 Hz, 2H), 3.29 (s, 3H), 2.90 – 2.51 (m, 7H), 2.36 (s, 3H), 2.15 – 2.07 (m, 1H), 2.01 – 1.90 (m, 2H), 1.85 (m, 2H), 1.57 (p, *J* = 7.5 Hz, 2H), 1.47 – 1.37 (m, 2H). $^{13}$C NMR (126 MHz, $CDCl_3$): δ 195.6, 170.9,

168.2, 167.3, 167.1, 165.8, 158.0, 157.2, 156.9, 156.6, 149.5, 148.8, 144.2, 144.0, 136.6, 133.8, 130.5, 128.9, 128.0, 127.1, 123.1, 121.4, 120.8, 118.9, 117.2, 115.9, 114.7, 112.8, 112.5, 110.7, 105.6, 70.5, 69.1, 62.9, 61.8, 58.9, 55.2, 50.2, 49.1, 42.5, 36.6, 33.2, 31.4, 29.9, 28.4, 25.9, 25.3, 22.6, 19.5. LC-MS ($ES^+$): m/z 451.35 $[M+2H]^{2+}$ and m/z 901.14 $[M+H]^+$. ESI-HRMS (m/z): calcd. for $[C_{50}H_{54}N_6O_{11} + H]^+$ 901.3694; obsd. 901.3746.

**Compound 21.** To a solution of **20** (30 mg, 0.072 mmol) in DMF (1mL) was added $Cs_2CO_3$ (40 mg, 0.13 mmol) and $CH_3I$ (0.01 mL, 0.11 mmol) at rt. The reaction mixture was stirred for 2h at the same temperature and additional $CH_3I$ was added and let it stir for another 2 hours. Then the mixture was diluted with EtOAc and quenched with HCl 1N. Then washed 3 times with $H_2O$ and one with brine, dried with $MgSO_4$ and concentrated in vacuo. Purification by PTLC with 30:1 $CH_2Cl_2$:MeOH (25 mg, 84%). ($R_F$: 0.78 in $CH_2Cl_2$:MeOH 30:1). $^1$H NMR (400 MHz, $CDCl_3$) δ 7.66 (dd, *J* = 8.5, 7.3 Hz, 1H), 7.44 (dd, *J* = 7.3, 0.7 Hz, 1H), 7.20 (dd, *J* = 8.5, 0.7 Hz, 1H), 5.00 – 4.91 (m, 1H), 4.18 (t, *J* = 6.4 Hz, 2H), 3.29 (t, *J* = 6.9 Hz, 2H), 3.20 (s, 3H), 3.02 – 2.67 (m, 3H), 2.13 – 2.05 (m, 1H), 1.95 – 1.85 (m, 2H), 1.69 – 1.42 (m, 6H).

**HPP-9 inact**. To a solution of HPI-1-F (18 mg, 36 µmol), **21** (15 mg, 36 µmol) and TBTA (5.7 mg, 10.5 µmol) in DMF (1 mL), a solution of $CuSO_4$ (5.7 mg, 36 µmol) and sodium ascorbate (4.8mg, 24 µmol) in $H_2O$ was added. The mixture was vigorously stirred at rt for 2h. The crude was purified by reverse phase chromatography (BGB Scorpius C18 4.5 g, $H_2O$ + 0.1% TFA/$CH_3CN$ + 0.1% TFA 95:5 to 10:90) to give **HPP-9 inactive** as a pale-yellow powder (27 mg, 83%). $^1$H NMR (500 MHz, DMSO) δ 9.16 (d, *J* = 53.2 Hz, 2H), 8.21 (d, *J* = 9.5 Hz, 1H), 7.80 (dd, *J* = 8.5, 7.3 Hz, 1H), 7.50 (d, *J* = 8.5 Hz, 1H), 7.44 (d, *J* = 7.2 Hz, 1H), 7.29 – 7.20 (m, 2H), 7.15 (t, *J* = 7.8 Hz, 1H), 6.99 – 6.91 (m, 2H), 6.84 – 6.72 (m, 3H), 5.14 (dd, *J* = 13.1, 5.4 Hz, 1H), 5.06 (d, *J* = 2.3 Hz, 1H), 4.91 (d, *J* = 21.4 Hz, 1H), 4.36 (t, *J* = 7.1 Hz, 2H), 4.18 (t, *J* = 6.4 Hz, 2H), 4.12 – 4.04 (m, 2H), 3.77 (d, *J* = 13.2 Hz, 3H), 3.51 – 3.41 (m, 3H), 3.21 (d, *J* = 5.1 Hz, 2H), 3.00 (s, 3H), 2.97 – 2.89 (m, 1H), 2.81 – 2.71 (m, 2H), 2.67 – 2.53 (m, 3H), 2.46 (p, *J* = 1.9 Hz, 1H), 2.28 (d, *J* = 11.4 Hz, 3H), 2.08 – 1.99 (m, 1H), 1.85 (p, *J* = 7.3 Hz, 2H), 1.74 (p, *J* = 6.6 Hz, 2H), 1.52 – 1.44 (m, 2H), 1.32 (tt, *J* = 9.8, 6.3 Hz, 2H). $^{13}$C NMR (126 MHz, DMSO) δ 194.4, 171.8, 169.7, 166.8, 165.3, 157.8, 157.7, 156.6, 156.0, 151.1, 149.2, 145.4, 142.7, 137.1, 133.2, 130.6, 129.0, 128.6, 127.8, 127.6, 126.9, 126.8, 124.3, 120.5, 120.3, 120.1, 119.8, 116.2, 115.2, 114.3, 111.3, 110.9, 110.7, 110.5, 103.3, 70.0, 68.7, 62.3, 60.9, 60.8, 58.0, 55.4, 55.3, 49.3, 42.6, 42.0, 35.9, 35.6, 32.1, 31.9, 31.1, 29.6, 28.1, 26.5, 25.4, 24.6, 21.2, 18.3. ESI-HRMS (m/z): calcd. for $[C_{51}H_{56}N_6O_{11} + H]+$ 915.3851; obsd. 915.3938.

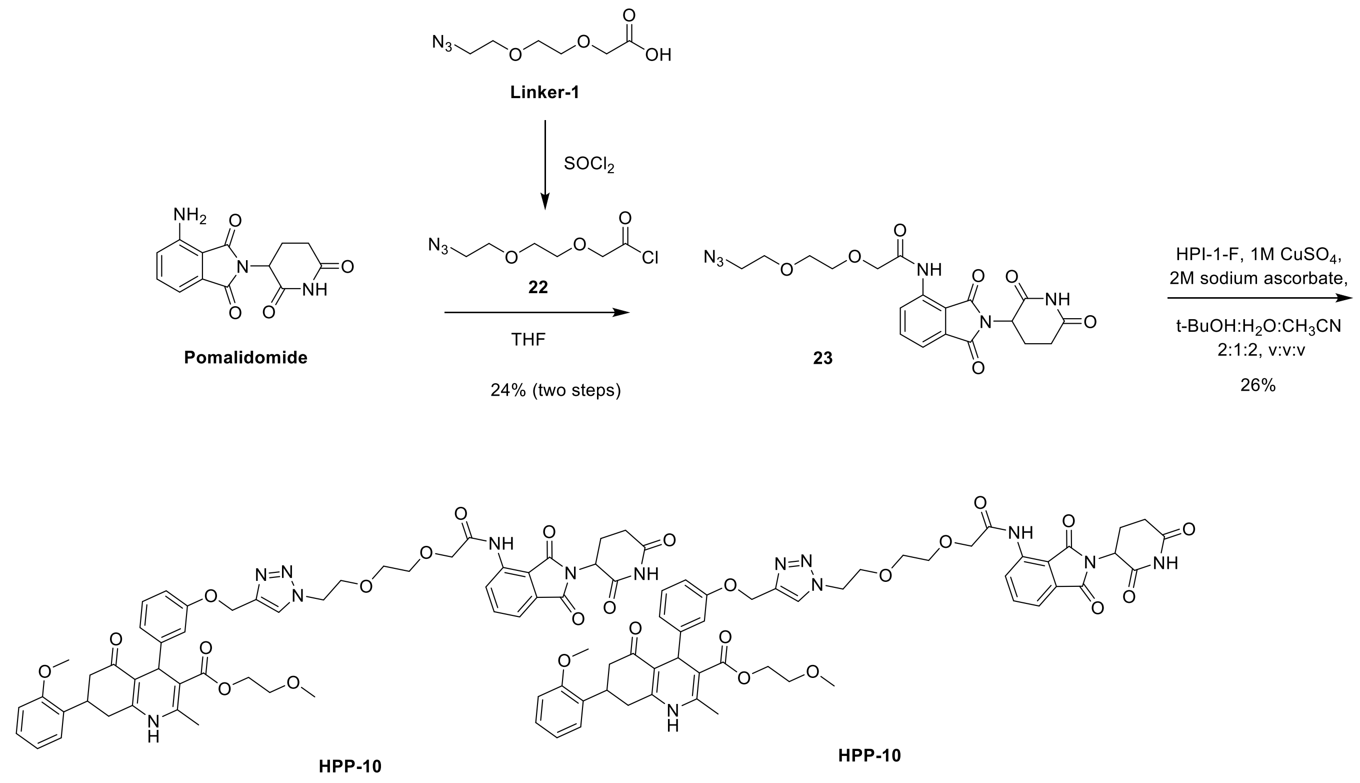


**Scheme S9.** Synthesis of **HPP-10**

**Compound 23.** To a solution of **linker-1** (100 mg, 0.53 mmol) was added $SOCl_2$ (77 µL, 1.06 mmol). The reaction mixture was refluxed for 3 h, after which it was checked for completion by TLC (diluted in MeOH,-pentane/EtOAc 1:1, $R_f$ = 0.53), then cooled to rt. The solution containing **22** was used directly for the next step without further purification. To a solution of **Pomalidomide** (66 mg, 0.24 mmol) in anhydrous THF (1.0 mL) was added the solution containing **22** (110 mg, theoretical mass, 0.53 mmol). The reaction mixture was refluxed for 4 h, after which it was concentrated and extracted with $CH_2Cl_2$. The organic layer was washed with sat. $NaHCO_3$ aq., dried over $MgSO_4$ and concentrated. The resulting crude was purified by reverse phase chromatography (BGB Scorpius C18 4.5 g, $H_2O$ + 0.1% TFA/$CH_3CN$ + 0.1% TFA 80:20 to 30:70) to give **23** as an off-white solid (50 mg, 24% two steps).; $R_f$= 0.24 ($CH_2Cl_2$/MeOH 20:1). $^1H$ NMR (400 MHz, $CDCl_3$): δ 10.45 (br, 1H, NH), 8.88 (d, *J* = 8.5 Hz, 1H), 7.95 (br, 1H, NH), 7.73 (dd, *J* = 8.4, 7.3 Hz, 1H), 7.58 (d, *J* = 7.3 Hz, 1H), 5.03 – 4.88 (m, 1H), 4.22 (s, 2H), 3.88 – 3.68 (m, 6H), 3.39 (t, *J* = 5.2 Hz, 2H), 2.99 – 2.66 (m, 3H), 2.23 – 2.11 (m, 1H). LC-MS ($ES^+$): m/z 444.89 $[M+H]^+$.

**HPP-10.** To a solution of compound **23** (13 mg, 0.029 mmol) and **HPI-1-F** (15 mg, 0.029 mmol) in degassed *t*-BuOH/$CH_3CN$/$H_2O$ (1.5 mL, 2:2:1) were added degassed aqueous solutions of 2 M sodium ascorbate (150 µL, 0.29 mmol) and 1 M $CuSO_4$ (15 µL, 0.015 mmol, 50 mol%) and the mixture was stirred overnight at rt. $CH_2Cl_2$ and $H_2O$ were added to the reaction mixture and after partitioning of the layers, the aqueous layer was extracted with $CH_2Cl_2$. The organic layer was dried over $MgSO_4$ and concentrated. The resulting crude was purified by reverse phase chromatography (BGB Scorpius C18 4.5 g, $H_2O$ + 0.1% TFA/$CH_3CN$ + 0.1% TFA 75:25 to 30:70) to give **HPP-10** as an off-white solid (5 mg, 26%).; $R_f$= 0.13 ($CH_2Cl_2$/MeOH 20:1). $^1$H NMR (500 MHz, $CDCl_3$): δ 10.42 (br, 1H, NH), 8.83 (d, *J* = 8.4 Hz, 1H), 8.75 (br, 1H, NH), 7.80 (s, 1H), 7.72 – 7.64 (m, 1H), 7.55 (d, *J* = 7.3 Hz, 1H), 7.26 – 6.62 (m, 8H), 6.34 (s, 1H), 5.13 (s, 3H), 4.96 (m, 1H), 4.62 – 4.43 (m, 2H), 4.15 (m, 4H), 3.96 (m, 2H), 3.82 – 3.71 (m, 7H), 3.64 – 3.55 (m, 1H), 3.52 (t, *J* = 4.9 Hz, 2H), 3.29 (s, 3H), 2.90 – 2.51 (m, 7H), 2.36 (s, 3H), 2.21 – 2.09 (m, 1H). $^{13}$C NMR (126 MHz, $CDCl_3$): δ 195.9, 171.2, 169.0, 168.5, 168.2, 168.1, 167.3, 166.7, 158.1, 157.2, 149.9, 148.8, 144.3, 144.1, 136.6, 136.4, 131.4, 130.5, 128.9, 128.0, 127.1, 125.2, 124.3, 121.4, 120.8, 118.9, 116.2, 114.4, 112.6, 110.7, 105.6, 71.7, 70.9, 70.5, 70.4, 69.9, 62.9, 61.6, 58.8, 55.2, 50.3, 49.3, 42.4, 36.5, 33.2, 32.9, 31.4, 22.7, 19.4. LC-MS (ES+): m/z 473.84 $[M+2H]^{2+}$ and m/z 946.14 $[M+H]^+$. ESI-HRMS (m/z): calcd. for $[C_{49}H_{51}N_7O_{13} + H]^+$ 946.3545; obsd. 946.3600.

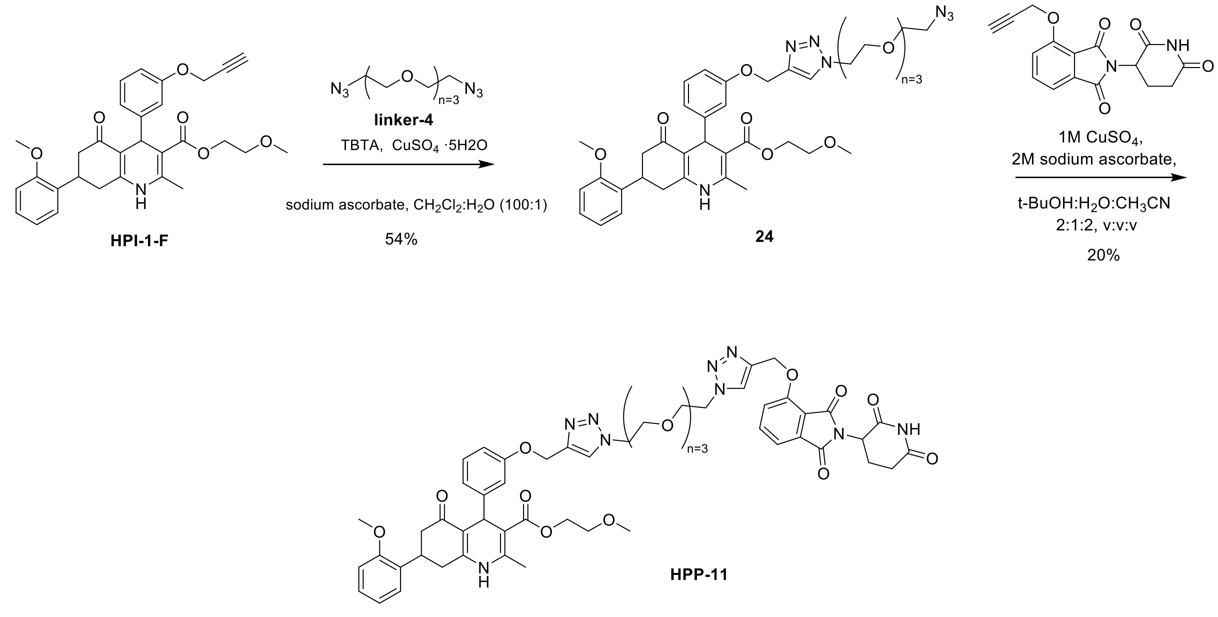


**Scheme S10.** Synthesis of **HPP-11**

**Compound 24.** To a solution of **HPI-1-F** (12 mg, 24 µmol), **linker-4** (60 mg, 245 µmol, 10eq) and TBTA (3.8 mg, 7 µmol) in $CH_2Cl_2$ (1.0 mL), a solution of $CuSO_4$ (3.8 mg, 24 µmol) and sodium ascorbate (4.8 mg, 24 µmol) in $H_2O$ (50 µL) was added. The mixture was vigorously stirred at rt and under nitrogen atmosphere for 2 h. The crude was purified by reverse phase chromatography (BGB Scorpius C18 4.5 g, $H_2O$ + 0.1% TFA/$CH_3CN$ + 0.1% TFA 95:5 to 10:90) to give **23 inactive** as a yellow oil (9.7 mg, 54%).

**HPP-11.** To a solution of compound **24** (14 mg, 0.019 mmol) and compound **10** (6 mg, 0.019 mmol) in degassed *t*-BuOH/$CH_3CN$/$H_2O$ (1.0 mL, 2:2:1) were added degassed aqueous solutions of 2 M sodium ascorbate (19 µL, 0.039 mmol) and 1 M $CuSO_4$ (4 µL, 0.004 mmol, 20 mol%) and the mixture was stirred overnight at rt. $CH_2Cl_2$ and $H_2O$ were added to the reaction mixture and after partitioning of the layers, the aqueous layer was extracted with $CH_2Cl_2$. The organic layer was dried over $MgSO_4$ and concentrated. The resulting crude was purified by PTLC ($SiO_2$, $CH_2Cl_2$/MeOH 15:1, eluted once) to give **HPP-11** as an off-white solid (4 mg, 20%).; $R_f$= 0.25 ($CH_2Cl_2$/MeOH 15:1). $^1$H NMR (500 MHz, $CDCl_3$): δ 8.44 (br, 1H, NH), 7.90 (s, 1H), 7.83 (s, 1H), 7.67 – 7.62 (m, 1H), 7.51 (dd, *J* = 8.4, 2.9 Hz, 1H), 7.45 (d, *J* = 7.2 Hz, 1H), 7.24 – 6.70 (m, 8H), 6.36 (br, 1H, NH), 5.41 (s, 2H), 5.18 – 5.07 (m, 3H), 4.92 (m, 1H), 4.54 – 4.45 (m, 4H), 4.16 (m, 2H), 3.86 (t, *J* = 5.1 Hz, 2H), 3.83 – 3.73 (m, 5H), 3.58 – 3.49 (m, 11H), 3.30 (s, 3H), 2.88 – 2.51 (m, 7H), 2.35 (s, 3H), 2.10 (m, 1H). $^{13}$C NMR (126 MHz, $CDCl_3$): δ 195.6, 171.0, 168.2, 168.2, 167.3, 166.9, 165.7, 158.2, 157.1, 155.7, 149.6, 148.8, 144.2, 142.8, 136.7, 133.7, 130.5, 128.9, 128.0, 127.1, 124.7, 124.2, 121.3, 120.7, 120.1, 116.5, 114.5, 112.3, 110.7, 105.5, 70.5, 70.5, 70.3, 69.3, 69.2, 63.2, 62.9, 61.7, 58.8, 55.2, 50.3, 50.2, 49.1, 42.5, 36.4, 33.2, 32.9, 31.4, 22.6, 19.4. LC-MS ($ES^+$): m/z 529.67 $[M+2H]^{2+}$ and m/z 1058.25 $[M+H]^+$. ESI-HRMS (m/z): calcd. for $[C_{54}H_{59}N_9O_{14} + H]^+$ 1058.4181; obsd. 1058.4255.

### 2.NMR Spectra

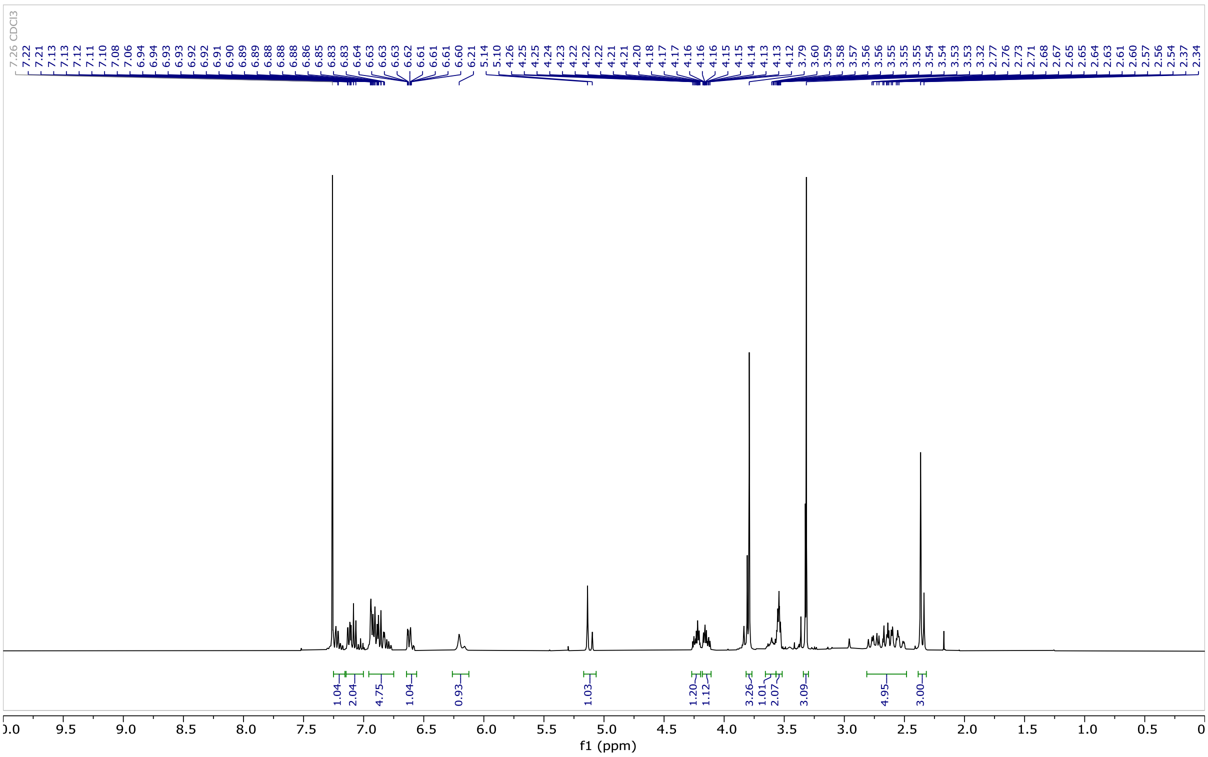


**Figure S1**. $^{1}H$ NMR (400 MHz, $CDCl_3$) of **HPI-1**

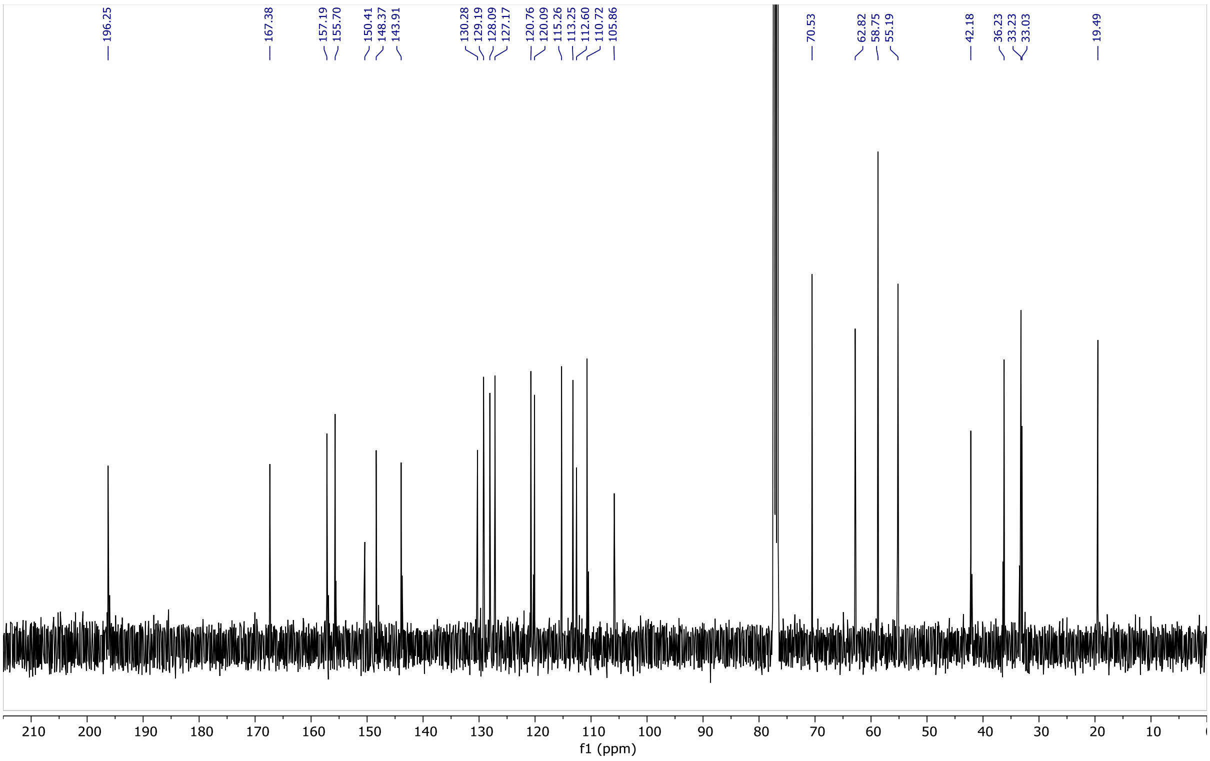


**Figure S2**. $^{13}C$ NMR (101 MHz, $CDCl_3$) of **HPI-1**

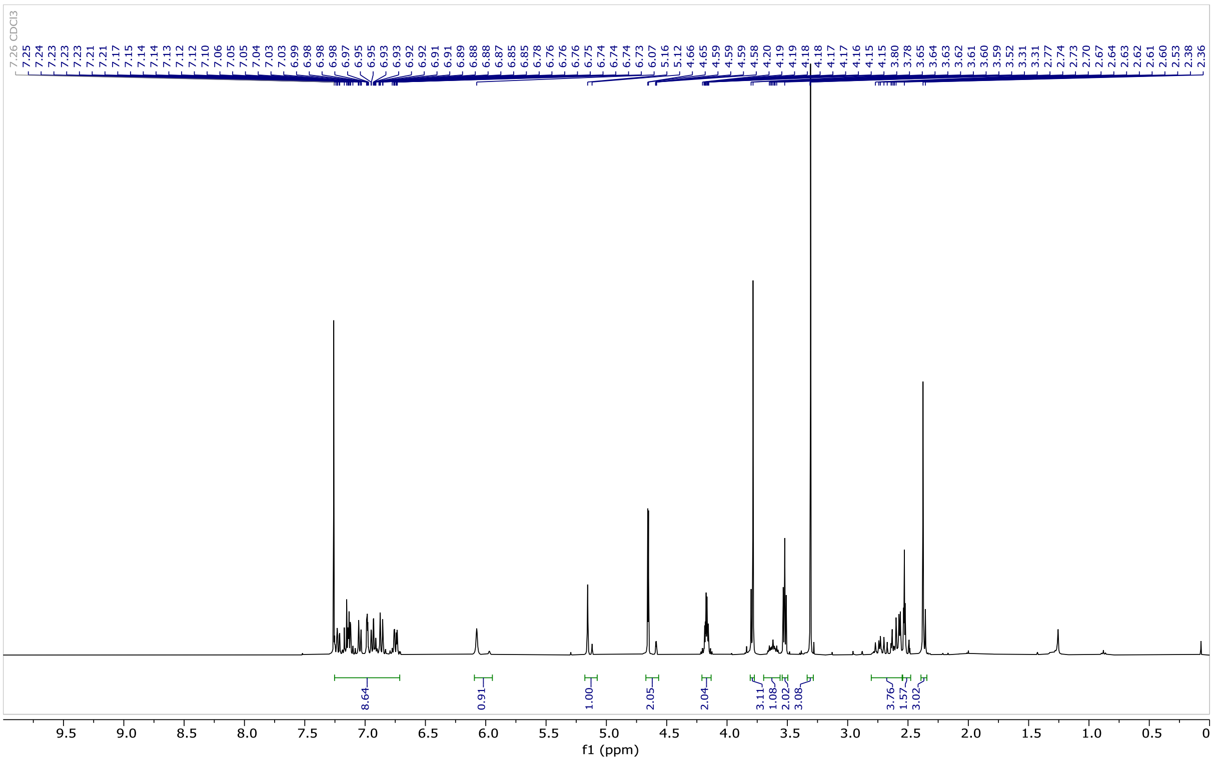

**Figure S3**. $^{1}$H NMR (400 MHz, $CDCl_3$) of **HPI-1-F**

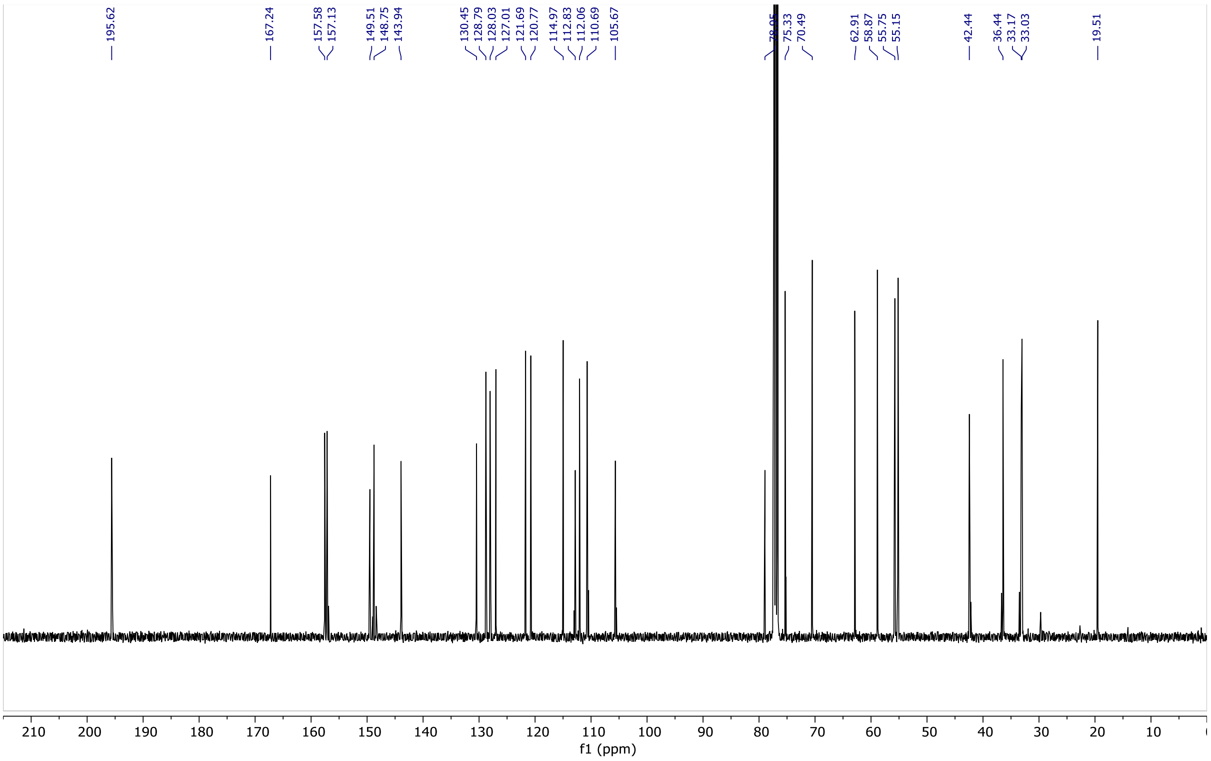

**Figure S4**. $^{13}$C NMR (101 MHz, $CDCl_3$) of **HPI-1-F**

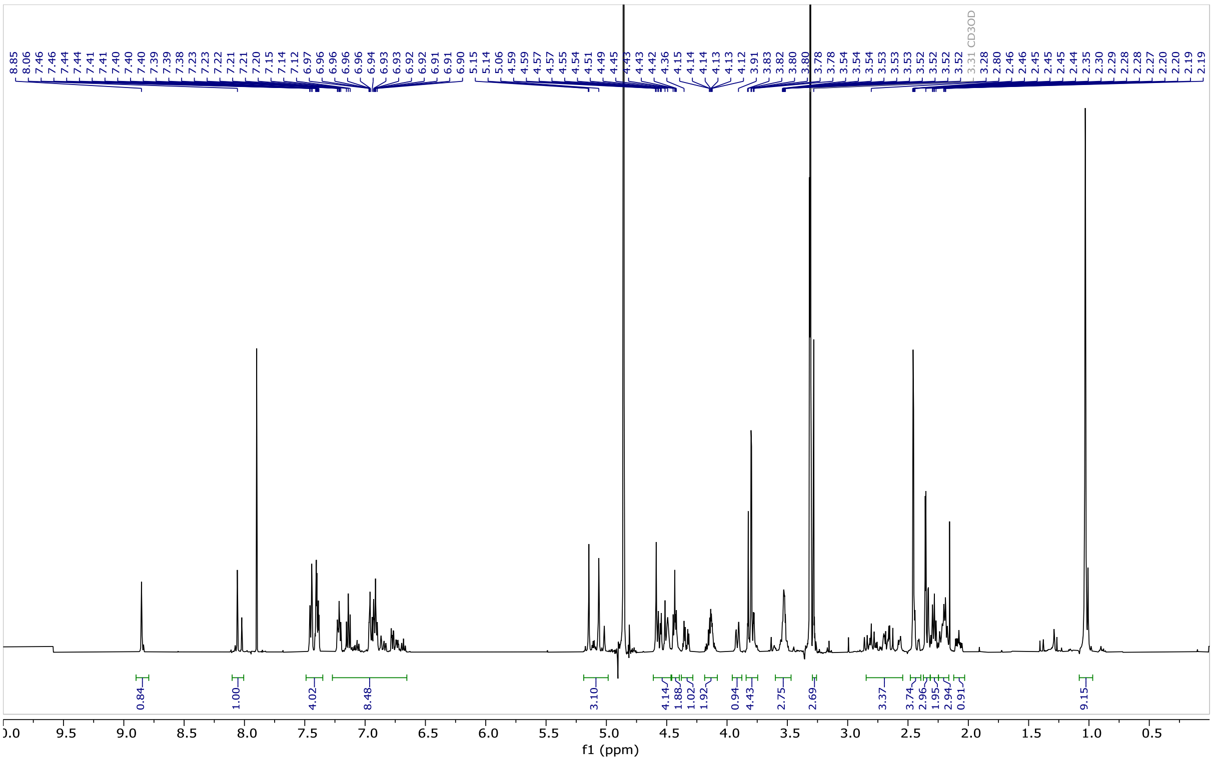

**Figure S5**. $^{1}H$ NMR (500 MHz, $CD_3OD$) of **HPP-1**

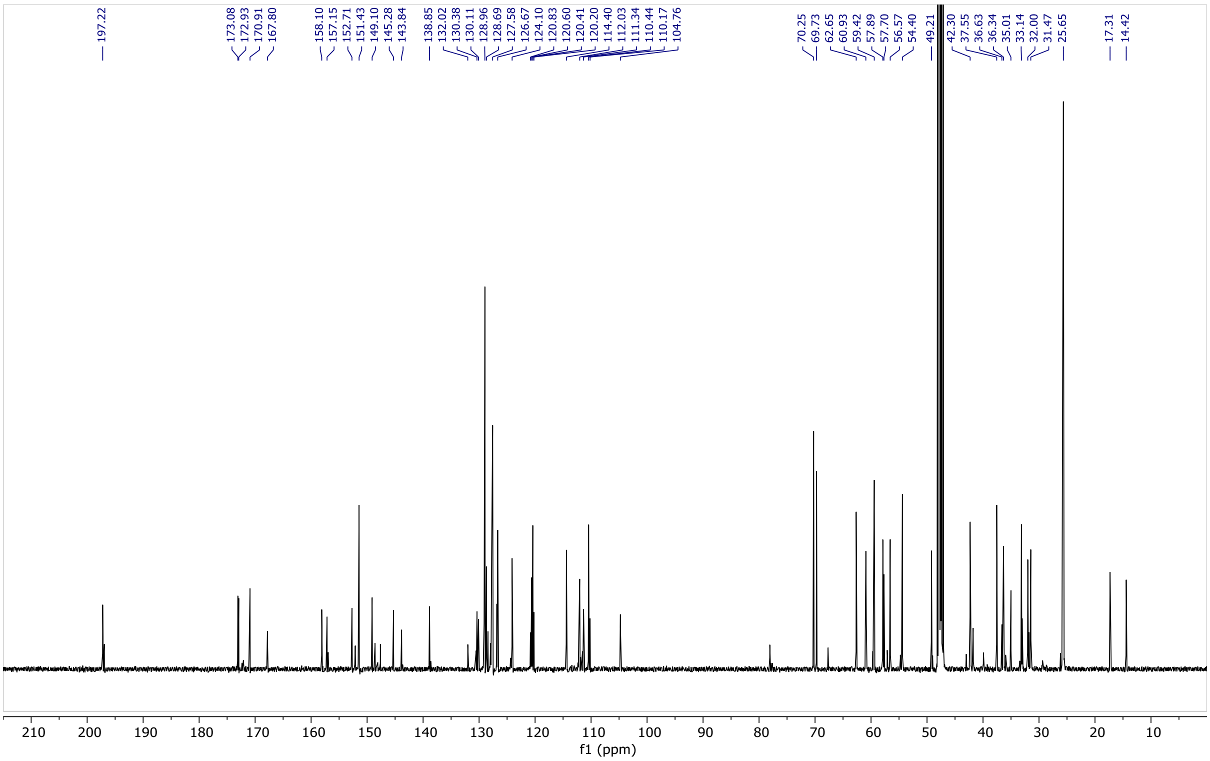

**Figure S6**. $^{13}C$ NMR (126 MHz, $CD_3OD$) of **HPP-1**

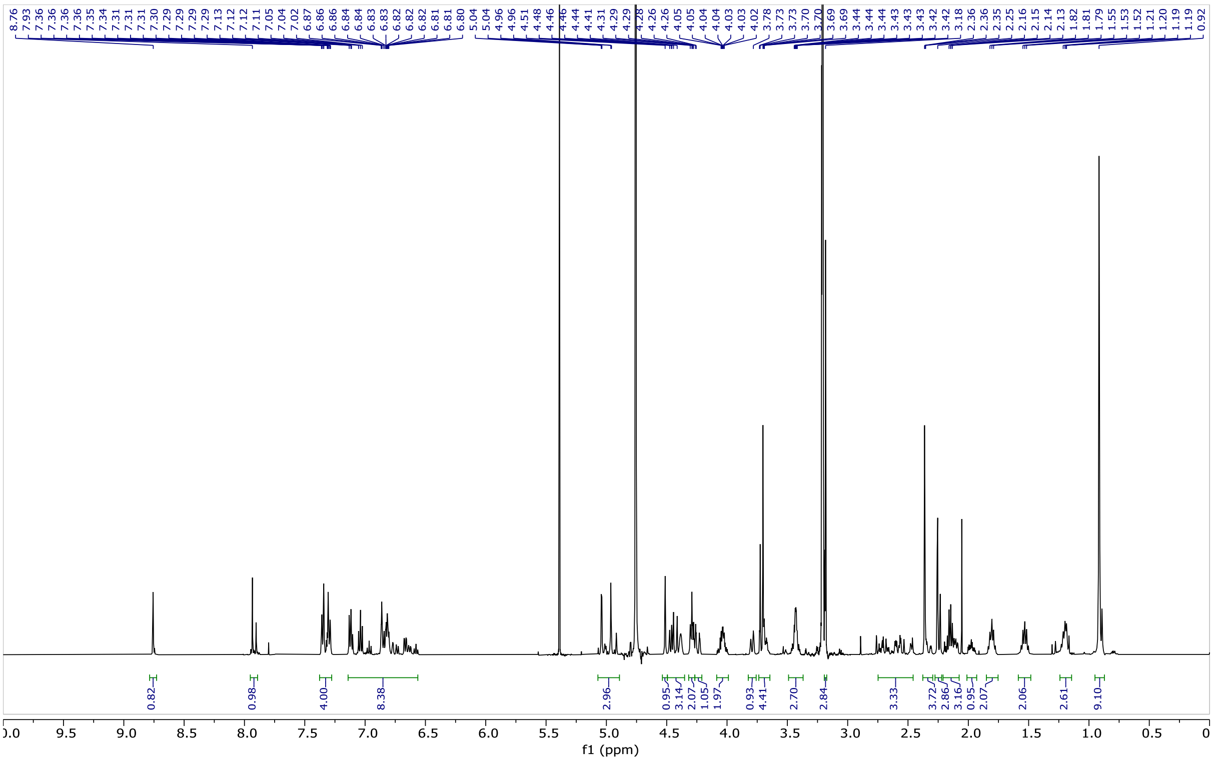

**Figure S7**. $^{1}H$ NMR (500 MHz, $CD_3OD$) of **HPP-2**

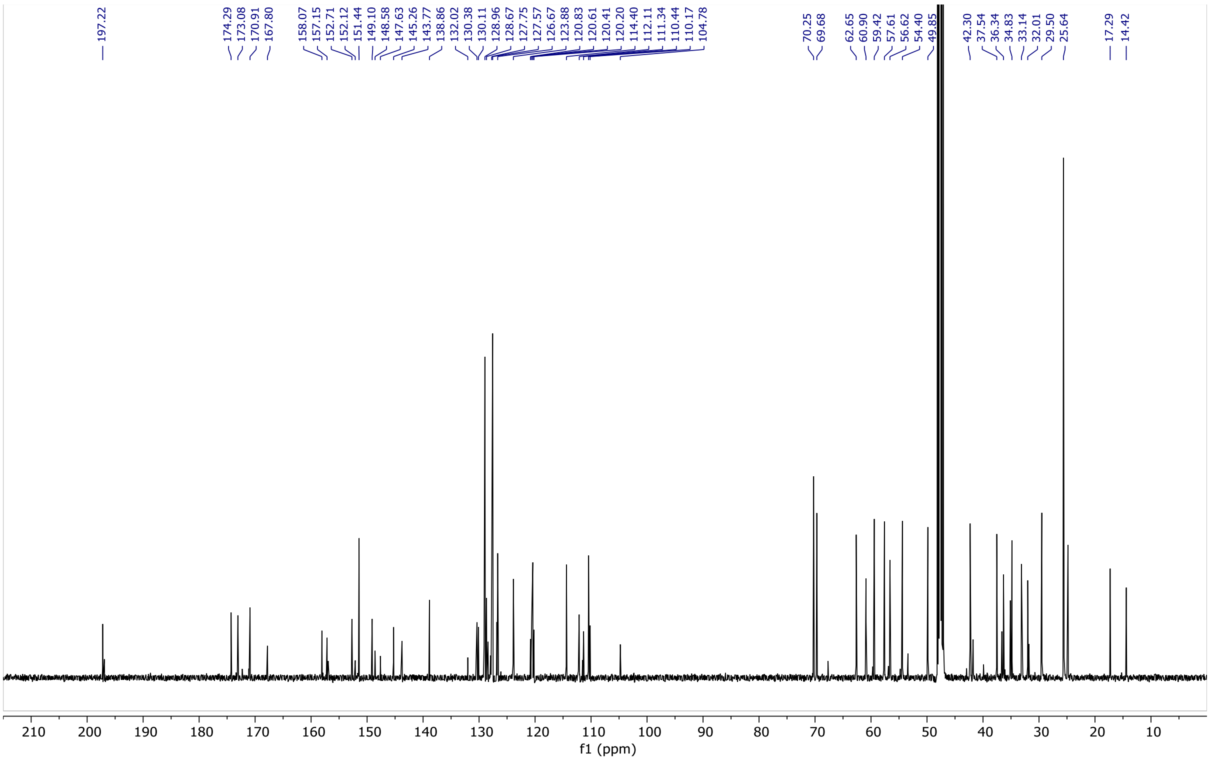

**Figure S8**. $^{13}C$ NMR (126 MHz, $CD_3OD$) of compound **HPP-2**

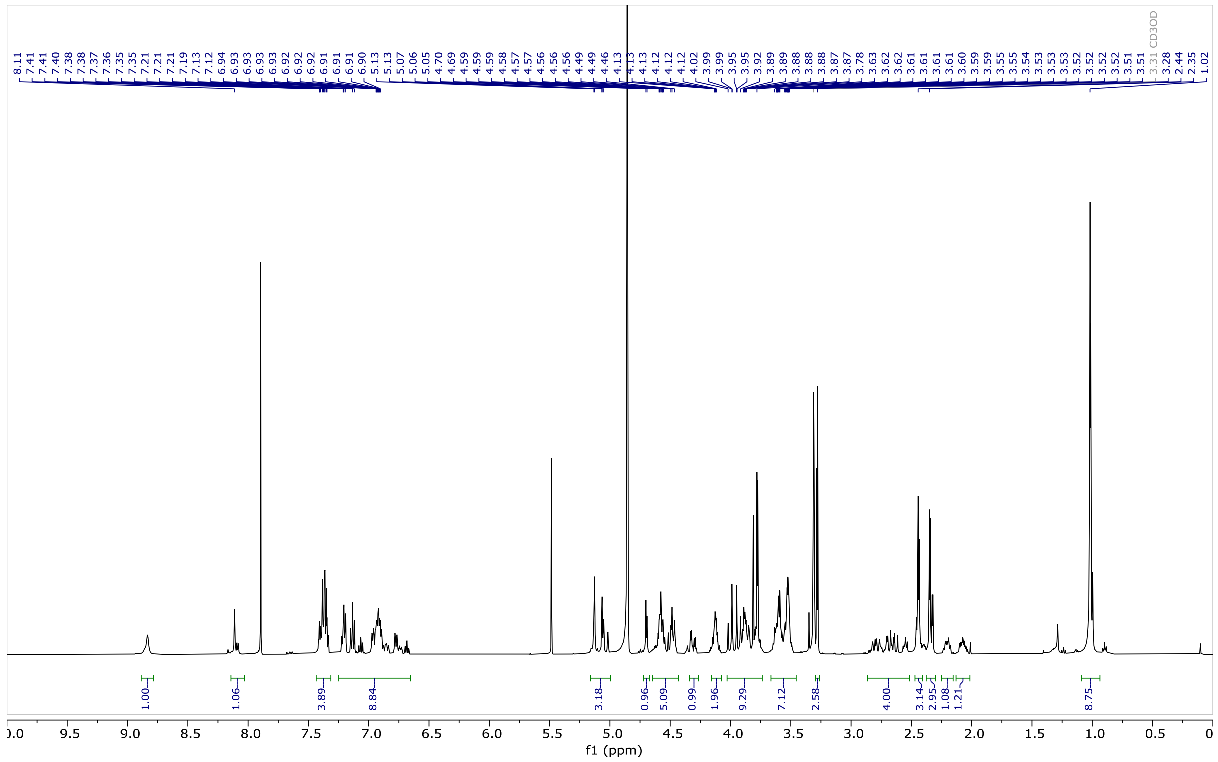

**Figure S9**. $^{1}H$ NMR (500 MHz, $CD_3OD$) of **HPP-3**

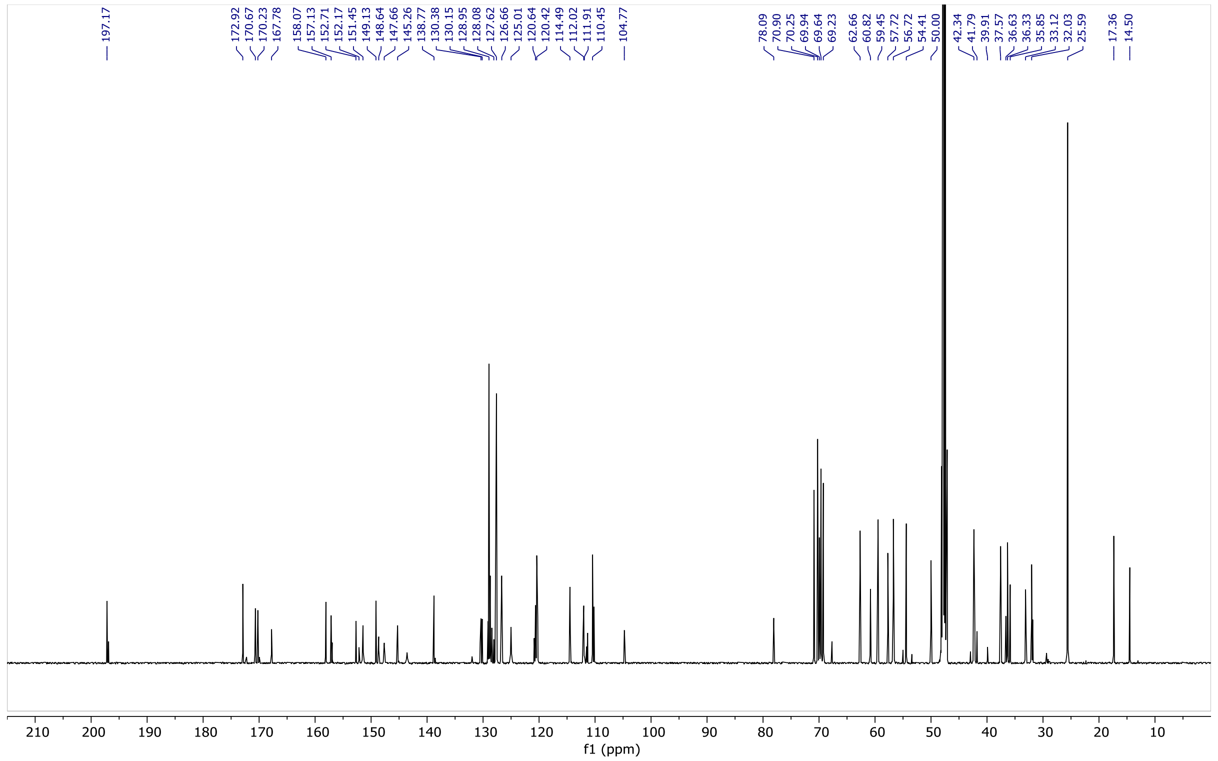

**Figure S10**. $^{13}C$ NMR (126 MHz, $CD_3OD$) of **HPP-3**

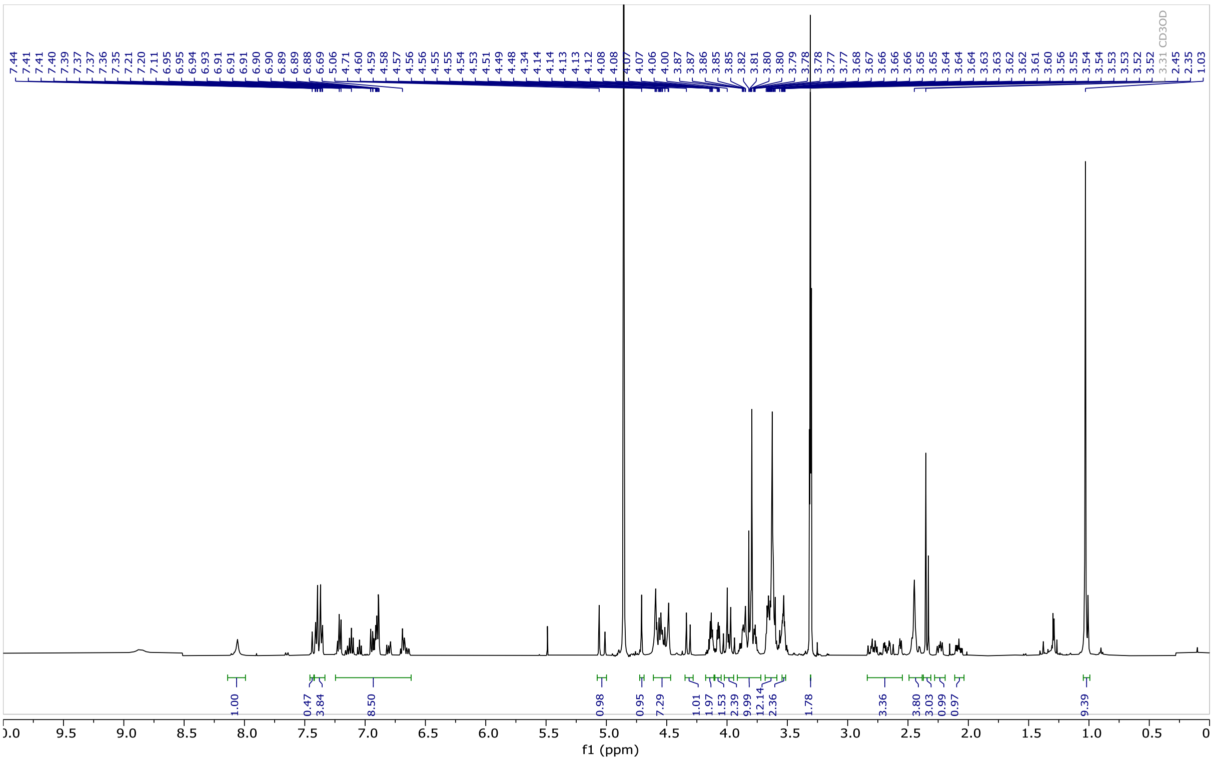


**Figure S11**. $^{1}H$ NMR (400 MHz, $CD_3OD$) of **HPP-4**

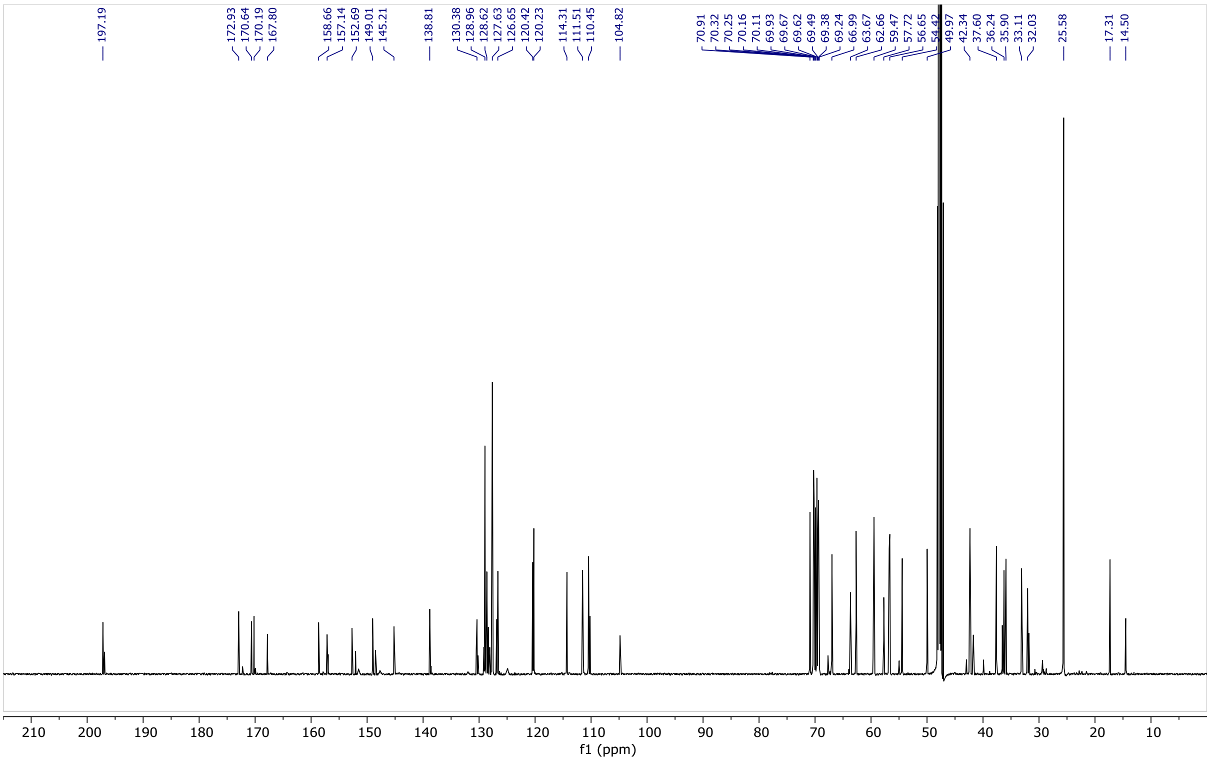


**Figure S12**. $^{13}C$ NMR (126 MHz, $CD_3OD$) of compound **HPP-4**

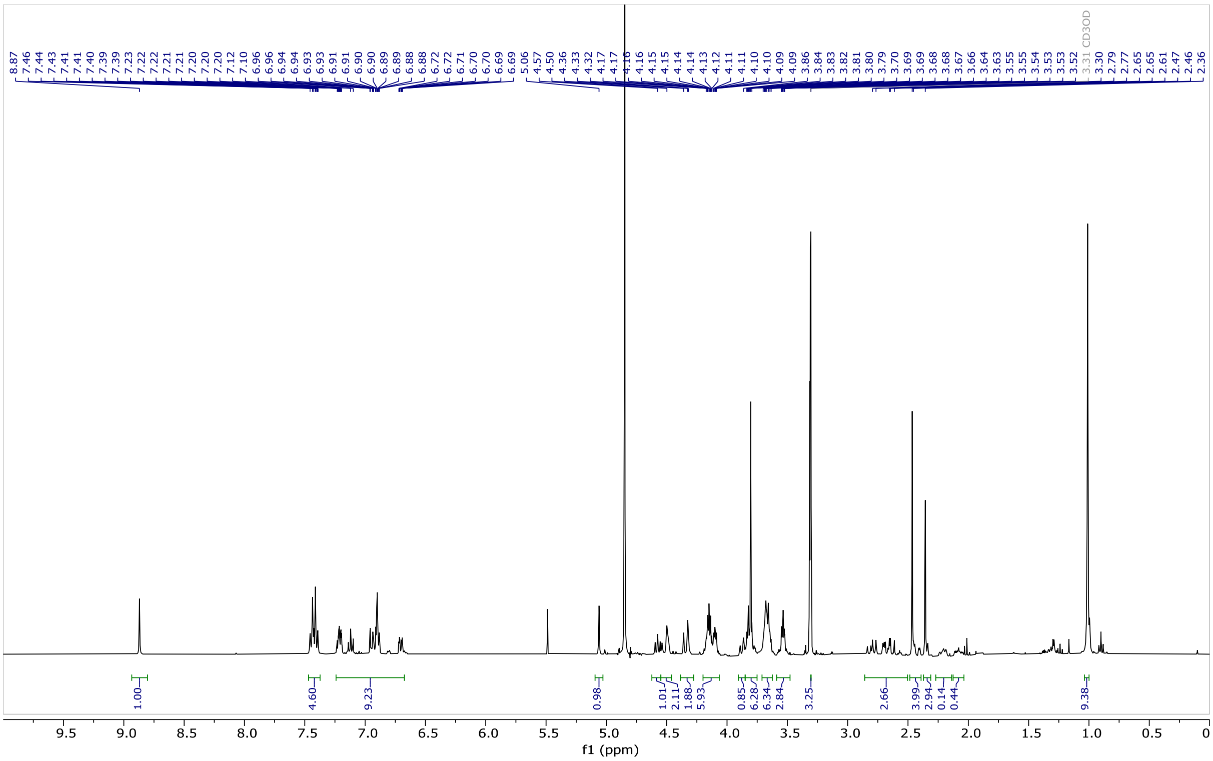


**Figure S13**. $^{1}H$ NMR (500 MHz, $CD_3OD$) of **HPP-5**

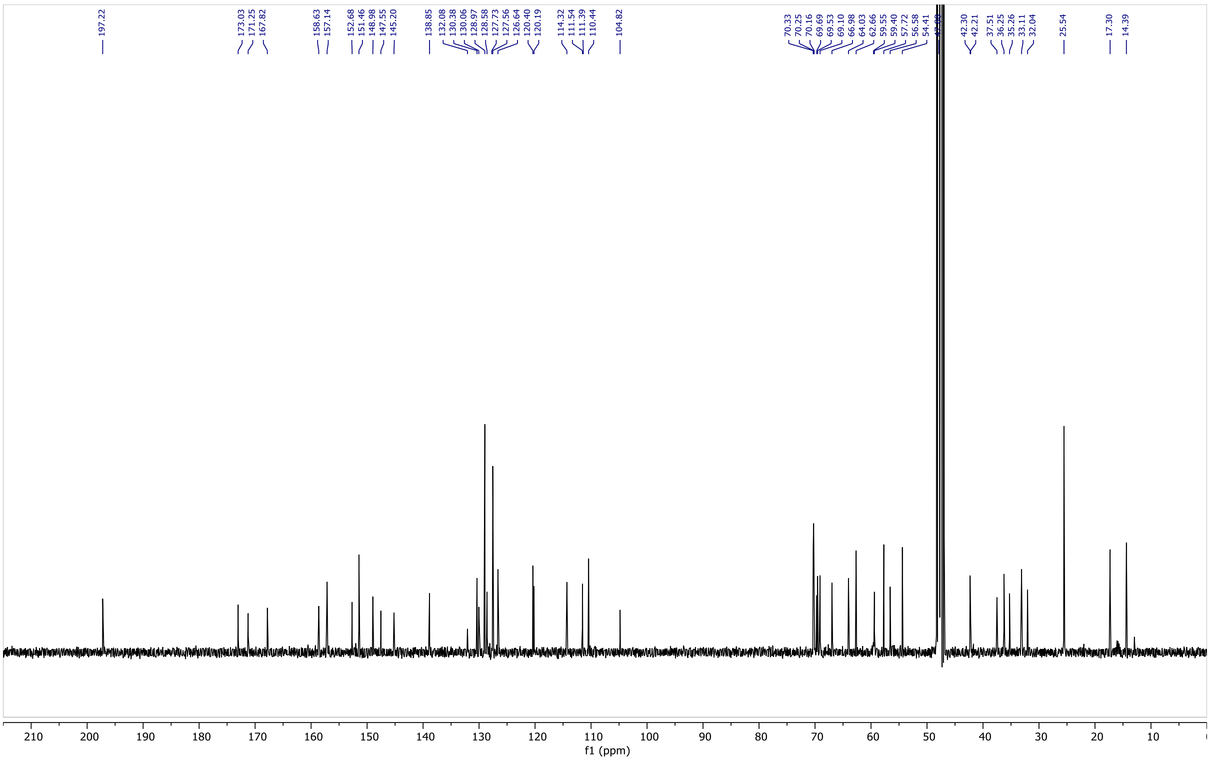


**Figure S14**. $^{13}C$ NMR (126 MHz, $CD_3OD$) of **HPP-5**

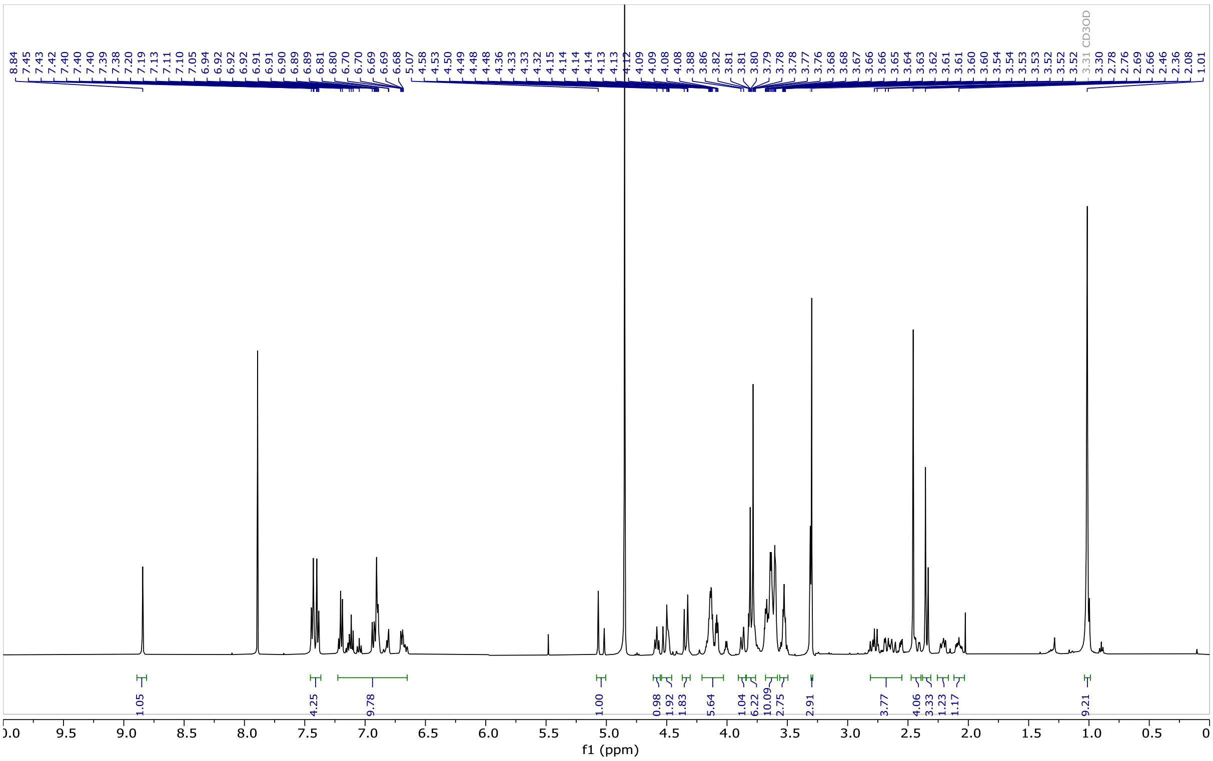

**Figure S15**. $^{1}H$ NMR (500 MHz, $CD_3OD$) of **HPP-6**

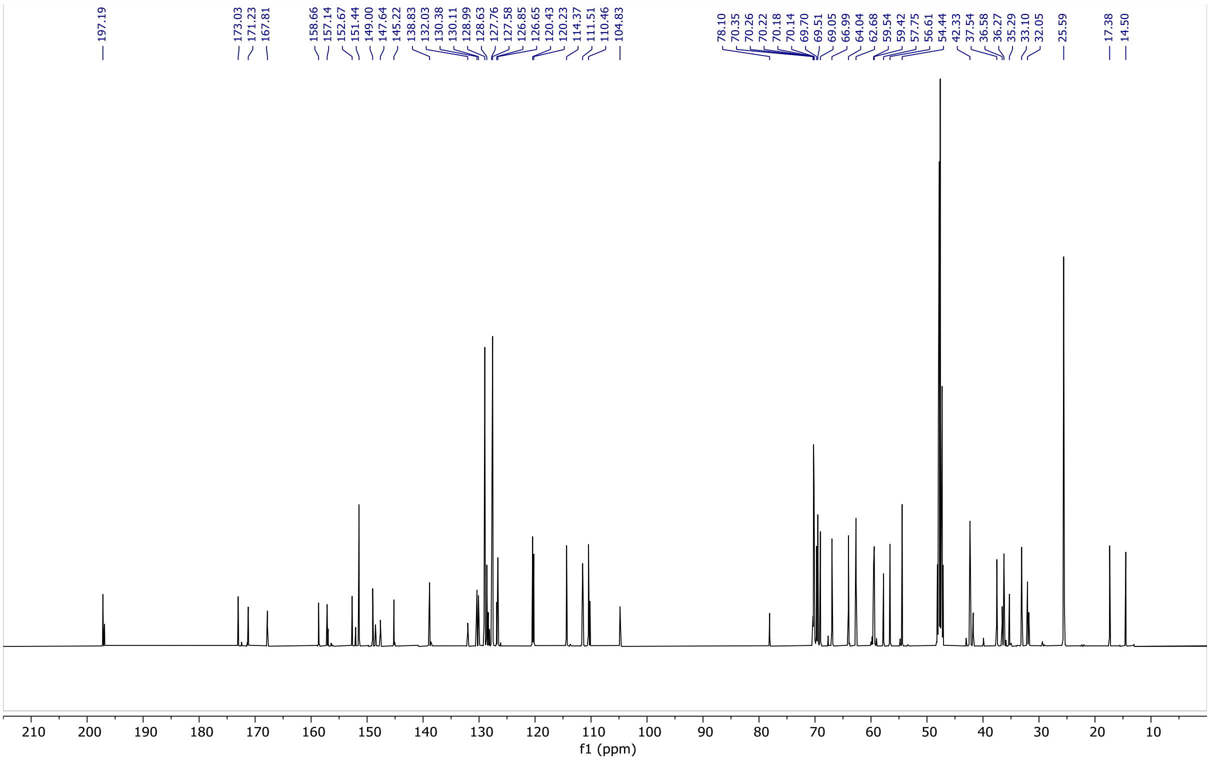

**Figure S16**. $^{13}C$ NMR (126 MHz, $CD_3OD$) of **HPP-6**

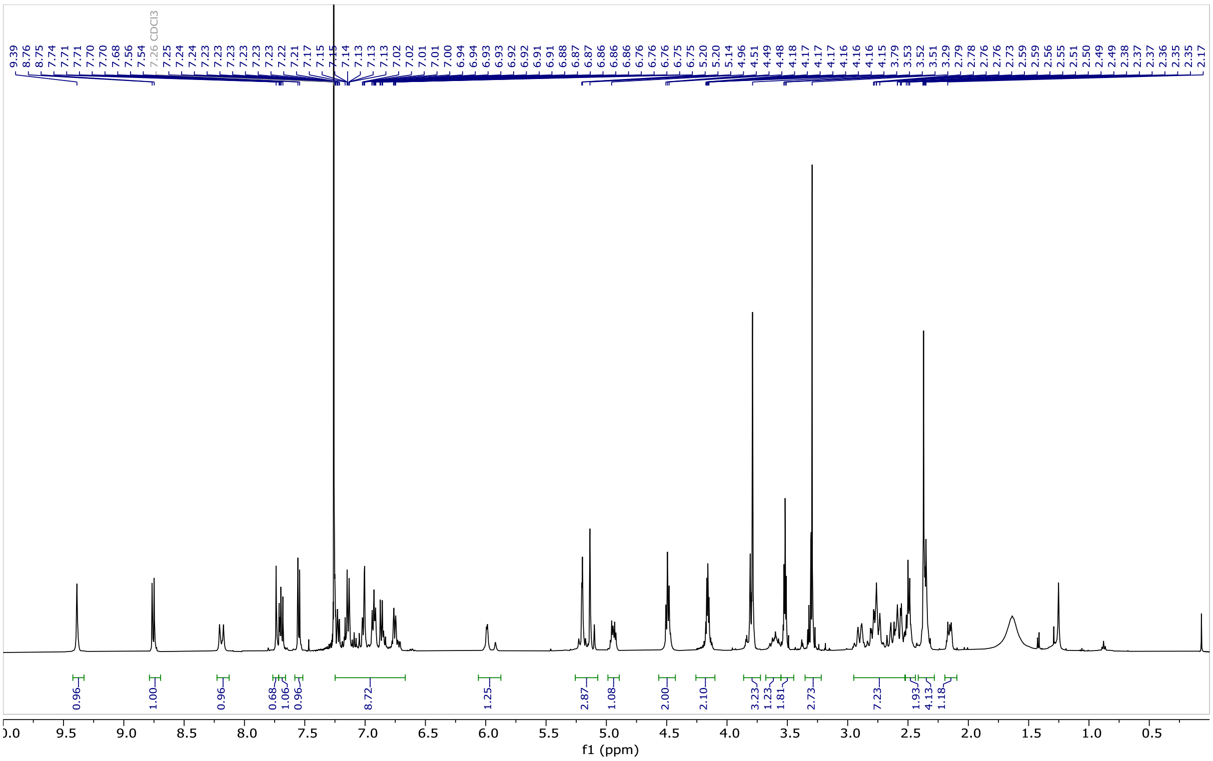


**Figure S17**. $^{1}H$ NMR (500 MHz, $CDCl_3$) of **HPP-7**

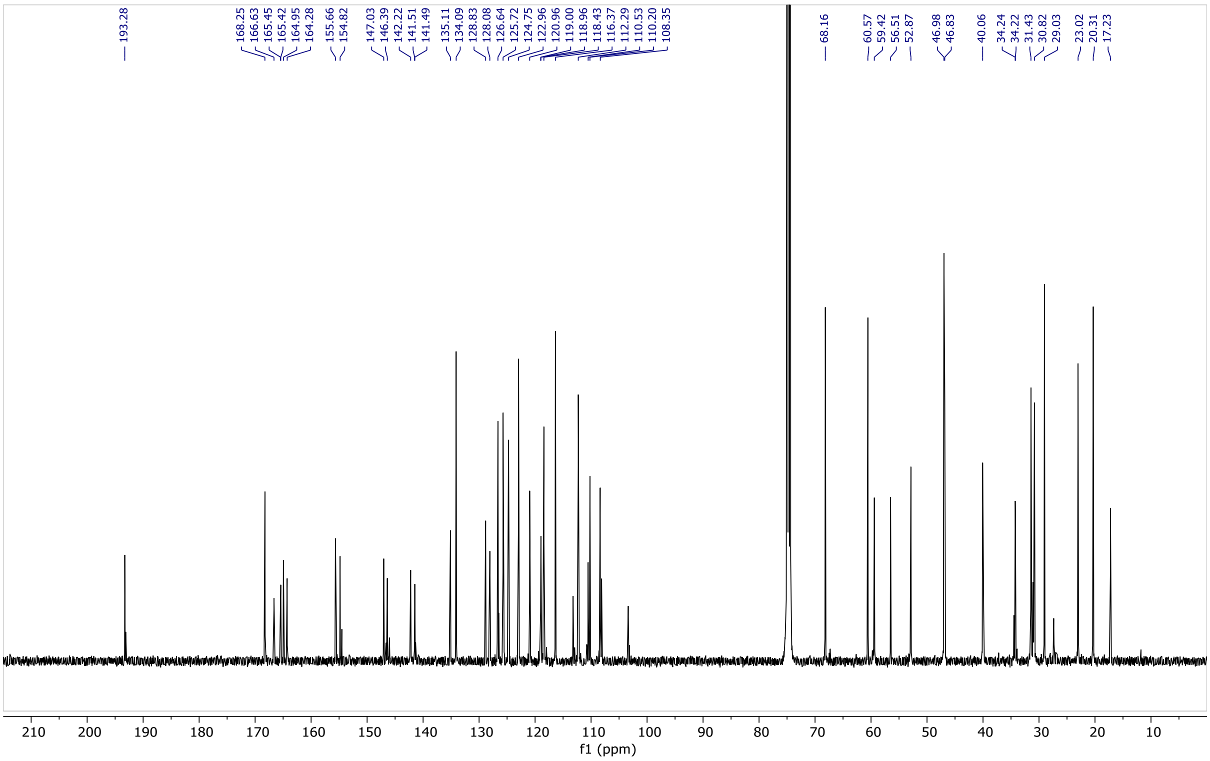


**Figure S18**. $^{13}C$ NMR (126 MHz, $CDCl_3$) of **HPP-7**

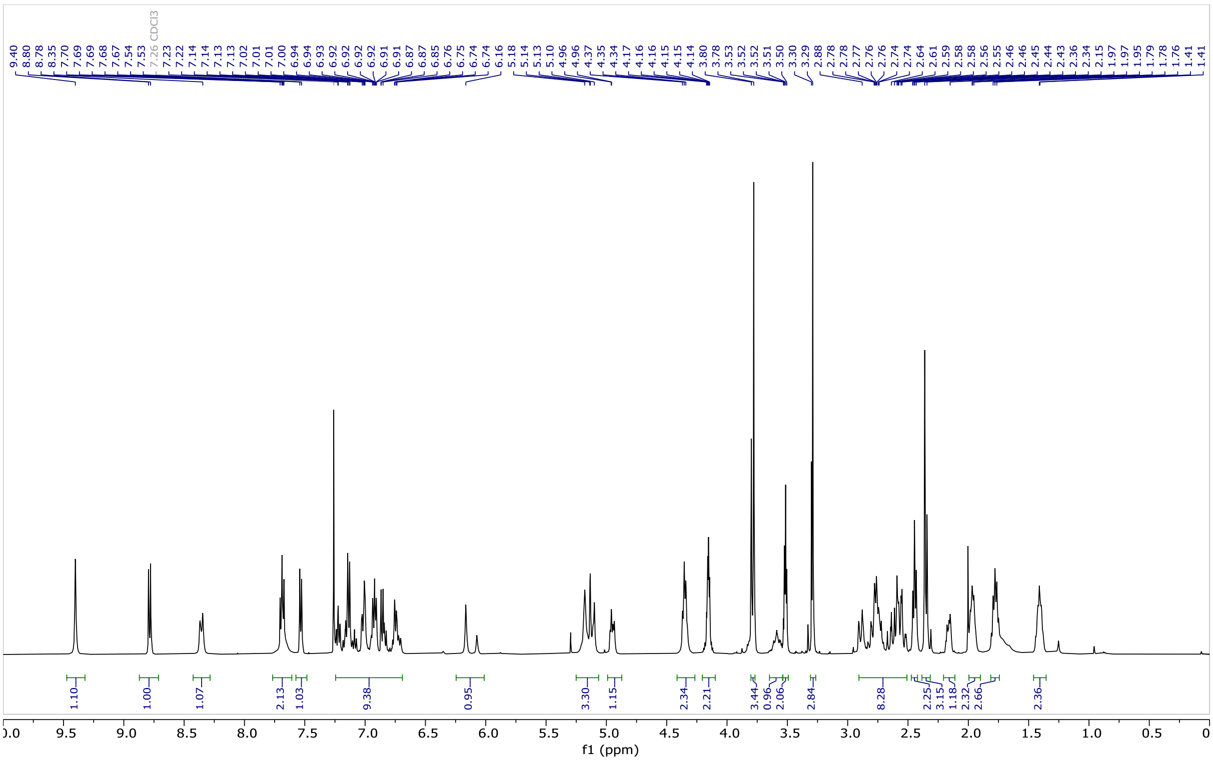

**Figure S19**. $^{1}H$ NMR (500 MHz, $CDCl_3$) of **HPP-8**

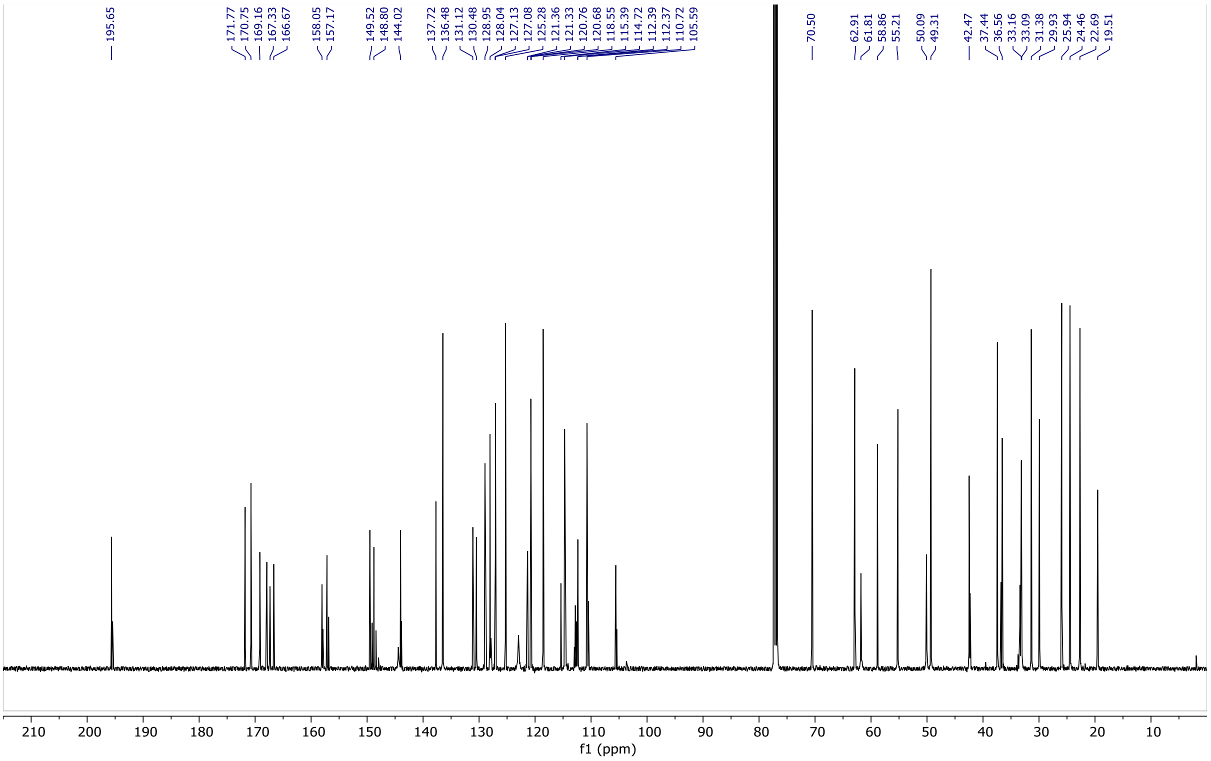

**Figure S20**. $^{13}C$ NMR (126 MHz, $CDCl_3$) of **HPP-8**

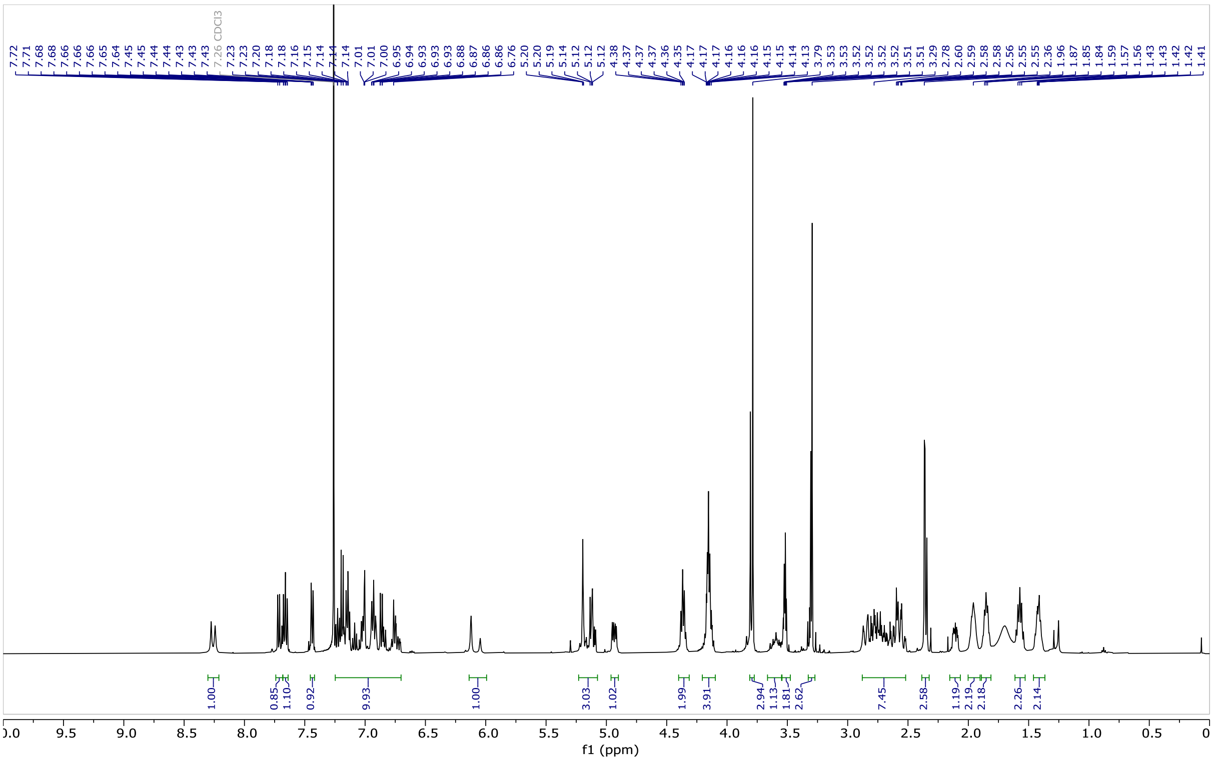

**Figure S21**. $^{1}H$ NMR (500 MHz, $CDCl_3$) of **HPP-9**

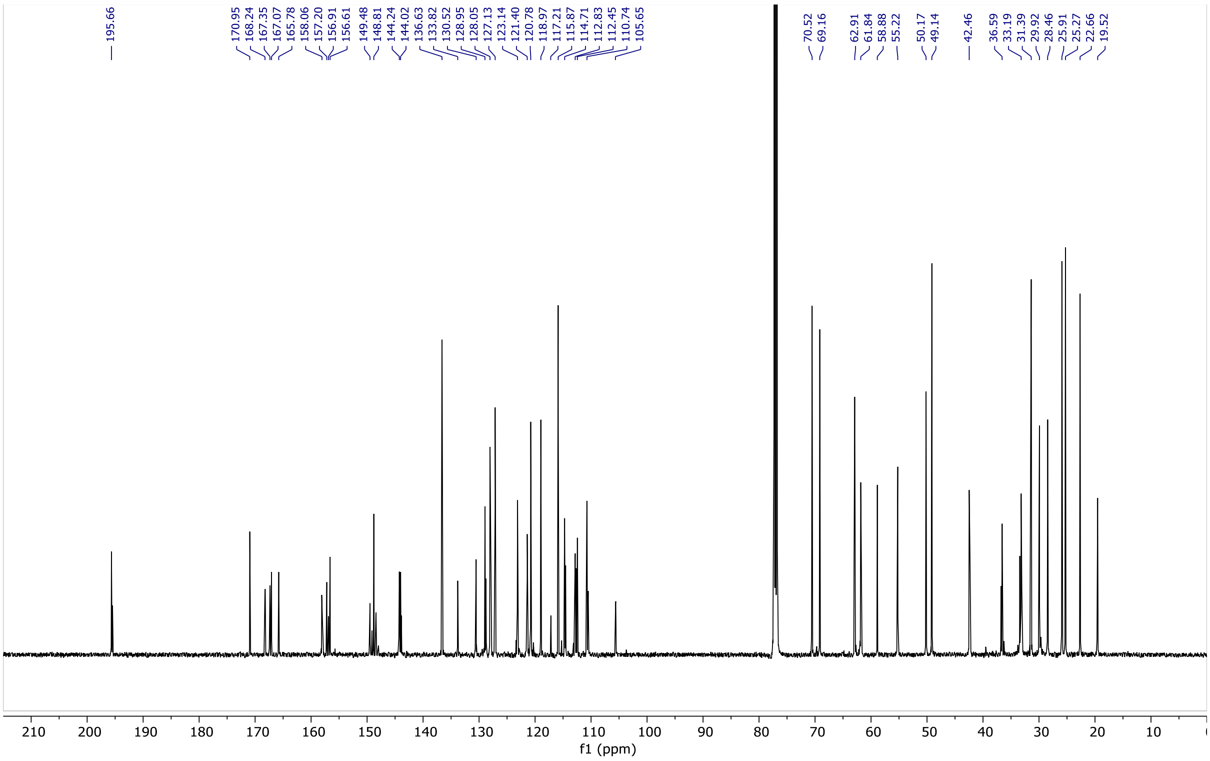

**Figure S22**. $^{13}C$ NMR (126 MHz, $CDCl_3$) of **HPP-9**

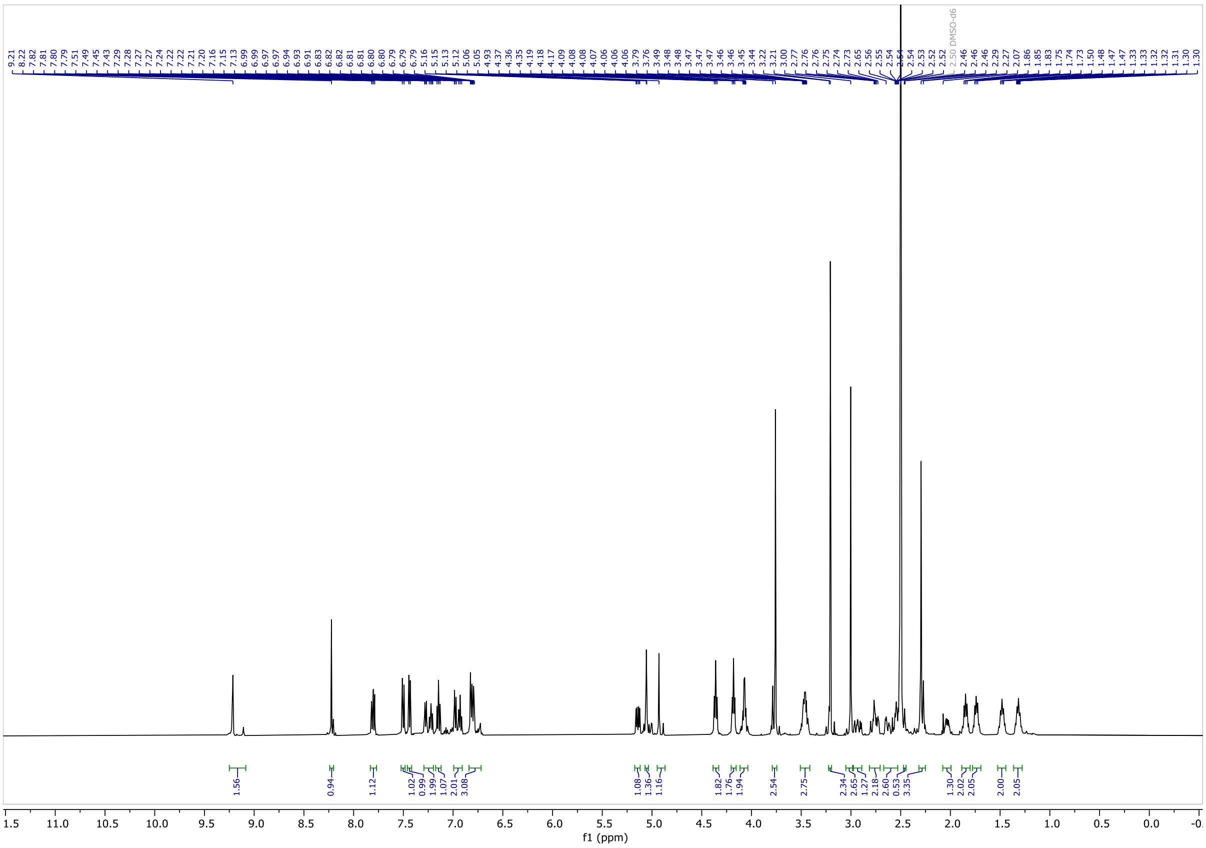

**Figure S23**. $^{13}$C NMR (126 MHz, DMSO-d$_6$) of **HPP-9-inact.**

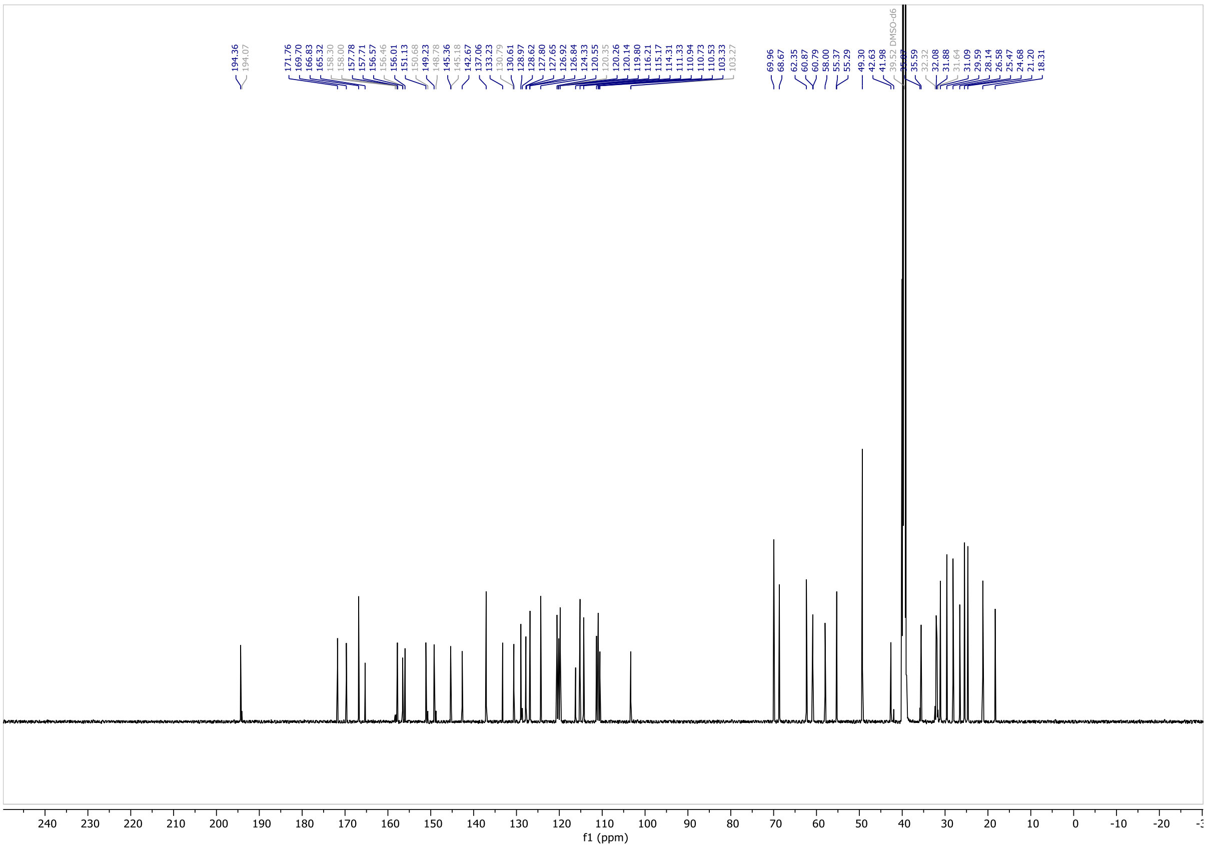

**Figure S24**. $^{13}$C NMR (126 MHz, DMSO-d$_6$) of **HPP-9-inact**

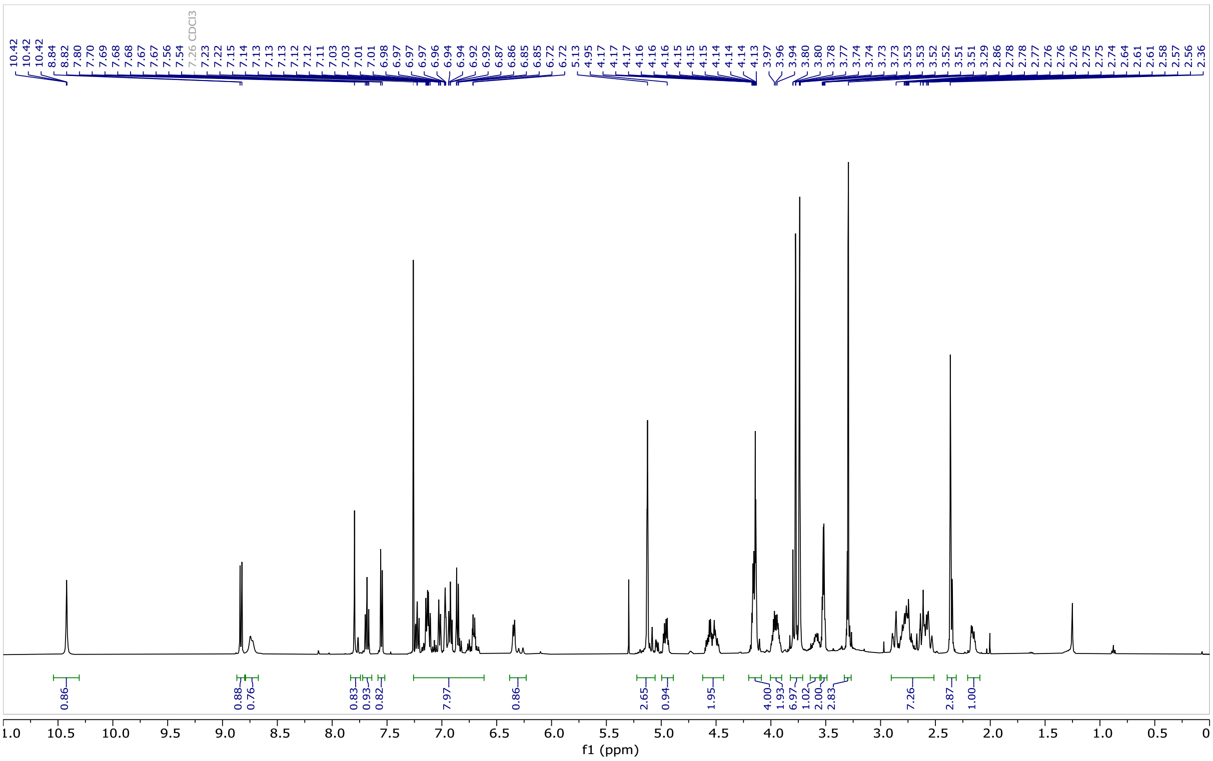

**Figure S25**. $^{1}$H NMR (500 MHz, CDCl$_3$) of **HPP-10**

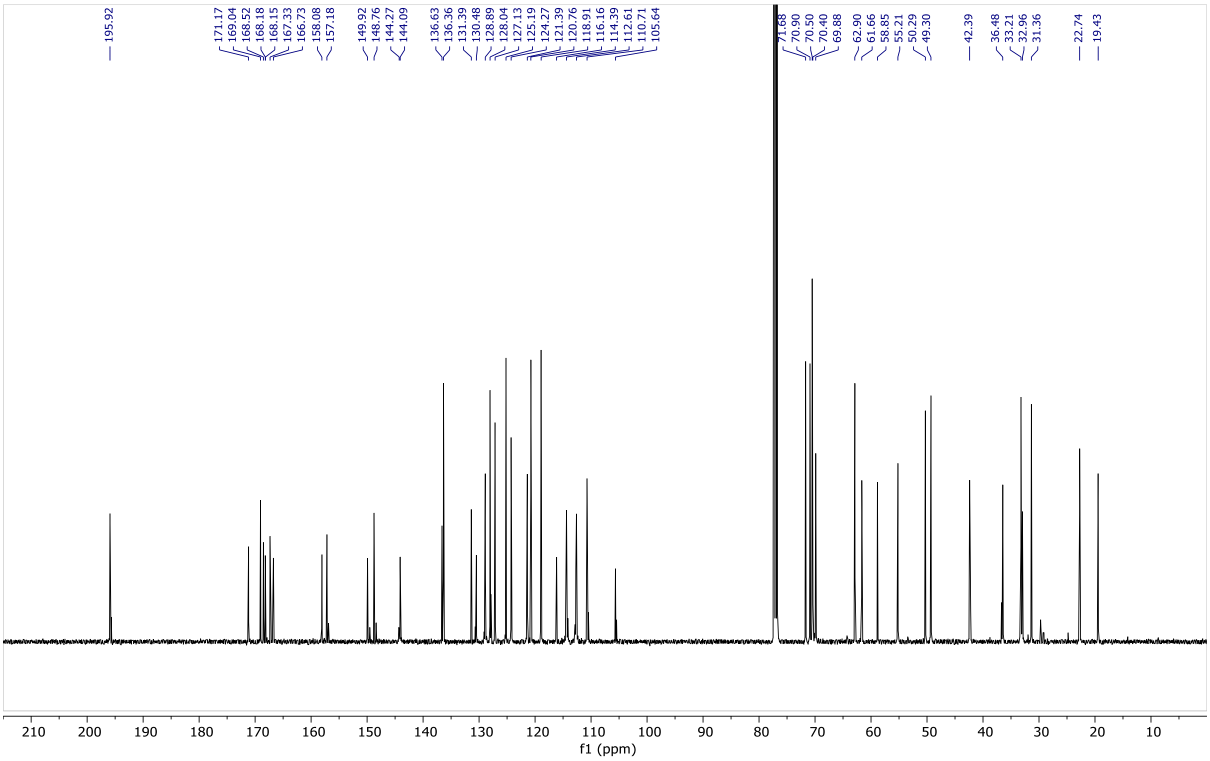

**Figure S26**. $^{13}C$ NMR (126 MHz, $CDCl_3$) of **HPP-10**

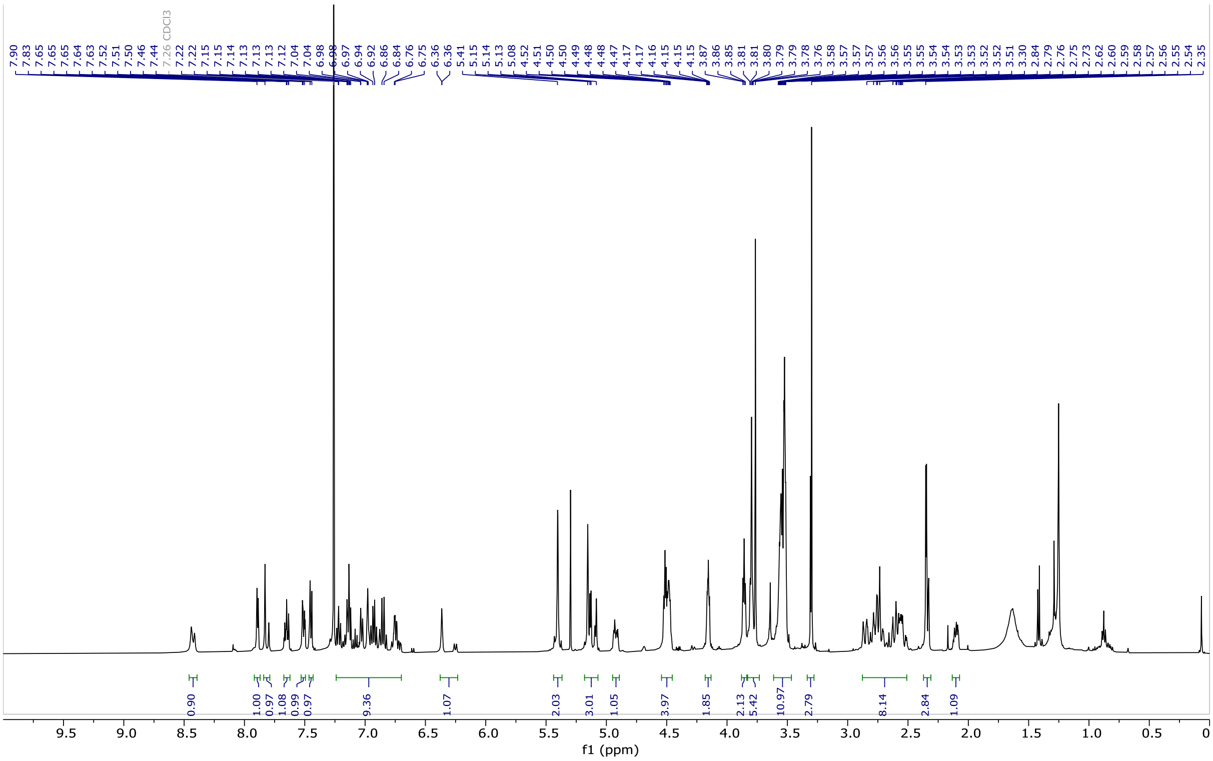

**Figure S27**. $^{1}H$ NMR (500 MHz, $CDCl_3$) of **HPP-11**

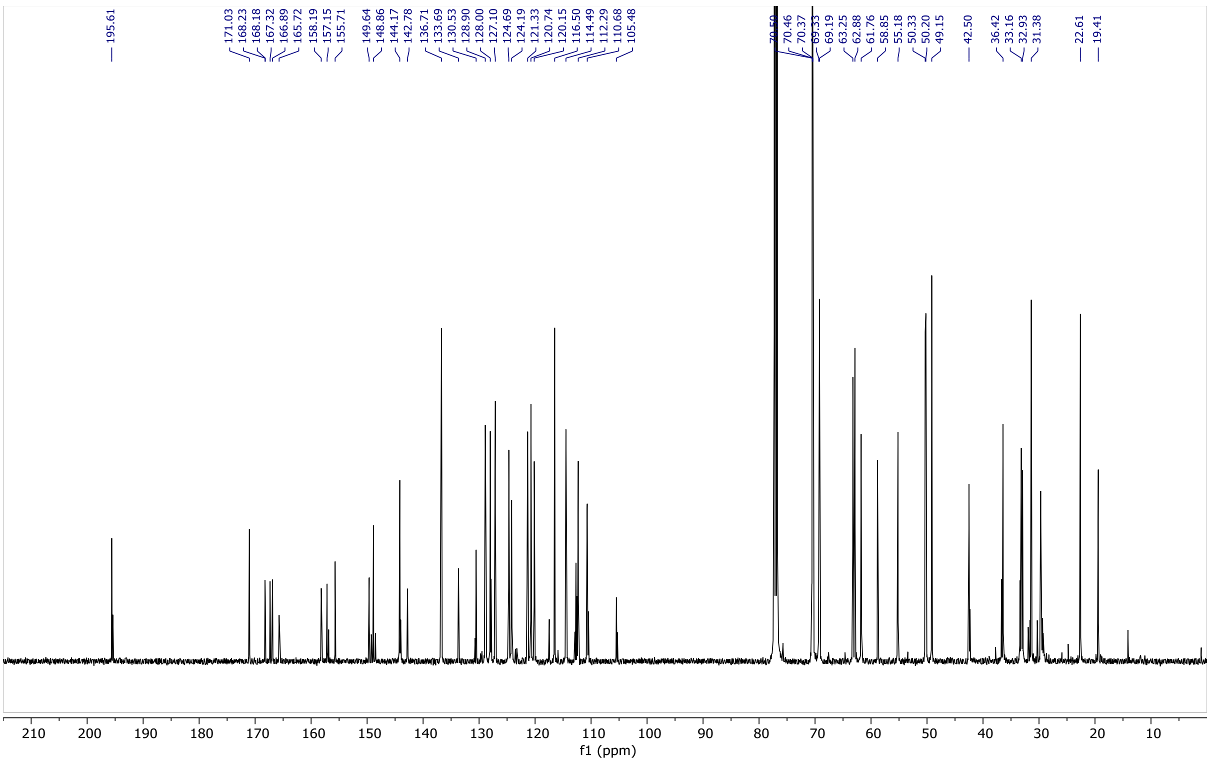


**Figure S28**. $^{13}$C NMR (126 MHz, $CDCl_3$) of **HPP-11**
